# Serotonergic Polypharmacology of 2-Halogenated Tryptamines

**DOI:** 10.64898/2026.04.16.718915

**Authors:** Jeanine Yacoub, Elena Bray, Jude Bayyat, Grant C. Glatfelter, Alex Leake, Emilia M. Buitrago, Hubert Nguyen, Alexander D. Maitland, John Partilla, John L. McKee, Natalie G. Cavalco, Serena S. Schalk, Josie C. Lammers, Michael H. Baumann, John D. McCorvy, James W. Leahy, Danielle Gulick, Christopher G. Witowski, Jacqueline L. von Salm

## Abstract

Serotonergic psychedelics such as *N,N*-dimethyltryptamine (DMT) and 4-phosphoryloxy-*N,N*-dimethyltryptamine (psilocybin) show therapeutic promise for psychiatric and neurodegenerative disorders but may be limited by liabilities from serotonin (5-HT)-2A mediated psychoactive effects and potential cardiotoxicity via 5-HT_2B_ activation. To address these limitations, we designed and synthesized 2-halogenated derivatives of DMT and psilacetin to reduce 5-HT_2A_/5-HT_2B_ activity while retaining engagement of therapeutically relevant targets, particularly 5-HT_6_, 5-HT_2C_, and 5-HT_1B_. This study demonstrated that 2-position halogenation decreased affinities, potencies, and efficacies at 5-HT_2A_ and 5-HT_1A_ receptors while preserving potent 5-HT_6_ agonism, especially for 2-Br-psilocin. The analogues exhibited reduced affinities at 5-HT_2B_ and hERG ion channels, suggesting safer cardiac valve and cardiotoxic profiles. In C57BL/6J mice, 2-Br-psilacetin did not induce the head-twitch response and attenuated 2,5 dimethoxy-4-iodoamphetamine (DOI)-induced head-twitch behavior, suggesting a reduced potential for inducing psychedelic effects. Behavioral assays further revealed improvements in stress-induced affective measures and hippocampus-independent cued learning at intermediate doses. These findings identify 2-halogenated tryptamines as polypharmacological serotonergic ligands with reduced psychoactivity and cardiac valve and toxic liabilities, supporting their potential as next-generation psychedelic-inspired therapeutics.

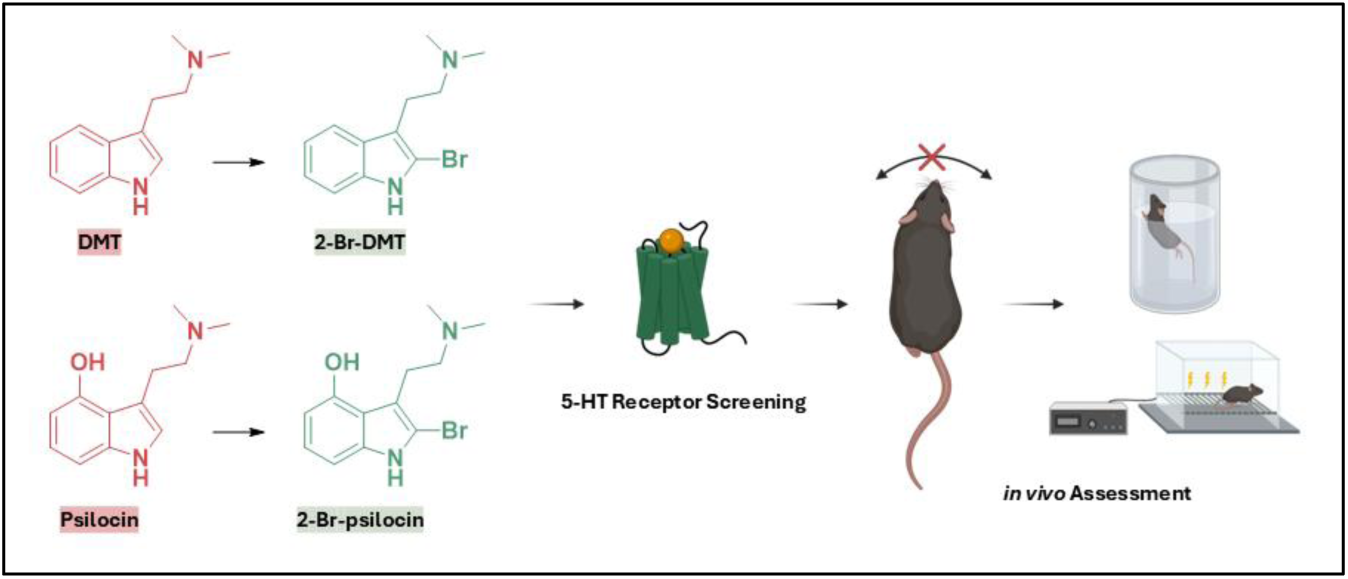

## Introduction

Natural products have played a central role in modern drug discovery, with 84% of FDA approved drugs for central nervous system (CNS) disease indications being natural products or derivatives thereof, and psychedelics are no exception.^1^ Some psychedelics, such as psilocybin, found in various mushroom species, as well as DMT, present in plants and animals, are found in nature.^2^ When administered in controlled environments, psychedelics are documented to facilitate individual and group healing, conflict resolution, and community cohesion.^3–6^ The therapeutic potential of psychedelics builds upon millennia-old indigenous practices of communal psychedelic ceremonies, practices that persist today.^3,7–9^ Innovation in the past century has advanced the development of semi-synthetic and synthetic psychedelics, yielding key insights into structure-activity relationships (SARs) for the major chemotypes of psychedelics.^10–14^

The serotonin (5-HT) system consists largely of G protein-coupled receptors with diverse physiological roles, including regulation of mood, cognition, cardiovascular function, and inflammation. Serotonergic psychedelics, such as lysergic acid diethylamide (LSD), psilocybin, and DMT (**Figure 1**) are best known for their strong effects on cognition, perception, mood, and sense of self, mediated through agonist actions at the 5-HT_2A_ receptor.^15^ In clinical settings, psychedelics have demonstrated significant rapid-acting and durable clinical potential in treating major psychiatric conditions, pain, sleep disorders, neuroinflammation, and neurodegenerative diseases.^16–30^

**Figure 1.**
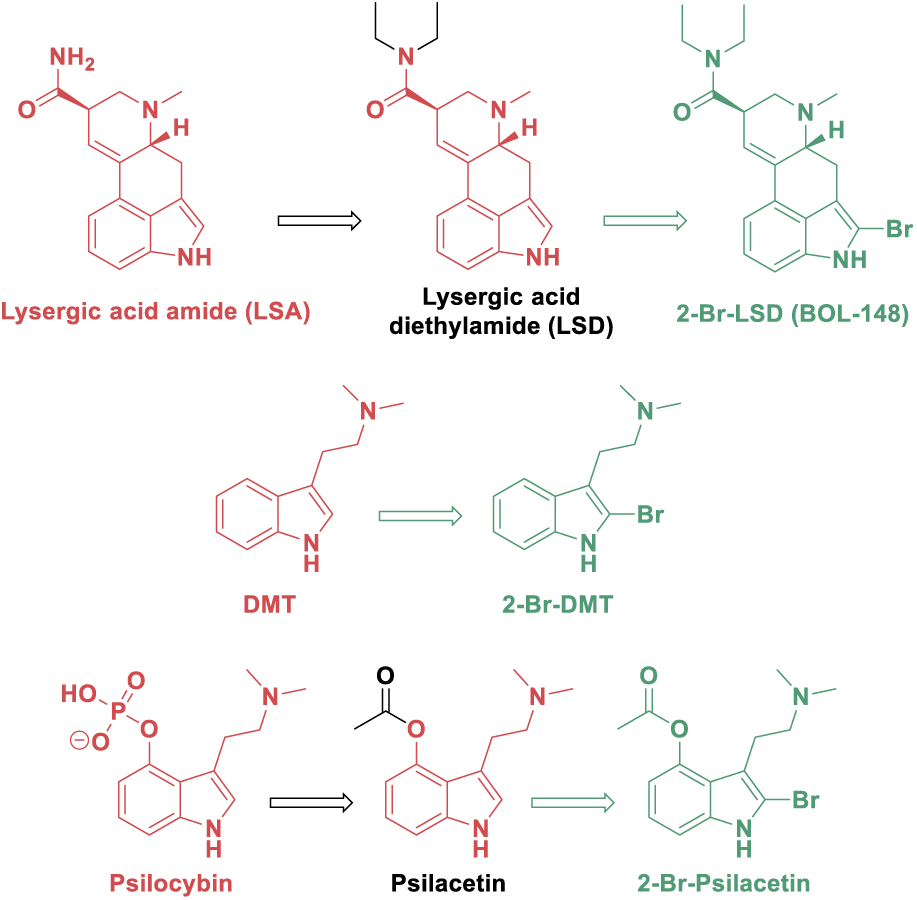
Structures of naturally derived serotonergic tryptamine psychedelics (red), semi-synthetic (red/black), and halogenated next generation analogs (green)

Beyond 5-HT_2A_, receptors such as 5-HT_2C_, 5-HT_2B_, and 5-HT_6_ are of particular interest due to their function in safety and therapeutic potential. ^31–36^ For instance, 5-HT_2C_ plays a role in obesity, schizophrenia, and Alzheimer’s related psychosis.^37^ Agonism at 5-HT_2B_ is associated with cardiac valvulopathy and pulmonary hypertension.^38,39^ Therefore, in drug design, reducing agonist activity or shifting drug interactions towards antagonism at 5-HT_2B_ would be ideal due to safety concerns.^40,41^ The 5-HT_6_ receptor, which seems to be exclusively expressed in the CNS, has emerged as a promising target for cognitive enhancement and neuropsychiatric disorders, with both agonists and antagonists showing benefit in different preclinical models.^42–44^

The pharmacological promiscuity of psychedelics has motivated efforts to develop psychedelic-inspired compounds that offer therapeutic benefits without inducing psychedelic-like subjective effects.^45–51^ For instance, LSD, synthesized from naturally derived ergotamine and lysergic acid, was developed into non-psychoactive derivatives (i.e., 2-Br-LSD and lisuride, to list a few).^52,53^ BOL-148, known as 2-Br-LSD, was first synthesized in 1950’s and has shown potential in treating cluster headaches. Recent research shows 2-Br-LSD has a safer cardiotoxicity profile since it lacks agonism at 5-HT_2B_ and does not potently bind to hERG.^53^ 2-Br-LSD also has a distinct serotonergic pharmacology compared to LSD and reverses behavioral effects of chronic stress in mice.^53^ Unlike LSD, 2-Br-LSD has reduced Gq signaling efficacy at 5-HT_2A_ acting as a partial agonist well below the threshold of known psychedelic activation of 5-HT_2A_, which is predictive of psychoactive potential in humans.^54^

The effects of ring substitution at the 2-position of tryptamines generally disrupts key interactions within the orthosteric binding pocket of the 5-HT_2A_ receptor (D155^3.32^ and S242^5.46^) reducing affinity and in most cases, may convert compounds to 5-HT_2A_ antagonists.^55–57^ This information, combined with the promising research on 2-Br-LSD, guided our design and synthesis of 2-halogenated tryptamines (**Figure 1**). Herein, we present the synthesis of 2-halogenated derivatives of DMT and psilacetin, a synthetic psilocybin equivalent. These compounds were evaluated for receptor binding affinities across aminergic receptor families, along with functional activity at the serotonin receptors. Additionally, behavioral studies were performed in mice to assess potential psychoactive effects via head-twitch response (HTR), cognition, and social interaction.

## Results and Discussion

### Design and Synthesis of 2-Halogenated Tryptamines

Psilacetin is a synthetic equivalent to psilocybin *in vitro* and in human trip reports.^58^ It was previously synthesized by David Nichols group through a one pot debenzylation/acetylation of O-benzylpsilocin.^59^ For this study, psilocin was synthesized via the Speeter-Anthony route, followed by acetylation with acetyl chloride to produce psilacetin (**Scheme S1**).^60^

Bromination at the 2-position of C3-substituted indoles can be achieved with *N*-bromosuccinimide (NBS) which was the original method utilized to synthesize 2-Br-LSD.^61^ Although this worked for 2-Br-LSD, it is not as straightforward to apply this method to tryptamines due to the formation of oxindoles and the potential cyclization product (**Scheme 2**).^62,63^ Initially, halogenation with the corresponding *N*-halosuccinimide (NXS) exclusively formed cyclized byproduct **1a** and **3a**. Therefore, a different brominating agent, which forms bromine *in situ*, was used to brominate psilacetin and DMT (**Scheme S1-2**). Finally, the hydrolysis of the ester was achieved in 2M aqueous hydrobromic acid to give 2-Br-psilocin (**4**). The synthesis of 2-Cl-DMT (**2b**) was successfully achieved via chlorination of the tryptamine hydrochloride salt and *N*-chlorosuccinimide (NCS) in glacial acetic acid and formic acid, followed by reductive amination (**Scheme 1**).^64^ The strong acidic conditions kept the amine protonated, thereby reducing its nucleophilicity and preventing cyclization from occurring.

**Scheme 1.**
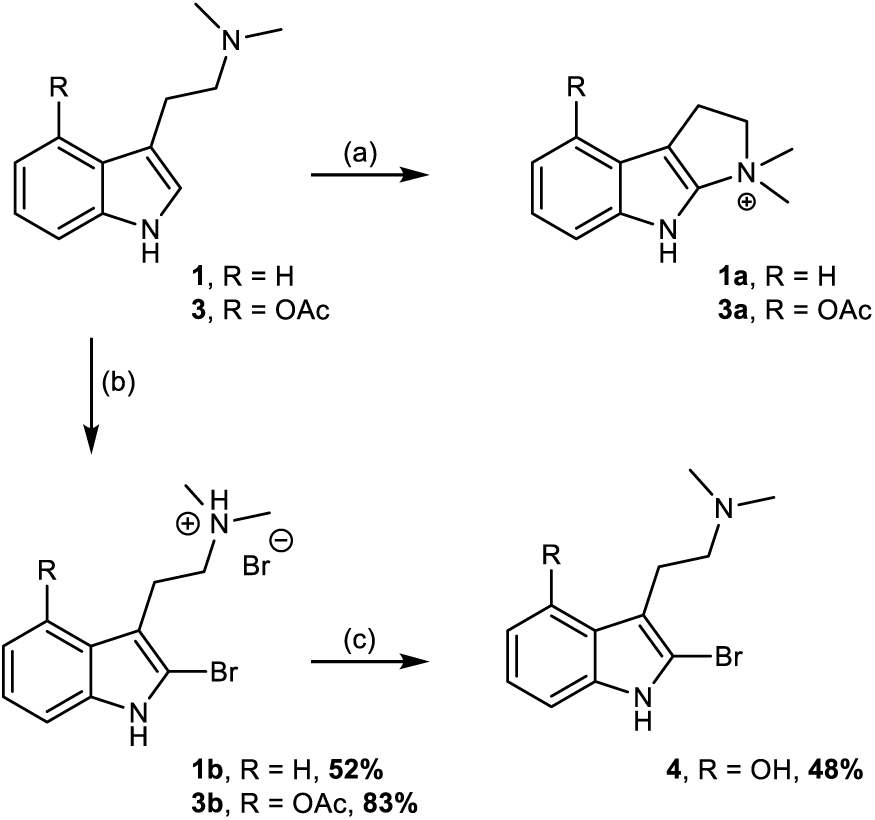
(a) NBS, DCM, 20 min; (b) PhMe_3_NBr_3_, DCM, 20 min; (c) 1M aq. HBr, 70 °C, 2h

### Receptor Binding Profiles and Mouse Brain Binding

In most binding pockets of 5-HT receptors, the salt bridge formed by the conserved aspartate residue (D155^3.32^) with the amine appear essential for anchoring the tryptamine in place.^65^ It is well known that 2-aryl-tryptamines show increased selectivity for 5-HT_6_ receptors.^66^ The 2-halogenated tryptamines were initially tested in comprehensive binding screens to determine potentially relevant pharmacological targets of interest and compare halogenated compounds to their template structures. Across 5-HT receptors, the 2-position halogenated analogues lacked or had reduced affinity across 5-HT_1_ receptors, while showing similar or reduced binding affinities at other 5-HT receptors relative to DMT and psilocin (**Table 1**). Notably, 2-position halogens retained affinities for 5-HT_6_ and 5-HT_7a_ receptors, where the analogues generally displayed the highest affinities vs. all other 5-HT receptors. It has been demonstrated that psilacetin rapidly converts to psilocin under physiological conditions.^58^ As a result, 2-Br-psilocin was evaluated for its binding affinity at serotonin receptors (**Table S2)** and served as the representative active metabolite form of 2-Br-psilacetin for further *in vitro* and *in vivo* studies.^67^

**Table 1:**
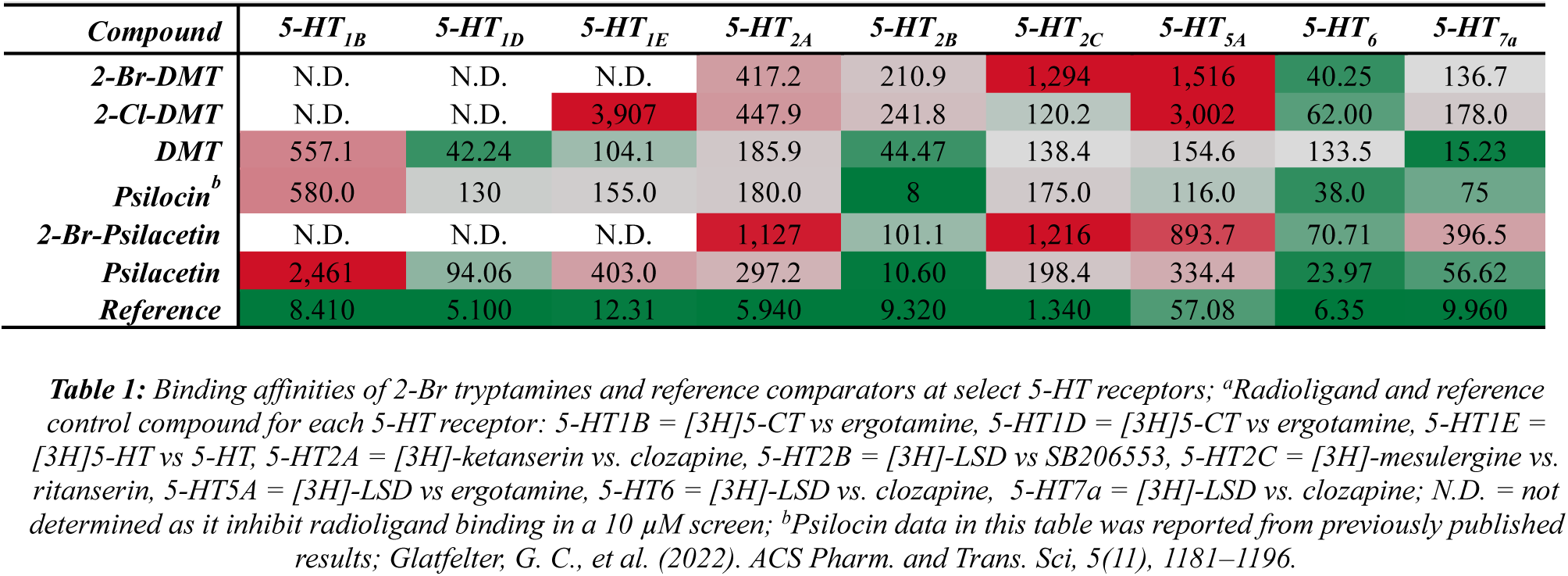
Affinities of test compounds in radioligand binding assays (K_i_, nM) for human 5-HT receptors^a^.

Outside the 5-HT system, the halogenated analogues showed additional interactions with histamine, sigma, alpha adrenergic, dopamine, and muscarinic receptors, generally with Ki values ≥1 µM, although several improvements in affinity were noted (**Table S1-3**). Compared to DMT, 2-Br-DMT and 2-Cl-DMT showed increased affinity for histamine H_2_ (K_i_ = 678 – 752 vs. 3,645 nM for DMT), alpha_2A_ (K_i_ = 670 – 866 vs. 2,034 nM), and dopamine D_3_ (K_i_ = 462 – 602 vs. 1,267 nM) receptors. Overall, target screening data suggests that 2-position halogenation reduces promiscuity of the compounds at 5-HT_1_ receptors. Moreover, the compounds have several potentially relevant interactions with histamine, sigma, alpha, dopamine, and muscarinic receptors.

Prior to mouse studies, several of the compounds were assessed for abilities to compete for radioligand binding at 5-HT_1A_ ([^3^H]8-OH-DPAT) and 5-HT_2A_ ([^3^H]M100907) receptors in mouse brain using established methods.^34^ At m5-HT_1A_ and m5-HT_2A_, halogenation of DMT or psilocin resulted in reduced affinities without exception (**Figure S1**, **Table S4**). More specifically, template compounds DMT and psilocin displayed low to mid nM affinities at both receptors, while the halogenated analogues displayed high nM to µM affinities. Overall, these data align with observations from receptor binding screening results using human receptors, showing that these compounds target 5-HT_1A_ and 5-HT_2A_ with reduced affinity relative to the non-halogenated template compounds.

### G protein-dissociation

Across all the 5-HT receptors, the 2-halogenated tryptamines displayed weak potency and efficacy at 5-HT_1A_ (EC_50_ = 1600-5400 nM, E_max_ = 68-83%) (**Figure 2A-D**, **Table 2**). However, this is not the case at the other 5-HT_1_ receptors. For 5-HT_1B_, the test compounds showed moderately potent agonism (EC_50_ = 88-730 nM, E_max_ = 41-84%), which is similar and slightly higher for 5-HT_1D_ (EC_50_ = 67-300 nM, E_max_ = 67-110%). In both cases, 2-Br-psilocin has the highest activity at these 5-HT_1B/1D_ receptor types. Finally, at 5-HT_1e_, all test compounds are not very potent but display almost full agonist activity relative to 5-HT (EC_50_ = 270-1000 nM, E_max_ = 87-100%).

**Figure 2:**
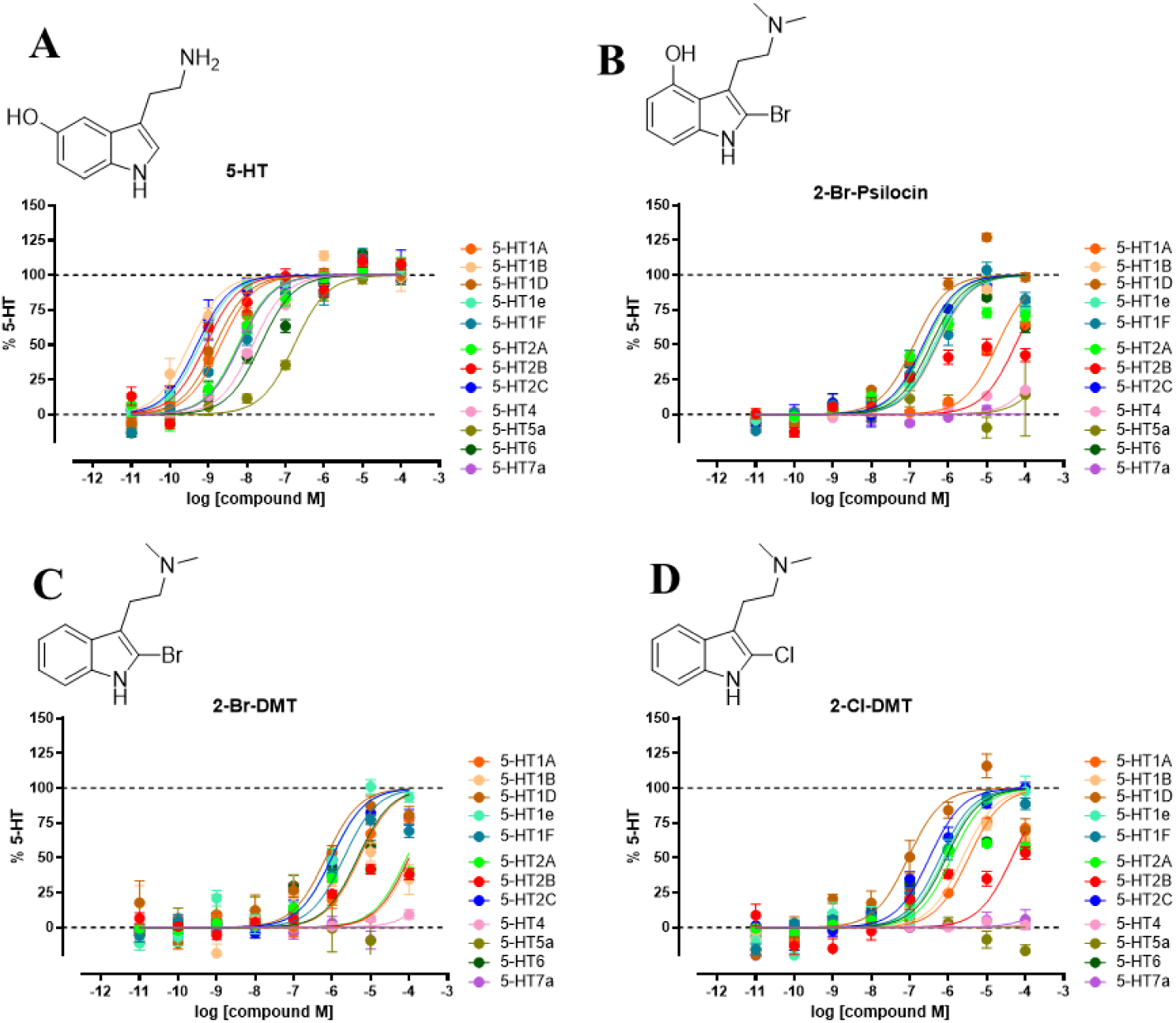
(A-D) 5-HT GPCRome activity for 2-substituted analogues. Data are normalized to percent 5-HT response and represent the mean and SEM from three or more independent experiments.

**Table 2.**
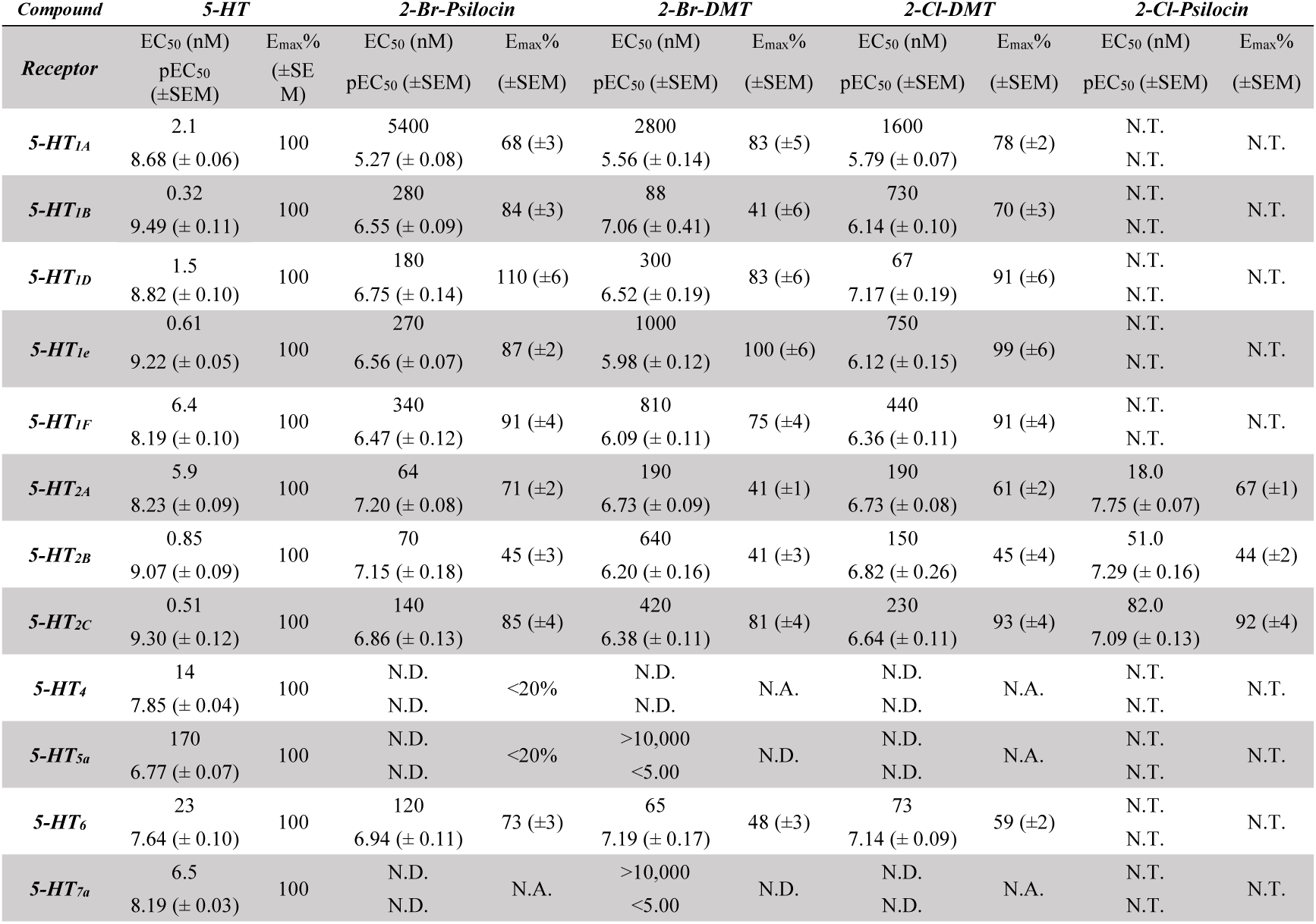
Parameter estimates for 5-HT GPCRome G protein dissociation activity as measured by BRET. Data represents the mean and SEM from three or more independent experiments. N.A. not active, N.D. not determined, N.T. not tested.

Moderate partial agonism was observed at 5-HT_2B_ receptors (EC_50_ = 51-640 nM, E_max_ = 41-45%). At 5-HT_2C_, the 2-Cl derivatives showed modest increases in potency and efficacy compared to their 2-Br counterparts (2-Cl-psilocin, EC_50_ = 82 nM, E_max_ = 92% vs. 2-Br-psilocin, EC_50_ = 140 nM, E_max_ = 85% and 2-Cl-DMT, EC_50_ = 230 nM, E_max_ = 93% vs. 2-Br-DMT, EC_50_ = 420 nM, E_max_ = 81%). Finally, at 5-HT_2A_, the 2-halogenated tryptamines analogs showed moderate potency (EC_50_ = 18 - 64 nM) and weak partial agonism for 2-Br-DMT (E_max_ = 41%) vs. the rest of the test compounds (E_max_ = 61-71%). At the 5-HT_6_ receptor, 2-Br-psilocin maintains moderately potent partial agonism (EC_50_ = 120 nM, E_max_ = 73%) compared to 2-Br-DMT and 2-Cl-DMT which are potent but were weaker partial agonists (EC_50_ = 65 nM, E_max_ = 48%, EC_50_ = 73 nM, E_max_ = 59%, respectively). All analogues displayed no agonism at 5-HT_7a_. Direct comparison with parent compounds demonstrated that 2-Br-DMT exhibited broad reductions in potency and efficacy as an agonist at 5-HT_2A_, 5-HT_2B_, and 5-HT_6_, while 2-Br-psilocin selectively preserved 5-HT_6_ activity (**Figure 3**, **Table 2**). Among the receptors tested where agonist activity was detected, agonism at 5-HT_2B_ is the weakest (E_max_ = 45%) as compared to the other 5-HTRs (E_max_ = 71-85%).

**Figure 3.**
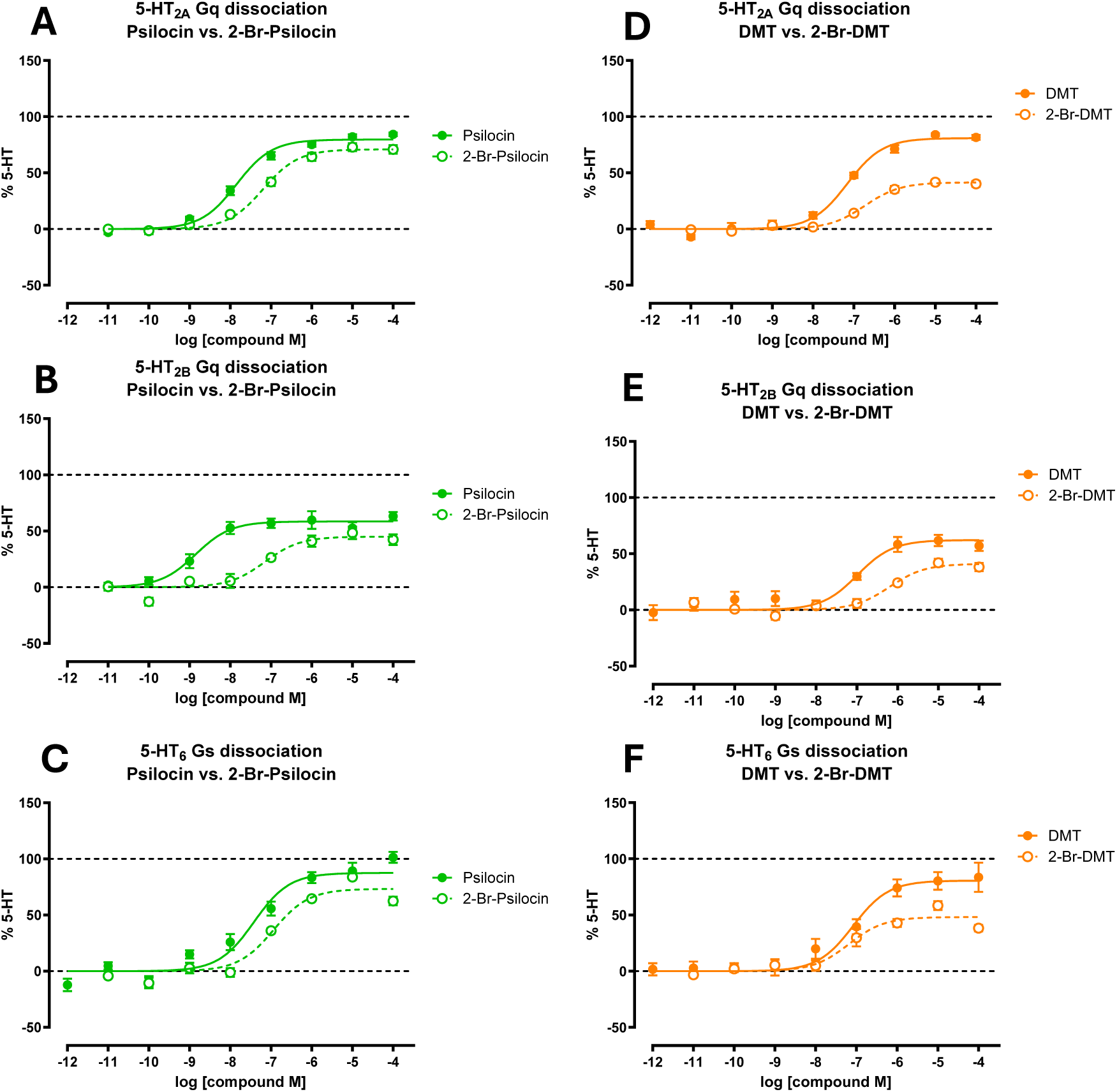
Effect of 2-Br substitution on G-protein dissociation as determined by BRET in 5-HT_2A_, 5-HT_2B_, and 5-HT_6_ for (A-C) 2-Br-psilocin (green) and (D-F) 2-Br-DMT (orange). Data represent the mean and SEM from three or more independent experiments.

### Orthogonal assays (Ca^2+^ flux and cAMP signaling)

Second messenger assays were performed for 2-Br-psilocin at select 5-HT_1_ receptors, as well as 5-HT_2_, 5-HT_6_, and 5-HT_7a_ receptors. Overall, 2-Br-psilocin displayed weak efficacy in G_q/11_-mediated calcium flux at 5-HT_2_ receptors (**Table 3, Figure S2**). 2-Br-psilocin showed weak potency measuring cAMP inhibition at the 5-HT_1A_ receptor but was quite efficacious to activate 5-HT_1B_ (EC_50_ = 34.8 nM, E_max_ = 88%) and 5-HT_1F_ receptors (EC_50_ = 35.0 nM, E_max_ = 110%), consistent with their G protein dissociation activities. 2-Br-psilocin shows potent low nanomolar potency at 5-HT_6_ receptor (EC_50_ = 4.10 nM) and with almost full agonist efficacy (E_max_ = 93%). Along with the other 2-halogenated tryptamines, 2-Br-psilocin showed weak inverse agonism at 5-HT_7a_, an effect similarly reported for 2-Br-LSD that was much more potent.^53^

**Table 3:**
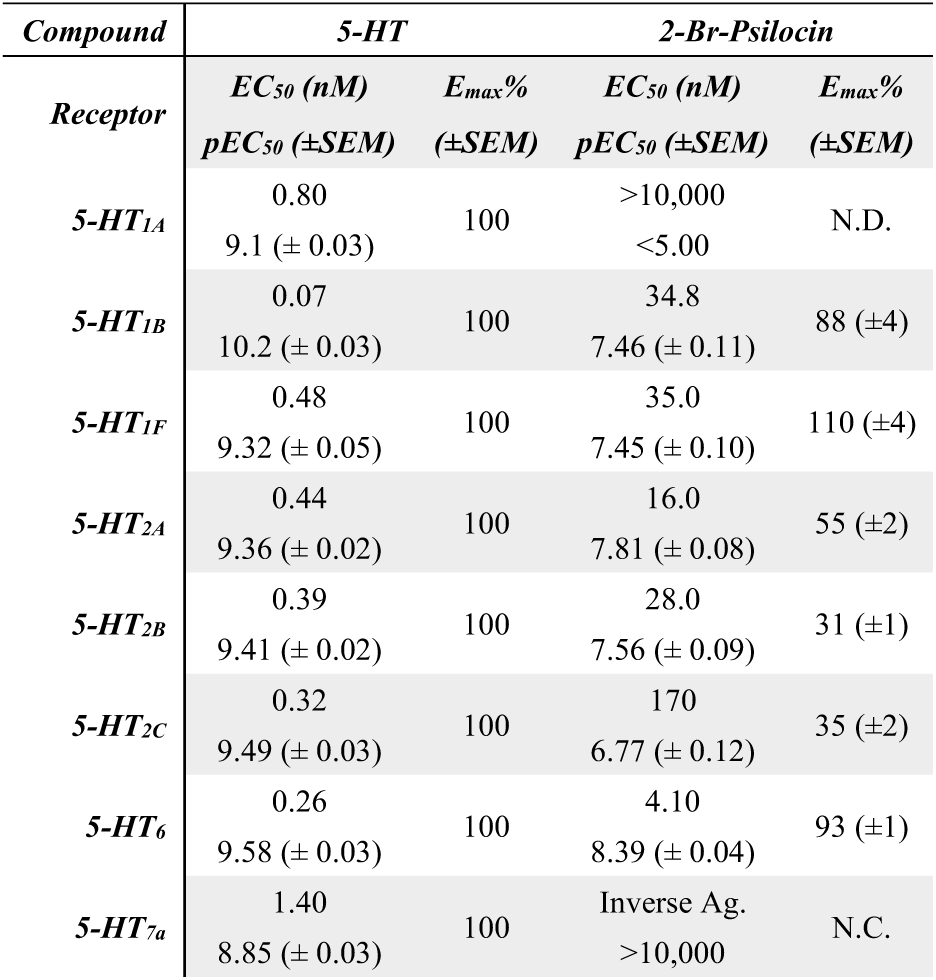
Parameter estimates for 5-HT receptor second messenger activities measuring Gi/o-mediated cAMP inhibition for 5-HT_1A_, 5-HT_1B_, and 5-HT_1F_, Gq/11-mediated calcium flux for 5-HT_2A_, 5-HT_2B_, and 5-HT_2C_, and Gs-mediated cAMP accumulation for 5-HT_6_ and 5-HT_7a_. Data represent the mean and SEM from three or more independent experiments.

**Table 3:**
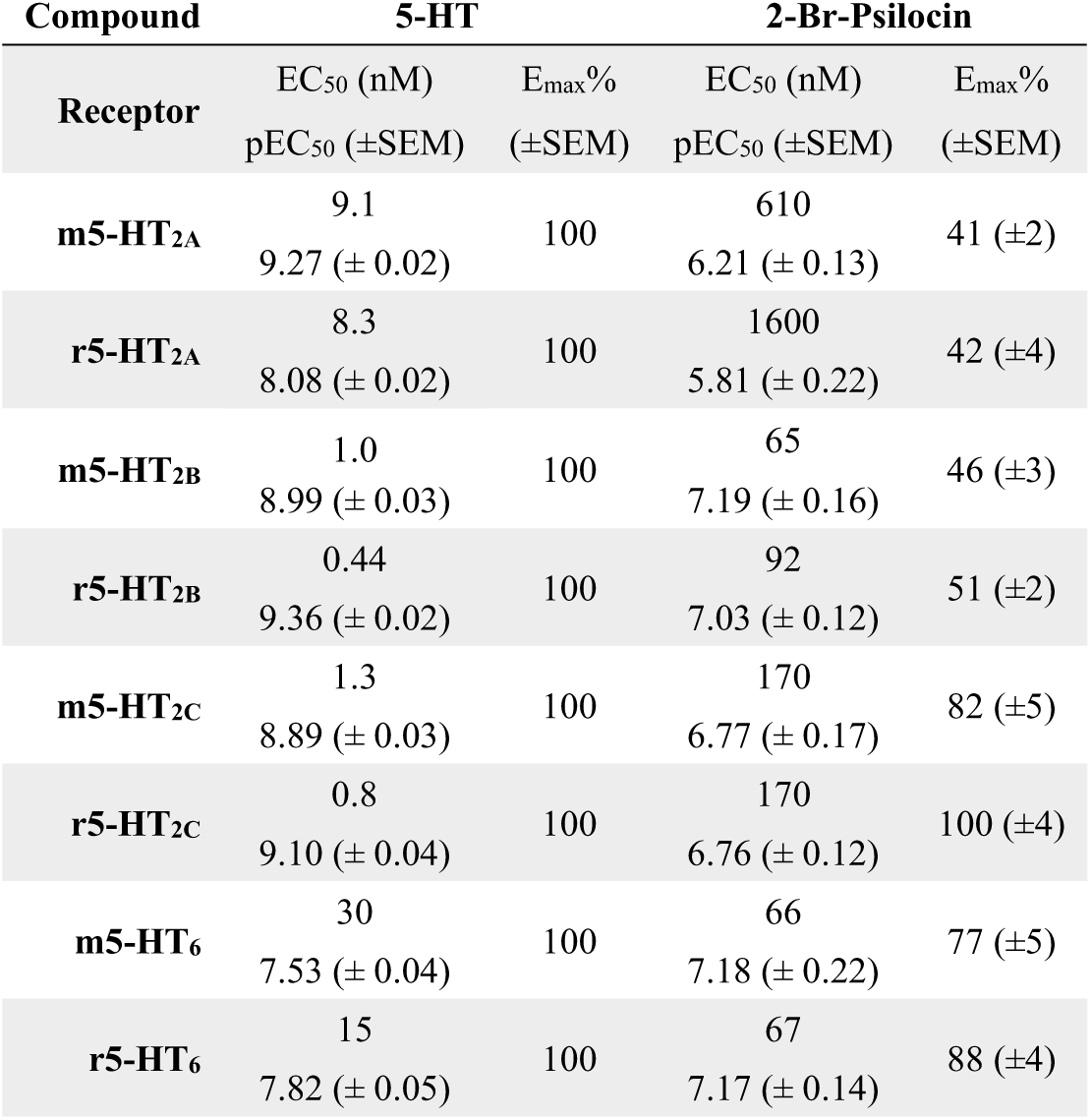
Parameter estimates for rodent species 5-HT_2A_, 5-HT_2B_, 5-HT_2C_, and 5-HT_6_ receptors measuring G protein dissociation activity as measured by BRET. Data represent the mean and SEM from three or more independent experiments.

The polypharmacological profile examining second messenger assays of 2-Br-psilocin can be summarized as a weak partial agonist at 5-HT_2A,_ 5-HT_2B,_ and 5-HT_2C_, an almost full agonist at 5-HT_6_ and 5-HT_1B_, as well as a full agonist at 5-HT_1D_. It is well known that agonist actions at 5-HT_1B/1D/1F_ are all hypothesized to be targets of anti-migraine medications.^68^ Furthermore, the 5-HT_1B_ receptor plays a role in modulating amygdala-related behaviors by psilocybin.^69^ These results were also aligned with orthogonal assays tested in an analogous assay (**Figure S3, Table S5**).

### 2-halogenation decreases 5-HT_2B_ activity

The 2-halogenated tryptamines potentially have a safer cardiotoxicity profile as compared to their parent compounds. In **Table 1**, the test compounds had a ∼5-fold lower binding affinity at 5-HT_2B_ as compared to DMT, psilocin, and psilacetin. Furthermore, all the 2-halogenated tryptamines are weak partial agonists (E_max_ = ∼41-45%). 2-brominated DMT and psilocin analogues have reduced potencies and efficacies as compared to their parent psychedelic compounds **(Figure 3B, E**). There was no hERG activity displayed by any of the test compounds except 2-Cl-DMT (IC_50 =_ ∼5 µM) in a radioligand binding assay (**Table S6**). The weaker binding affinities observed at 5-HT_2B_ and lack of activity at hERG supports a safer cardiotoxicity profile for the halogenated tryptamines.^70^ binding assay (**Table S6**). The weaker binding affinities observed at 5-HT_2B_ and lack of activity at hERG supports a safer cardiotoxicity profile for the halogenated tryptamines.^70^

### Species differences in 5-HTR functional data

There are relevant species differences in amino acid sequences of the human and mouse 5-HT_2A_ receptors. The key difference lies in residue S242^5.46^, which in mice is an alanine, but in humans it is a serine.^71^ This residue appears important in ligand binding interactions in the orthosteric site of 5-HT_2A_, and can change affinity, potency, and efficacy in some cases.^72,73^ Given the significance of these species differences, we examined the functional activity of 2-Br-psilocin at human vs. rodent 5-HT receptors. **Table 3** lists the G protein dissociation activity for 2-Br-psilocin in mouse and rat serotonin receptors, and **Table 2** lists the human serotonin receptor profile.

The functional activities of 2-Br-psilocin at 5-HT_6,_ 5-HT_2B_, and 5-HT_2C_ are similar in rodents and humans (**Figure 4**), whereas the compound has reduced agonism at m5-HT_2A_ and r5-HT_2A_ (EC_50_ = 610 nM, E_max_ = 41%, EC_50_ = 1600 nM, E_max_ = 42%, respectively) as compared to the human species (EC_50_ = 64 nM, E_max_ = 71%). The agonist activity of 2-Br-psilocin at 5-HT_2C_ is similar for human and mouse receptors (EC_50_ = 140 nM, E_max_ = 85%, EC_50_ = 170 nM, E_max_ = 82%, respectively) but it displays full agonism at r5-HT_2C_ (EC_50_ = 170 nM, E_max_ = 100%). For 5-HT_6_, the activity was similar across all species (EC_50_ = 67-120 nM, E_max_ = 73-88%). In general, potency and efficacy of 2-Br-psilocin were weaker for rodent variants compared to the human 5-HT_2A_ receptor (**Table 3**). This is important to note for any preclinical behavioral assays that are primarily modulated by these receptors, as efficacy in rodent models may not translate directly to humans.

**Figure 4.**
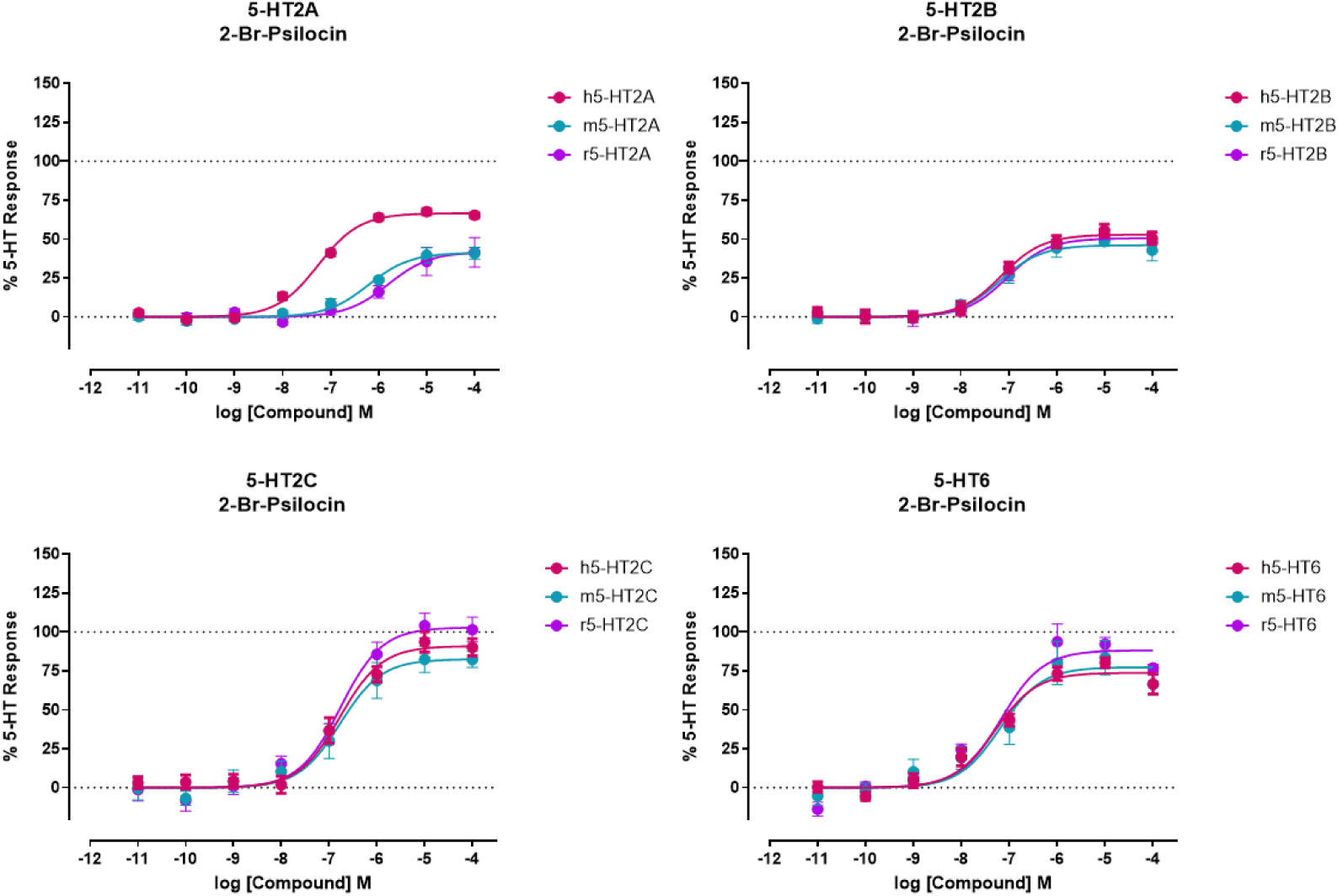
Effect of rodent species on 5-HT receptor activity; G protein-dissociation BRET experiments for 2-Br-psilocin in human, mouse, and rat serotonin receptors at (A) 5-HT_2A_, (B) 5-HT_2B_, (C) 5-HT_2C_, and (D) 5-HT_6_. Data represent the mean and SEM from three or more independent experiments.

### Pharmacokinetic profile of 2-Br-psilacetin

Across all test compounds, 2-halogenation greatly increased permeability, as determined through blood-brain barrier (BBB) parallel artificial membrane assay (PAMPA) (**Table S7**). Upon oral administration of 2-Br-psilacetin in rats, it is quickly converted to 2-Br-psilocin and is ∼66% orally bioavailable (**Table S8**). **Table S9** shows the exposure levels of 2-Br-psilocin in the brain, as further evidence that 2-Br-psilacetin reaches the brain and is in the form of 2-Br-psilocin. This was predicted to be the case since psilacetin is also quickly hydrolyzed under physiological conditions to form psilocin.^58^

### Head Twitch Response (HTR) assessment of 2-Br-psilacetin

The HTR in mice is a rapid, high-frequency rotational head movement mediated by cortical 5-HT_2A_ receptor activation that represents the most pharmacologically validated and translationally relevant preclinical assay for predicting the potential for psychedelic-like psychoactivity in humans (refs).^74–77^ Given this well-established ability to gauge relative potencies of 5-HT_2A_ agonists in C57BL/6J mice for psychedelic effects in humans, HTR quantification was used to evaluate the psychedelic-like effects of 2-Br-psilocin prodrug, 2-Br-psilacetin compared to the psilocin prodrug, psilacetin.^77^ Using previously published methods examining similar tryptamine psychedelics, the acute effects of psilacetin and 2-Br-psilacetin on HTR, motor activity, and thermoregulation over a 30 min testing period were examined in C57BL/6J mice.^34,78–81^ Psilacetin produced a dose-dependent biphasic HTR curve with an ascending limb potency (ED_50_ = 0.26 mg/kg) and maximal number of events (E_max_ = 34 total HTR counts) consistent with previous studies (**Figure 5A**).^34,82^

**Figure 5:**
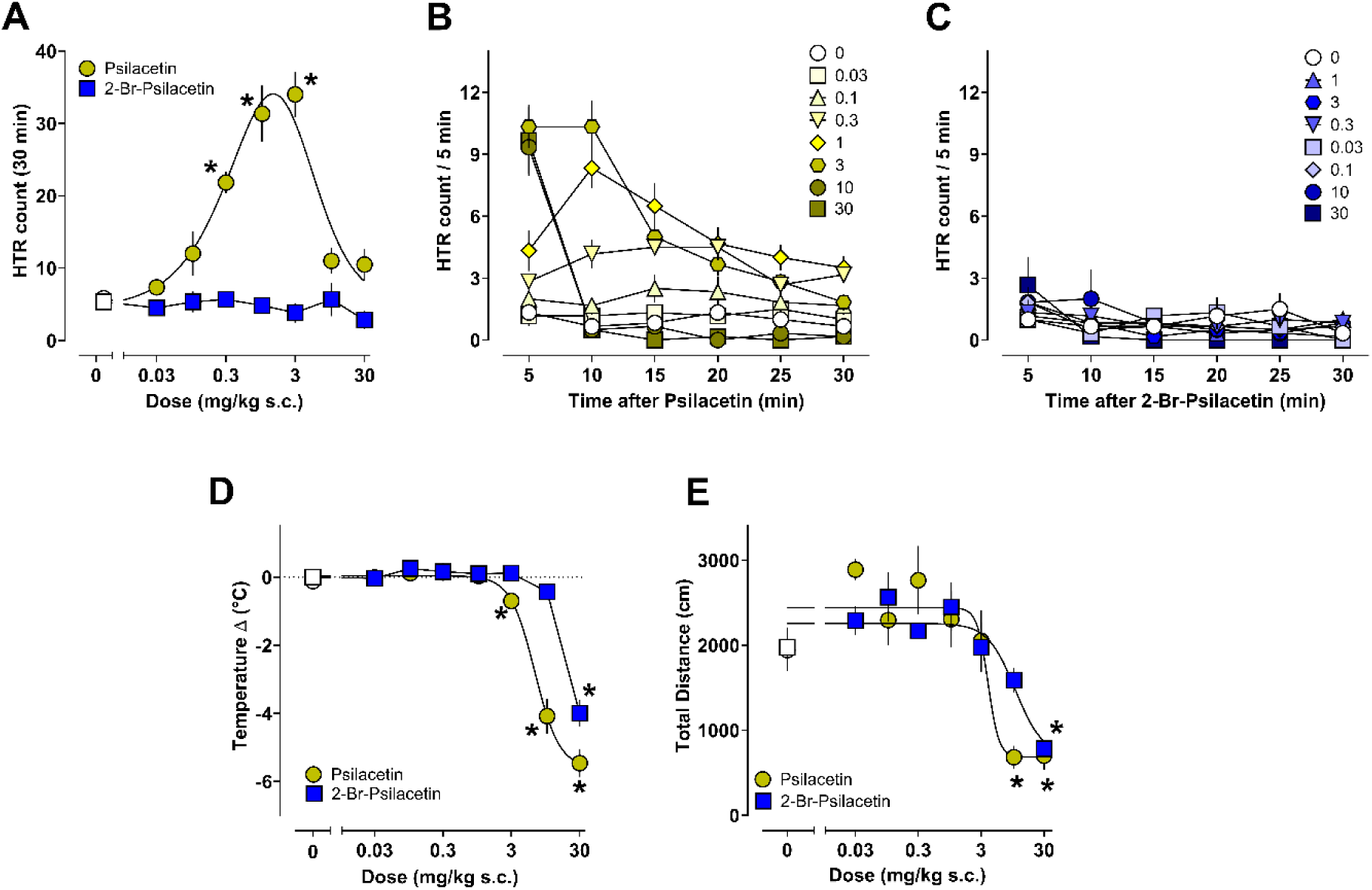
Psychedelic-like effects of psilacetin and 2-Br-psilacetin in the HTR assay. All values are mean ± SEM and represent n = 6 mice/dose. Asterisks represent statistical differences vs. vehicle controls (p < 0.05).

In contrast, 2-Br-psilacetin did not produce a significant increase in the number of HTRs vs. vehicle controls at any dose tested, while psilacetin did from 0.3-3 mg/kg. The time-course of effects for both compounds on HTR are shown in **Figure 5B-C**, with 2-Br-psilacetin only producing vehicle levels throughout the session (i.e. ∼1 - 3 HTRs/5 min). Psilacetin produced consistent increases in HTR frequency (∼5-10 HTRs/ 5 min) from 0.3-3 mg/kg that largely resolved by the end of the session, but at higher doses (i.e. 10-30 mg/kg) the response quickly peaked at 5 min before returning to vehicle levels due to onset of hypothermic and hypolocomotor effects as previously reported (**Figure 5D-E**). Given the utility of the HTR assessment for gauging potencies for psychedelic subjective effects in humans,^83^ these data suggest that relative to psilacetin, 2-Br-psilacetin may lack or have reduced psychoactive and psychedelic effects.

Since 2-Br-psilacetin did not elicit the HTR and 2-Br-psilocin displayed a partial agonist profile at 5-HT^2A^ in functional *in vitro* assays, we surmised that 2-Br-psilacetin may be a partial agonist/antagonist *in vivo*. To test this, the ability of 2-Br-psilacetin (10 mg/kg) to block the HTR induced by the 5-HT^2^ selective ligand, 2,5-dimethoxyamphetamine (DOI, 0.5 mg/kg) was examined. Administration of DOI induced a peak in HTR counts within 10-15 mins post injection (∼12-15 HTRs/5 min) relative to vehicle and 2-Br-psilacetin control conditions (∼1-3 HTRs/5 min, **Figure 6A**). Importantly, no basal HTR differences were observed across conditions after injection of vehicle or 2-Br-psilacetin (**Figure 6B**). In contrast to mice pretreated with vehicle prior to DOI, mice pretreated with 2-Br-psilacetin prior to DOI administration displayed less HTRs (∼ 3-5 HTRs/5 min) that were significantly reduced from DOI alone (**Figure 5C**). Notably, there were no hypothermic or hypolocomotor effects observed because of 2-Br-psilacetin treatment (**Figure 6D-G**), supporting that the reductions in DOI-induced HTR could be 5-HT^2A^-mediated. However, given the polypharmacological profile of 2-Br-psilocin, effects of the drug at other 5-HT receptors may also play a role in its ability to attenuate behavioral effects of DOI in the HTR paradigm.^34^ Notably, although 2-Br-psilacetin reduces the HTR in mice as compared to psilacetin, a slight difference in agonist activity at human vs. mouse 5-HT^2A^ was observed in vitro. This difference may therefore limit interpretations of HTR data with 2-Br-psilacetin, given interactions with 5-HT^2A^ residue S242^5.46^ are seemingly pertinent. Given the increased potency and efficacy of 2-Br-psilocin at h5-HT^2A^ relative to m5-HT^2A^, further studies will be needed to account for translational effects due to species differences.^71^

**Figure 6:**
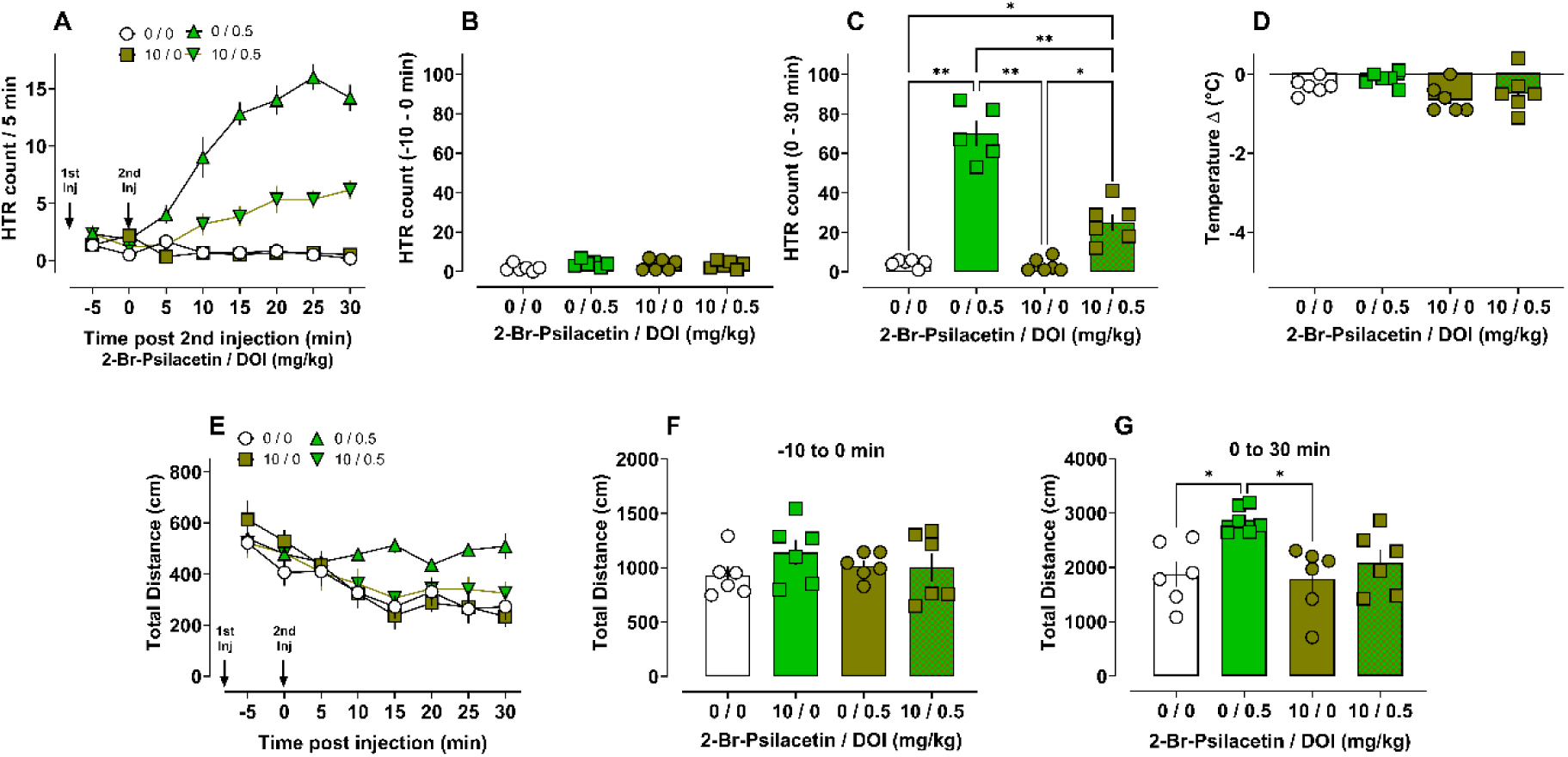
Psychedelic-like effects of DOI (0.5 mg/kg) are reduced by pretreatment with 2-Br-psilacetin (10 mg/kg). Time-course for effects on the HTR (A) and total HTRs for the pretreatment period (B) as well as the 30 mins post DOI treatment (C). Summary data for hypothermic (D) and locomotor effects (E – G). All data are mean ± SEM of n = 5 – 6 mice/dose. Asterisks indicate statistical differences between groups (p < 0.05).

Due to the interesting polypharmacology profile of 2-Br-psilacetin, we employed a comprehensive behavioral battery in mice including: Pavlovian cued and contextual fear conditioning, Porsolt forced swim test (PFST), social behavior in the open field test (OFT), and acoustic startle response (ASR) with prepulse inhibition (PPI) (**Figures 7-9**). All of these studies were completed in a single cohort of male mice. Drug treatment began 3 days prior to the first behavioral test, and was given i.p. daily for 18 days, within one hour of the start of the behavioral test for each day. Vehicle (5% EtOH, 12% PEG400, 2% Tween80, and 20% propylene glycol in physiological saline) was used as a negative control, and 20 mg/kg psilacetin was used as a positive control.

**Figure 7.**
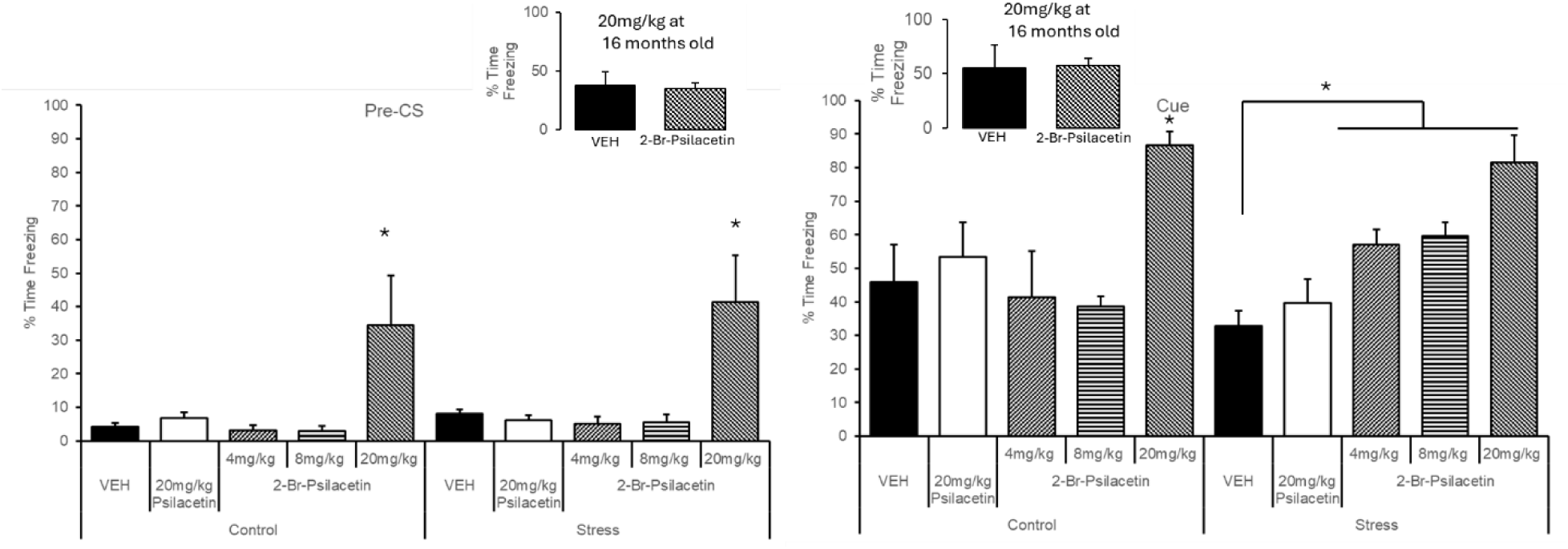
Fear conditioning in 2-3-month-old C57BL/6 mice (A) preconditioned stimulus and (B) cued stimulus, inset chart for each condition corresponds to data in 16-month-old C57BL/6 mice administered 20 mg/kg 2-Br-psilacetin

### Fear conditioning

In the training session for fear conditioning, 20 mg/kg 2-Br-psilacetin tended to increase freezing compared to VEH, but there were no significant differences between groups. However, 20 mg/kg 2-Br-psilacetin increased freezing compared to VEH in the control condition during the tone-shock pairing period (p<0.05), with a similar trend in the stress condition (p=0.06) (**Figure S4A-B).** Only 20 mg/kg 2-Br-psilacetin increased freezing compared to vehicle in the control condition during the context test (p<0.05) (**Figure S4C)**. In the pre-cued stimulus (CS) period of the cue test, 20 mg/kg 2-Br-psilacetin increased freezing compared to VEH in both the control and stress conditions (p<0.05). 20 mg/kg 2-Br-psilacetin also increased freezing compared to vehicle in the control condition during the CS presentation (p<0.05), whereas all doses of 2-Br-psilacetin increased freezing in the stress condition (p=0.06). Together, the results suggest that the 20 mg/kg dose of 2-Br-psilacetin produces a general increase in freezing behavior, possibly related to lethargy rather than anxiety, based on the baseline freezing in the training period and on other data we have acquired. In contrast, 4 and 8 mg/kg 2-Br-psilacetin produce an increase in freezing to the cue, indicating an improvement in hippocampus-independent cued learning (**Figure 7A-B, Figures S4A-C).** To determine whether lower doses of 2-Br-psilacetin might be effective in an aged model, in which cognitive deficits remove any potential ceiling effects, we also tested this dose against VEH in the same paradigm, using 16 month-old C57BL/6J mice (**Figure 7A-B, inset**), but found no rescue at the older age.

### Porsolt Forced Swim Test

Although there was no effect of drug treatment alone (p>0.05), there were significant effects of stress conditioning and treatment x stress interactions. For Day 1 immobile time, the interaction, F(4,45)=3.13, p=0.026 indicated that 8 mg/kg 2-Br-psilacetin reduced the total time immobile in the group exposed to restraint stress compared to the VEH-treated restraint group (p<0.05). For Day 1 latency to immobility, the interaction, F(4,45)=3.30, p=0.20 indicated that the control group administered psilacetin had a longer latency compared to the VEH-treated control, whereas that 8 mg/kg 2-Br-psilacetin reduced the latency in the group exposed to restraint stress compared to the VEH-treated restraint group (p<0.05) (**Figure 8A-B**). For Day 2 immobile time, the interaction, F(4,45)=3.98, p=0.010 indicated that 8 mg/kg 2-Br-psilacetin increased immobile time compared to VEH in the control groups but reduced the immobile time in the group exposed to restraint stress compared to the VEH-treated restraint group (p<0.05).

**Figure 8.**
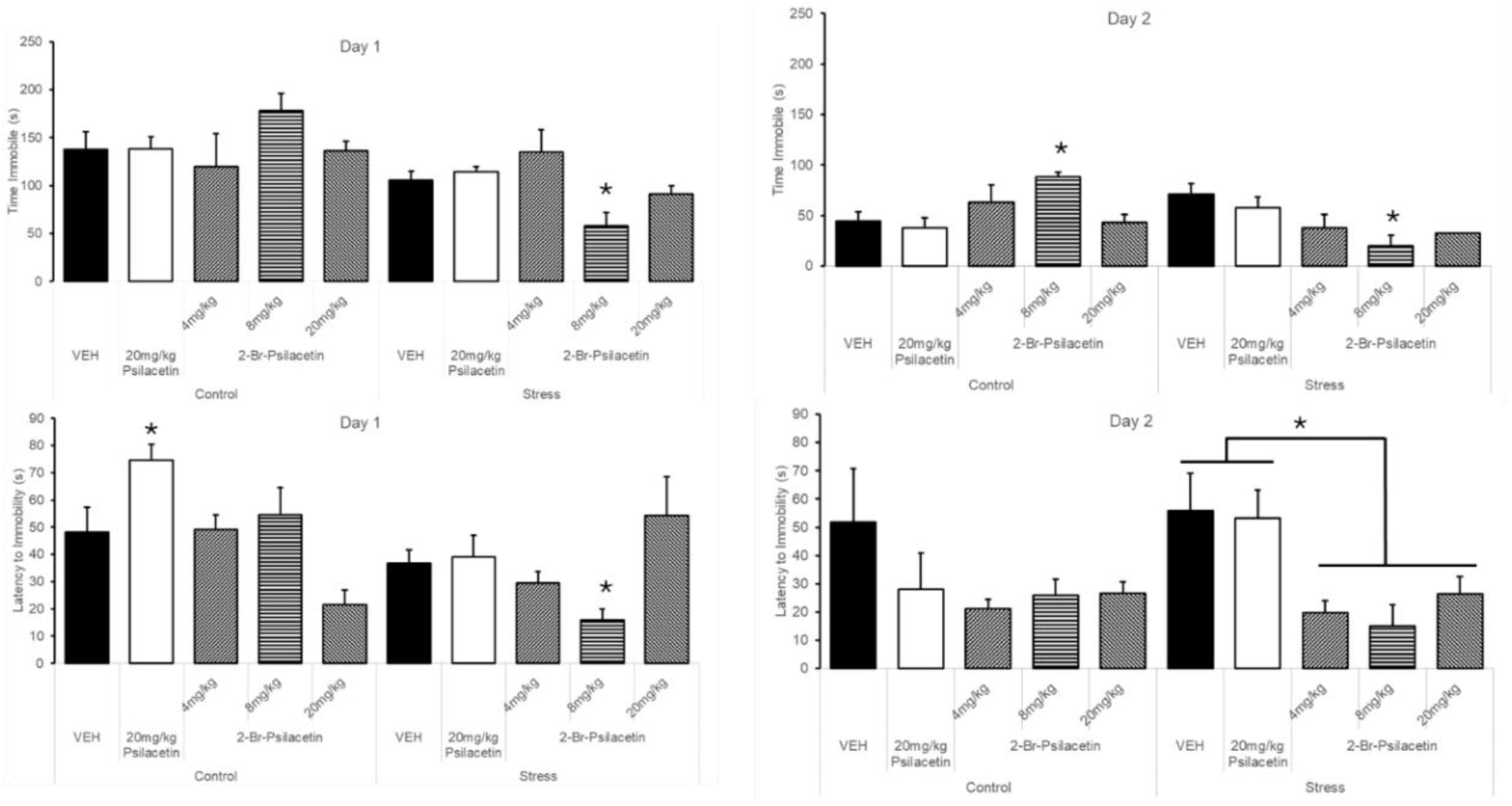
Porsolt forced swim test in C57BL/6 mice (A-B) Day 1 and (C-D) Day 2

For Day 2 latency to immobility, the interaction, F(4,45)=4.52, p=0.001 indicated that the groups administered 2-Br-psilacetin had a shorter latency to immobility compared to both VEH and psilacetin in the groups exposed to restraint stress (p<0.05). Together, these data suggest that the intermediate dose of 2-Br-psilacetin, 8 mg/kg, may have unique effects on stress-induced change in affect (**Figure 8C-D**).

### Social Behavior

In the assessment of social behavior, 8 mg/kg 2-Br-psilacetin increased positive social interactions in control mice, without altering behavior in the stress condition (**Figure 9**). Although 20 mg/kg 2-Br-psilacetin failed to increase social behavior in the younger mice, we did find a significant increase in exploratory rearing and a trend toward greater social interactions (p=0.055) in the aged mice (**Figure 9 inset)**.

**Figure 9.**
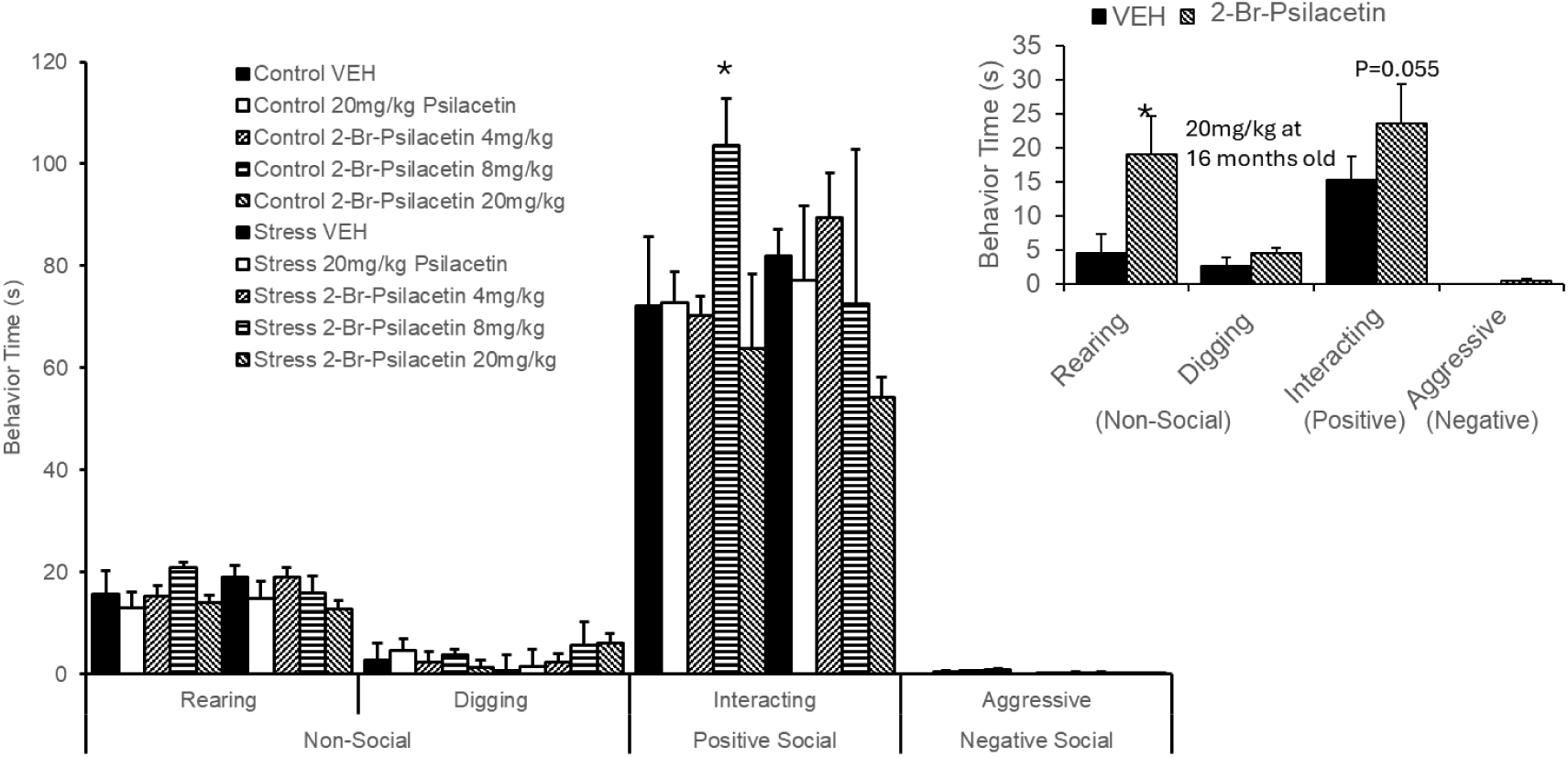
Social behavior in C57BL/6 mice, stressed and 16-month-old mice (inset)

### Startle Response

There was a main effect of drug treatment on startle at baseline (0 dB), F(4,54)=5.60, p=0.001, and at 120dB, f(4,54)=3.04, p=0.024, as well as an interaction of treatment by stress condition on 120 dB startle, F(4,54)=2.97, p=0.027. Psilacetin increased baseline activity in both the control and stress conditions compared to VEH (p<0.05), whereas 4 mg/kg 2-Br-psilacetin increased startle to 120dB compared to VEH only in the stress condition (p<0.05). Similar to head twitch, this suggests that the novel compound, 2-Br-psilacetin, differs from psilacetin in neurobehavioral outputs to affect and activity (**Figure S5A-B)**.

The median dose (8 mg/kg) of 2-Br-psilacetin seems to show a difference in neurobehavioral effect in stress-induced mice as compared to psilacetin. For instance, the Porsolt-forced swim test (**Figure 8**) showed a decrease in latency on day 1 and decrease in latency to immobility on day 2 for the stress-induced mice compared to vehicle dosing. A dose of 8 mg/kg, 2-Br-psilacetin increased positive social interactions in C57BL/6 mice but did not show a significant increase in social behavior at 20 mg/kg. Although the higher dose did improve social interactions in aged mice. It should be noted that at higher doses of 2-Br-psilacetin in mice showed signs of lethargy, as exhibited in the HTR paradigm, and (**Figure 5**) hypolocomotor activity could influence measures of freezing in fear conditioning at the 20 mg/kg dose. In contrast, 4 and 8 mg/kg 2-Br-psilacetin produce an increase in freezing to the cue, indicating an improvement in hippocampus-independent cued learning (**Figure 7A-B, Figures S4A-C).** These findings suggest that 2-Br-psilacetin exhibits dose-dependent behavioral effects, with moderate doses demonstrating promising prosocial and learning-related benefits. The overall results, demonstrate robust targeted effects of 2-Br-Psilacetin in these model systems at doses consistent with anticipated clinical exposures.^84^ Although the PFST has been discounted as a model of behavioral despair, it can instead be used as a behavioral response to stress. Immobility in this model is now widely interpreted as an adaptive stress-coping strategy. Within this new framework, it is less surprising that the FST has weak predictive validity for SSRIs, as it likely represents more of an internal processing model for the stress response.^85^ A final limitation is that no blockade studies were performed to evaluate how receptor activity influences the behavioral assays in mice.

## Conclusion

In conclusion, this work demonstrates that 2-position halogenation of tryptamine psychedelics yields compounds with markedly altered pharmacological and behavioral profiles compared to their parent molecules. The synthesized 2-halogenated derivatives of DMT and psilacetin/psilocin showed reduced affinity and partial agonism at 5-HT^2A^ and 5-HT^2B^ receptors, diminished HTRs, and improved cardiotoxicity profiles, while retaining high potency and efficacy at the CNS-selective 5-HT^6^ receptor. Among the compounds, 2-Br-psilocin and its prodrug 2-Br-psilacetin emerged as displaying 5-HT receptor promiscuity with minimal non-serotonergic activity, robust 5-HT^6^ activity, and minimal psychedelic-like behavior in mice. Behavioral studies further revealed dose-dependent effects on stress-related behaviors, cognition, and social interaction, suggesting therapeutic potential. Importantly, species-dependent differences at 5-HT^2A^ were observed, which warrant further investigation. Overall, these findings support 2-halogenated tryptamines as potentially promising psychedelic-inspired scaffolds for developing CNS therapeutics with modified psychoactivity that could also be efficacious to treat cognitive, affective, and stress-related disorders.

## Methods

### Synthesis of target compounds

All reactions were performed under argon in heat-dried glassware unless stated otherwise. Controlled temperature reactions were performed using a sand heat bath. Solvents were removed under reduced pressure using an Ika rotary evaporator. All solvents and reagents were purchased through Thermo Fisher and Oakwood Chemicals. Thin-layer chromatography (TLC) was performed using silica gel F^254^ glass plates. Detection of UV spots was performed using a UV lamp with both 254/365 nm wavelengths. Flash column chromatography was performed on glass columns using silica gel from Silicycle (230-400 mesh). Low-resolution mass spectra (LRMS) data were obtained on an Agilent LC-MS SQ 6120 (ESI). High-resolution mass spectra (HRMS) data were obtained on an Agilent 6540 series Q-TOF (ESI) and a Shimadzu LC-MS Triple TOF AB Sciex 5600 (ESI). Proton (^1^H and ^13^C NMR) were obtained using Bruker Neo with Prodigy (400 MHz) and Bruker Neo (600 MHz). Melting points were obtained on a Vernier Go Direct Melt Station MP apparatus GDX-MLT. The purity of samples tested biologically was determined to be ≥95% by UV.

### 2-Cl-*N*,*N*-dimethyltryptamine (2-(2-chloro-1*H*-indol-3-yl)-*N*,*N*-dimethylethan-1-amine)

To a solution of tryptamine hydrochloride (4.00 g, 20.3 mmol) in a 1:1 solution acetic acid and formic acid (65.1 mL) was added N-chlorosuccinimide (2.74 g, 20.5 mmol). The reaction was allowed to stir for 20 minutes after which the solvent was removed under reduced pressure. The crude oil was purified by column chromatography, eluting with 0-5% 7N ammonia/methanol in DCM. The colorless oil (1.95 g, 49.0 %) was immediately used in the next reaction.

To a solution of 2-chloro-tryptamine (0.650 g, 3.35 mmol) in ice cold methanol (22.3 mL) was added acetic acid (0.960 mL, 16.8 mmol) and sodium cyanoborohydride (0.442 g, 7.03 mmol). A solution of formaldehyde (0.710 mL, 8.71 mmol) in methanol (2.00 mL) was added dropwise at 0 °C and the solution was allowed to warm up to room temperature and stirred for 3 hours. The reaction was quenched with 1.95 mL of 20% aqueous sodium hydroxide. The methanol was removed under rotary evaporation and the product was extracted with chloroform three times. The combined organic layers were dried over sodium sulfate, filtered, and concentrated down. The residue was purified by column chromatography eluting with 0-5% 7N ammonia/methanol in DCM to give the final product as a tan solid (493 mg, 66.0%). HRMS (ESI-qToF) calc’d for C^12^H^15^ClN^2^, expected: 222.0924; found, 222.0927. Melting point: 92.4-94.4 ℃. ^1^H NMR (DMSO-d^6^, 400 MHz) δ 11.60 (br s, 1H), 7.47 (d, 1H, *J*=7.8 Hz), 7.27 (d, 1H, *J*=8.0 Hz), 7.09 (t, 1H, *J*=8.1 Hz), 7.01 (t, 1H, *J*=7.5 Hz), 2.78 (t, 2H, *J*=7.8 Hz), 2.43 (t, 2H, *J*=7.8 Hz), 2.20 (s, 6H); ^13^C NMR (DMSO-d^6^, 101 MHz) δ 134.5, 126.9, 121.5, 120.4, 119.2, 117.9, 110.8, 108.8, 59.0, 45.1, 21.8.

### 2-Br*-N,N*-dimethyltryptamine (2-(2-bromo-1H-indol-3-yl)-*N,N*-dimethylethan-1-amine)

Phenyltrimethylammonium tribromide (PTT) (1.01 g, 2.70 mmol) was dissolved in 12.3 mL anhydrous DCM and added dropwise to a solution of *N,N*-dimethyltryptamine (0.508 g, 2.70 mmol) in 24.3 mL anhydrous DCM. The reaction was stirred for 20 minutes at room temperature. Upon completion, the reaction was concentrated under reduced pressure and purified by flash column chromatography eluting with 0-8% iPrOH:DCM to give the product (0.375 g, 52.1%) as a tan solid. HRMS (ESI-qToF) calc’d for C^12^H^15^BrN^2^, expected: 266.0419; found, 266.0418. Melting point: 179.6-180.6 ℃. Free base: ^1^H NMR (DMSO-d^6^, 400 MHz) δ 11.61 (s, 1H), 7.49 (d, 1H, *J*=7.8 Hz), 7.28 (d, 1H, *J*=8.0 Hz), 7.08 (t, 1H, *J*=1.0 Hz), 7.01 (t, 1H, *J*=1.0 Hz), 2.77 (t, 2H, *J*=1.0 Hz), 2.42 (t, 2H, *J*=1.0 Hz), 2.21 (s, 6H); ^13^C NMR (DMSO-d^6^, 151 MHz) δ 136.6, 127.6, 121.9, 119.6, 118.3, 112.4, 111.2, 108.9, 59.6, 45.6, 23.2. HBr salt: ^1^H NMR (DMSO-d_6_, 400 MHz) δ 11.84 (s, 1H), 9.71 (br s, 1H), 7.64 (d, 1H, *J*=7.8 Hz), 7.32 (d, 1H, *J*=8.0 Hz), 7.13 (t, 1H, *J*=1.0 Hz), 7.06 (t, 1H, *J*=7.4 Hz), 3.2-3.3 (m, 2H), 3.0-3.1 (m, 2H), 2.89 (br s, 6H); ^13^C NMR (DMSO-d_6_, 101 MHz) δ 136.7, 127.1, 122.4, 119.9, 118.4, 111.5, 109.8, 108.7, 56.2, 42.6, 20.5.

### Psilacetin (3-(2-(dimethylamino)ethyl)-1H-indol-4-yl acetate)

Psilocin (1.00 g, 4.89 mmol) was dissolved in anhydrous DCM. At 0 °C, a solution of acetyl chloride (384 µl, 5.38 mmol) in 5.38 ml of anhydrous DCM was added dropwise, followed by triethylamine (751 µl, 5.38 mmol). The reaction mixture was allowed to warm up to room temperature and stirred for three hours. The solvent was removed under reduced concentration and the residue was directly adsorbed onto silica gel. The crude material was purified by column chromatography eluting with 0-15% iPrOH in DCM to give a tan solid (1.18 g, 98.0%). ^1^H NMR (CHLOROFORM-d, 400 MHz) δ 8.62 (br s, 1H), 7.1-7.2 (m, 2H), 6.85 (s, 1H), 6.79 (d, 1H, *J*=7.1 Hz), 2.92 (t, 2H, *J*=7.6 Hz), 2.62 (t, 2H, *J*=8.5 Hz), 2.40 (s, 3H), 2.34 (s, 6H); ^13^C NMR (CHLOROFORM-d, 101 MHz) δ 170.3, 144.1, 138.8, 122.7, 122.0, 119.9, 112.7, 112.2, 109.4, 61.0, 45.6, 24.9, 21.3.

### 2-Br-psilacetin (2-bromo-3-(2-(dimethylamino)ethyl)-1*H*-indol-4-yl acetate)

Phenyltrimethylammonium tribromide (PTT) (1.51 g, 4.02 mmol) was dissolved in 18.0 mL anhydrous DCM and added dropwise to a solution of 4-Acetoxy-*N,N-*dimethyltryptamine (0.900 g, 3.66 mmol) in 36.0 mL anhydrous DCM. The reaction was stirred for 20 minutes at room temperature. Upon completion, the reaction was concentrated under reduced pressure and purified by flash column chromatography eluting with 0-8% iPrOH in DCM to give the product (0.980 g, 39.3 %) as a grey solid (HBr salt). HRMS (ESI-qToF) calc’d for C_14_H_17_BrN_2_O_2_, expected: 324.0473; found, 324.0473. Melting point: 167.9-170.1 ℃. Free base: ^1^H NMR (DMSO-d_6_, 600 MHz) δ 11.89 (br s, 1H), 7.17 (d, 1H, *J*=8.2 Hz), 7.07 (t, 1H, *J*=8.0 Hz), 6.73 (d, 1H, *J*=7.8 Hz), 2.71 (t, 2H, *J*=8.0 Hz), 2.35 (s, 3H), 2.31 (t, 2H, *J*=8.4 Hz), 2.20 (s, 6H); ^13^C NMR (DMSO-d_6_, 151 MHz) δ 169.9, 142.9, 138.6, 122.1, 119.9, 113.1, 110.6, 110.3, 109.3, 60.5, 45.6, 24.6, 21.2. HBr salt: ^1^H NMR (DMSO-d_6_, 600 MHz) δ 12.16 (s, 1H), 9.78 (br s, 1H), 7.22 (d, 1H, *J*=8.2 Hz), 7.11 (t, 1H, *J*=8.0 Hz), 6.82 (d, 1H, *J*=7.6 Hz), 3.15 (t, 2H, *J*=1.0 Hz), 3.02 (t, 2H, *J*=1.0 Hz), 2.88 (s, 6H), 2.42 (s, 3H); ^13^C NMR (DMSO-d_6_, 151 MHz) δ 170.0, 142.7, 138.6, 122.5, 119.5, 113.3, 111.4, 109.4, 107.0, 57.5, 42.9, 21.6, 21.4.

### 2-Br-psilocin (2-bromo-3-(2-(dimethylamino)ethyl)-1H-indol-4-ol)

To an aqueous solution of 1M hydrobromic acid (5.00 mL) was added 2-Br-psilacetin (0.100 g, 0.309 mmol) under inert atmosphere. The reaction was heated to 70 °C for two hours, after which saturated sodium bicarbonate was added to adjust the solution to pH 7. The product was extracted with ethyl acetate (3 x 50.0 mL). The combined organic layer was dried over sodium sulfate, filtered, and concentrated down. Purification was done by preparative thin layer chromatography eluting with 10% methanol in DCM to give the final product as a white solid (0.0420 g, 48.0%). HRMS (ESI-qToF) calc’d for C_12_H_15_BrN_2_O, expected: 282.0368; found, 282.0373. ^1^H NMR (METHANOL-d_4_, 600 MHz) δ 5.30 (t, 1H, *J*=6.7 Hz), 5.18 (d, 1H, *J*=6.9 Hz), 4.78 (d, 1H, *J*=6.5 Hz), 1.44 (t, 2H, *J*=5.8 Hz), 1.12 (t, 2H, *J*=5.9 Hz), 0.80 (s, 6H); ^13^C NMR (METHANOL-d_4_, 600 MHz) δ 151.9, 151.8, 140.1, 123.8, 123.7, 118.4, 112.5, 107.6, 105.6, 103.5, 61.4, 45.2, 24.8.

### 2-Cl-psilocin (2-chloro-3-(2-(dimethylamino)ethyl)-1H-indol-4-ol)

To a solution of 2 M hydrochloric acid in dioxane (4.26 mL, 8.52 mmol) was added 2-Br-psilacetin (0.200 g, 0.620 mmol) under inert atmosphere. The reaction was left to stir at room temperature for 18 hours, after which saturated sodium bicarbonate was added to adjust the solution to pH 7. The product was extracted with ethyl acetate (3 x 50.0 mL). The combined organic layer was dried over sodium sulfate, filtered, and concentrated down. The product was purified by flash column chromatography, eluting with 0-3% methanol in dichloromethane and collected as a white solid (0.0780 g, 53.1%). HRMS (ESI-triple TOF) calc’d for C_12_H_15_ClN_2_O, expected: 238.0875; found, 238.0888. ^1^H NMR (METHANOL-d4, 600 MHz) δ 6.88 (t, 1H, J=1.0 Hz), 6.74 (br d, 1H, J=8.4 Hz), 6.37 (br d, 1H, J=7.8 Hz), 3.0-3.0 (m, 2H), 2.7-2.7 (m, 2H), 2.38 (s, 6H); ^13^C NMR (METHANOL-d4, 600 MHz) δ 152.0, 138.4, 123.9, 120.3, 118.1, 108.7, 105.8, 103.6, 61.1, 45.0, 23.4.

### Pharmacological Target Screening

Receptor binding and functional screening assays in transfected cells were carried out as described in detail.^86^

### BBB PAMPA assays

The Blood-brain barrier (BBB) parallel artificial membrane permeability assay (PAMPA) procedure was realized by using a method developed by Pion, Inc. All liquid handling steps were performed on the TECAN Freedom EVO150 robot and analyzed by the Pion PAMPA Evolution Software. BBB PAMPA included brain the sink buffer (BSB), lipid solution (BBB-1) and Stirwell^TM^ PAMPA Sandwich plate preloaded with magnetic stirring disks. 4 μLof lipid solution was transferred in the acceptor well, followed by the addition of 200 μL of BSB (pH 7.4). 180 μL of diluted test compounds (50-250 μM in system buffer at pH 7.4 from a 10 mM DMSO solution) were added to the donor wells. The PAMPA sandwich plate was assembled, placed on the Gut-BoxTM and stirred with 60 μm Aqueous Boundary Layer (ABL) settings for 1 hour incubation. The distribution of the compounds in the donor and acceptor buffers (150 μL aliquot) was determined by UV spectra measurement from 250 to 498 nm using the TECAN infinite M-1000 pro microplate reader. Permeability (*P*app, 10-6cm/s) of each compound was calculated by the Pion PAMPA evolution software. The assay was performed in triplicate.

### 5-HT Receptor G protein dissociation BRET assays

All BRET assays were conducted using BRET^2^ with *R*luc8 and GFP tagged proteins expressed in HEK293T cells (ATCC CRL-11268). Cell lines were subcultured in high-glucose DMEM (VWR) supplemented with 10% FBS (Life Technologies). All 5-HT receptor and G protein constructs, including mouse and rat 5-HT receptor constructs were made and codon-optimized for this study. Approximately 48 hours before assays, cells were transfected in 1:1:1:1 ratios of 5-HT receptor/Gα-*R*luc8/β/GFP-γ constructs for G protein dissociation assays. All transfections were prepared in Opti-MEM (Invitrogen) and used a 3:1 ratio of TransIT-2020 (Mirus) μL/μg total DNA. Next day, cells were plated at an approximate density of 40-50,000 cells per well into poly-L-lysine-coated 96-well white assay plates (Grenier Bio-One). On the day of the assay, plates were decanted and 60 μL of drug buffer per well (HBSS, 20mM HEPES, pH 7.4) was added using a Multidrop (ThermoFisher Scientific), and plates were allowed to equilibrate at 37°C in a humidified incubator approximately 15 min before receiving stimulation. Compound dilutions (3X) were performed in McCorvy buffer (HBSS, 20mM HEPES, pH7.4, supplemented with 0.3% BSA fatty acid free (GoldBio), and 0.03% ascorbic acid). Next, plates were incubated with compounds at 37°C in a humidified incubator for total time of 60 min before reading. Approximately 15 minutes before reading, plates were removed from the incubator and coelenterazine 400a (5 μM final concentration; Nanolight Technology) was added per well. Immediately after 60 min incubation, plates were read at 410-480 nm (*R*Luc8) and 515-530 nm (GFP) emission filters for 1.0 second per well using either a PheraStarFSX or ClarioStarPlus (BMGLabTech, Cary NC). The BRET ratios of GFP/*R*luc8 luminescence were calculated per well and were plotted as a function of compound concentration using GraphPad Prism 11 (Graphpad Software Inc., San Diego, CA). Data were normalized to percent 5-HT response, which 5-HT control was present on every plate for within experiment comparisons. Data was analyzed using nonlinear regression “log(agonist) vs. response” to yield Emax and EC_50_ parameter estimates.

### Gq-mediated calcium flux assays

Stably expressing 5-HT_2A/2B/2C_ receptor Flp-In 293 T-Rex cell lines (Invitrogen) were used for calcium flux assays as described previously.^87^ Serotonin 5-HT_2A/2B/2C_ receptor expression was induced by addition of tetracycline (2 μg/mL) and seeded into 384-well poly-L-lysine-coated black plates at a density of approximately 7,000-10,000 cells/well in DMEM containing 1% dialyzed FBS. On the day of the assay, plating media was decanted and cells were incubated with Fluo-4 Direct dye (Invitrogen, 20 μl/well) for approximately 60 min at 37°C, which was reconstituted in drug buffer (HBSS, 20mM HEPES, pH 7.4) containing 2.5 mM of probenecid. After dye load, cells were allowed to equilibrate to room temperature for 15 min and then placed in a FLIPR^TETRA^ fluorescence imaging plate reader (Molecular Devices). Compound dilutions were prepared at a 5X final concentration in McCorvy buffer (HBSS, 20mM HEPES, pH7.4, supplemented with 0.3% BSA fatty acid free (GoldBio), and 0.03% ascorbic acid). Compounds were aliquoted into 384-well plates and placed in the FLIPR^TETRA^ for stimulation. Fluorescence for the FLIPR^TETRA^ was programmed to read baseline fluorescence for 10 s (1 read/s), and afterward, 5 μl of compound per well was added and read for a total of 120 s (1 read/s). Fluorescence in each well was normalized to the average of the first 10 reads for baseline fluorescence, and then area under the curve (AUC) was calculated. AUC was plotted as a function of compound concentration, and data were normalized to percent 5-HT stimulation, which was present on every plate to control for within experiment error. Data were plotted, and nonlinear regression was performed using “log(agonist) vs. response” in Graphpad Prism 11 to yield Emax and EC_50_ parameter estimates.

### Gi/o-mediated and Gs-mediated cAMP measurements

HEK293T cells (ATCC CRL-11268; mycoplasma-free) were co-transfected in 1:1 ratio of codon-optimized 5-HT receptors as described above and GloSensor-22F plasmid (Promega) in 10% dFBS. On the following day, cells were plated into poly-L-lysine-coated-384-well white assay plates (Grenier Bio-One) at a density of approximately 10,000 cells per well. After approximately 48 hrs post-transfection, plates were decanted and 20uL/well of drug buffer (1x HBSS, 20mM HEPES, pH 7.4) containing 4 mM D-luciferin (sodium salt; GoldBio) was added using a Multidrop (ThermoFisher Scientific). Cells were allowed to equilibrate for approximately 15 minutes at room temperature and then compounds serially diluted in McCorvy buffer (drug buffer containing 0.3% BSA fatty-acid free and 0.03% ascorbic acid). Automated compound dispensing was performed by a FLIPR^TETRA^ (Molecular Devices). For Gi/o-coupled receptors tested (5-HT_1A_, 5-HT_1B_, 5-HT_1F_), cells were incubated with drugs for exactly 15 minutes at room temperature and then stimulated with 0.2 µM isoproterenol to stimulated cAMP production. After 15 minutes post isoproterenol stimulation, plates were then read for luminescence (LCPS) for a total of 30 min drug incubation on a Microbeta Trilux (PerkinElmer). For Gs-coupled receptors tested (5-HT_6_, 5-HT_7_), cells were incubated approximately 30 min at room temperature and read for luminescence (LCPS). Data were plotted, and nonlinear regression was performed using “log(agonist) vs. response” in GraphPad Prism 11 to yield Emax and EC_50_ parameter estimates.

### Mouse Brain Competition Binding Assays

Mouse brain competition binding studies were conducted as previously described without modifications.^34^ Briefly, the experiments utilized C57BL/6 mouse brain tissue (BioIVT, Westbury, NY, USA) to assess ability of the compounds to compete for [^3^H]8-OH-DPAT binding to 5-HT_1A_ receptors (0.5 nM) or [^3^H]M100907 binding to 5-HT_2A_ receptors (1 nM). Radioligands were procured from PerkinElmer (Boston, MA, USA). Nonspecific binding was determined using 10 μM serotonin for 5-HT_1A_ and ketanserin for 5-HT_2A_ binding. Dissociation constants used for one site - fit K_i_ determinations in GraphPad Prism were derived from previously published studies using mouse brain (K_d_ = 1.03 and 0.35 nM respectively). All data represent 3 runs performed in triplicate for each value.

### Head Twitch Response Assay

Briefly, groups of 24 C57BL/6J mice (12 male, 12 female; 8 weeks; Jackson #000664) were housed 4/cage at NIDA IRP (Baltimore) on a 12/12 h light/dark cycle (lights on 6–7 AM). After 1–2 weeks acclimation, mice were briefly immobilized with isoflurane and implanted subcutaneously with TP500 transponders (upper back; Avidity) for noninvasive temperature recording (DAS-8027 IUS). Mice were then single-housed post-implantation.

For single drug assays, mice received subcutaneous test drug (0.03–30 mg/kg) or vehicle (saline), and behavior was video-recorded for 30 min (GoPro cameras; 1080P, 120 fps). For antagonist studies, the protocol was the same except mice received injection of 2-Br-psilactein (10 mg/kg s.c.) 10 min prior to a known HTR active dose of the 5-HT2 agonist DOI (0.5 mg/kg s.c.). Baseline temperature was recorded after brief chamber acclimation and compared to post session body temperature readings to obtain temperature change values used. Motor activity (distance traveled) was determined using the head pose-estimation key point further described below.

For video analyses used to determine HTR counts and distance traveled, a DeepLabCut (DLC) pose-estimation model (videos downscaled to 640×480, 120 fps via FFmpeg) was trained as recently described.^88^ A SimBA classifier (13 body parts; 32 features; pixels/mm set from chamber width; outlier correction skipped) was trained on historical HTR videos not used in DLC training and achieved precision 95.93, recall 93.74, F1 score 94.82 on 38 validation videos (879 HTRs). Hardware/software used: Dell Precision T5820 (Xeon W-2223 3.6 GHz, 32 GB RAM), Windows 11 Pro, NVIDIA RTX A100 GPU; FFmpeg; DLC v2.3.10; SimBA v2.2.4.

For data analyses, HTR dose-response curves were visualized with biphasic nonlinear fits, while four-parameter fits were used for motor activity and body temperature change data. HTR potencies were derived from the ascending limb of dose-response curves and maximal counts (efficacies) were the peak counts observed for each drug. Group comparisons were made via Welch’s ANOVA with Dunnett’s T3 vs vehicle. GraphPad Prism was used for all statistical analyses and figure creation for HTR experiments, with alpha set at 0.05 for all comparisons.

### *in vivo* Mouse studies

In this study, 8–12-week-old, male, C57BL/6J mice (Jackson Laboratory) were tested. For all studies, half of the mice were placed in restraint (50 mL conical tubes with air holes and a hole for the tail in the cap) to induce stress. Control mice remained in their housing for the two-hour period. At the end of this period, , all mice were treated daily with either vehicle, 20 mg/kg psilacetin, or 4-20 mg/kg 2-Br-psilacetin. Immediately after injections were completed, mice went through a behavioral battery, with one behavior assessed per day. Mice to be tested were placed onto a cart for at least 30 min to habituate them to the testing environment. Behavior was run in a room separate from all other animal housing. All behavioral equipment was cleaned in between mice, and after each assay was concluded. All relevant data and videos of assays were collected from the Any Maze software and subsequently scored manually by blinded researchers.

### Fear Conditioning

Contextual and cued fear conditioning evaluate fear-based learning and memory. Animals were tested as previously described.^89^ Briefly, animals were trained in the fear conditioning apparatus with two 30-s acoustic conditioned stimuli (CS), each paired with an un-conditioned stimulus (US): a 0.57 mA foot shock. To assess contextual fear conditioning, mice were placed back into the same context for 3 min, 24 h following the CS-US pairings. To assess cued fear conditioning, mice were placed in a novel context. Tests were scored by a researcher blinded to the experimental conditions, and the animal was freezing in the absence of movement except for respiration.

### Porsolt Forced Swim

The forced swim behavioral assay is performed to assess antidepressant efficacy and to evaluate a subject’s depressive state at the time of assay.^90^ When mice are placed in the water, escape-related movement is scored, with more movement aimed at escape demonstrating a less-depressive state on day one, whereas the animals have a shorter swim time on day two, demonstrating memory of the coping strategy used previously (floating).

The swim beaker, a weighted, opaque acrylic cup, was filled half full of room temperature water and placed in the open field apparatus under the video camera. Mice were gently placed in the water and allowed to swim or float for 6 min, then the mouse was removed, returned to its home cage. The water in the cup changed every 2-3 mice. Scores were collected for the latency to the first stop (the first time the mouse stopped swimming), and the total duration of the test that the mouse spent floating or not swimming.

### Socialization

The socialization assay is performed to assess both positive and negative social behaviors, as most antidepressant treatments boost the former without affecting the latter. The treated mouse is placed in an open field with an unfamiliar mouse of the same sex and similar age for 10 min. Curious behaviors towards the unfamiliar mouse are considered positive, avoidant or aggressive behaviors are considered negative, whereas grooming or solo exploration are categorized as neutral. Herein, the treated mice were placed in an open field Plexiglas box,18”x18”x16”, then an unfamiliar mouse was introduced, and the behavior of both mice was recorded and manually scored for 10 min.

### Startle Response

Startle responding was assessed as a measure of baseline stress reactivity, using an auditory tone. Animals were placed in a clear, cylindrical restrainer (Panlab, Barcelona, Spain) inside a sound-attenuation chamber. Animals were acclimated for 5 min in the presence of 65 dB white noise then were presented with two trial types in a pseudo-random order with an inter-trial interval of 10–20 s: a 40 ms, 120 dB sound burst (startle) or no stimulus (baseline). Startle responses were measured as force and latency to response using an accelerometer located beneath the restraint platform.

### Data Analysis

Data analysis was performed in Image Lab and IBM SPSS Statistics v27. For behavioral assays, multivariate ANOVAs were run with two independent variables, restraint and treatment condition; the latter was coded as a single variable with 5 levels: Veh (negative control), 20 mg/kg psilacetin (positive control), and 4-20 mg/kg 2-Br-psilacetin, followed by Tukey post hoc tests to assess differences between individual groups. Significance was assessed at p<0.05.

## Supporting information

SI_Serotonergic Polypharmacology of 2-Halogenated Tryptamines

## Acknowledgements

We would like to thank Dr. Laurent Calcul at the USF CPAS facility, Dr. Peidong Zhao at the USF NMR core facility, and the Florida High Tech Corridor for project fund matching. This study was partially supported by NIGMS R35 GM133421 (J.D.M), NIDA R01 DA061433 (J.D.M), the NIH/NIAAA Award #R44AA031925, and the Intramural Research Program of the National Institute on Drug Abuse (NIDA IRP; ZIA DA000522 to M.H.B). HTR experiments were also supported in part by a collaborative research and development agreement between NIDA IRP and Psilera. K_i_ determination and receptor binding profiles for 2-halogenated tryptamines were generously provided by the National Institute of Mental Health’s Psychoactive Drug Screening Program, Contract #75N95023C00021 (NIMH PDSP). The NIMH PDSP is Directed by Bryan L. Roth MD, PhD at the University of North Carolina at Chapel Hill and Project Officer Jamie Driscoll at NIMH, Bethesda MD, USA.

## Author Information

## Authors

Elena Bray – Psilera, Inc. Tampa, FL, US 33612

Jude Bayyat – Byrd Alzheimer’s Institute and Department of Molecular Medicine, University of South Florida, Tampa, FL, US 33612

Grant C. Glatfelter – Designer Drug Research Unit, National Institute on Drug Abuse, Baltimore, MD, US 21224

Alex Leake – Byrd Alzheimer’s Institute and Department of Molecular Medicine, University of South Florida, Tampa, FL, US 33612

Emilia Buitrago – Byrd Alzheimer’s Institute and Department of Molecular Medicine, University of South Florida, Tampa, FL, US 33612

Hubert Nguyen – Byrd Alzheimer’s Institute and Department of Molecular Medicine, University of South Florida, Tampa, FL, US 33612

Alexander D. Maitland – Designer Drug Research Unit, National Institute on Drug Abuse, Baltimore, MD, US 21224

John Partilla – Designer Drug Research Unit, National Institute on Drug Abuse, Baltimore, MD, US 21224

John L. McKee – Department of Cell Biology, Neurobiology, and Anatomy, Medical College of Wisconsin, Milwaukee, WI, US 53226

Natalie G. Cavalco – Department of Cell Biology, Neurobiology, and Anatomy, Medical College of Wisconsin, Milwaukee, WI, US 53226

Serene S. Schalk – Department of Cell Biology, Neurobiology, and Anatomy, Medical College of Wisconsin, Milwaukee, WI, US 53226

Josie C. Lammers – Department of Cell Biology, Neurobiology, and Anatomy, Medical College of Wisconsin, Milwaukee, WI, US 53226

Michael H. Baumann – Designer Drug Research Unit, National Institute on Drug Abuse, Baltimore, MD, US 21224

John D. McCorvy – Department of Cell Biology, Neurobiology, and Anatomy, Department of Pharmacology and Toxicology, Neuroscience Research Center, Cancer Center, and Cardiovascular Research Center, Medical College of Wisconsin, Milwaukee, WI, US 53226

James W. Leahy – Department of Molecular Medicine, Department of Chemistry, University of South Florida, Tampa, FL, US 33612

Danielle Gulick – Byrd Alzheimer’s Institute and Department of Molecular Medicine, University of South Florida, Tampa, FL, US 33612

Christopher G. Witowski - Psilera, Inc. Tampa, FL, US 33612

Jacqueline L. von Salm - Psilera, Inc. Tampa, FL, US 33612

## Author Contributions

J.Y. and E.B. led the design and synthesis of the test compounds. J.W.L. acted as supervisor for J.Y. regarding synthesis. J.Y. wrote the manuscript with major contributions from G.C.G., J.D.M, and D.G., as well as major edits from G.C.G., J.DM., J.vS., D.G., E.B., M.H.B., and C.W. E.B. wrote the Supplementary Information with edits from J.Y. 5-HT receptor functional assays were performed by J.L.M, S.S.S., and N.G.C. under J.D.M supervision. Expression constructs were made by J.C.L under J.D.M supervision. Mouse brain binding studies and head twitch response were done by G.C.G., J.P., and A.D.M. and behavioral studies were conducted by D.G. and J.B. All authors approved the final version of the manuscript.

