## Supplementary material for "Serotonergic Polypharmacology of 2-Halogenated Tryptamines": SI_Serotonergic Polypharmacology of 2-Halogenated Tryptamines

[1]Psilera, Inc. Tampa, FL; [2]Byrd Alzheimer's Institute, University of South Florida Health, Tampa, FL; [3]Department of Molecular Medicine, University of South Florida Health, Tampa, FL; [4]Designer Drug Research Unit, National Institute on Drug Abuse, Baltimore, MD; [5]Department of Cell Biology, Neurobiology, and Anatomy, Medical College of Wisconsin, Milwaukee, WI, USA; [6]Department of Pharmacology and Toxicology, Neuroscience Research Center, Cancer Center, and Cardiovascular Research Center, Medical College of Wisconsin, Milwaukee, WI, USA; [7]Department of Chemistry, University of South Florida, Tampa, FL, USA

**Graphical Abstract**

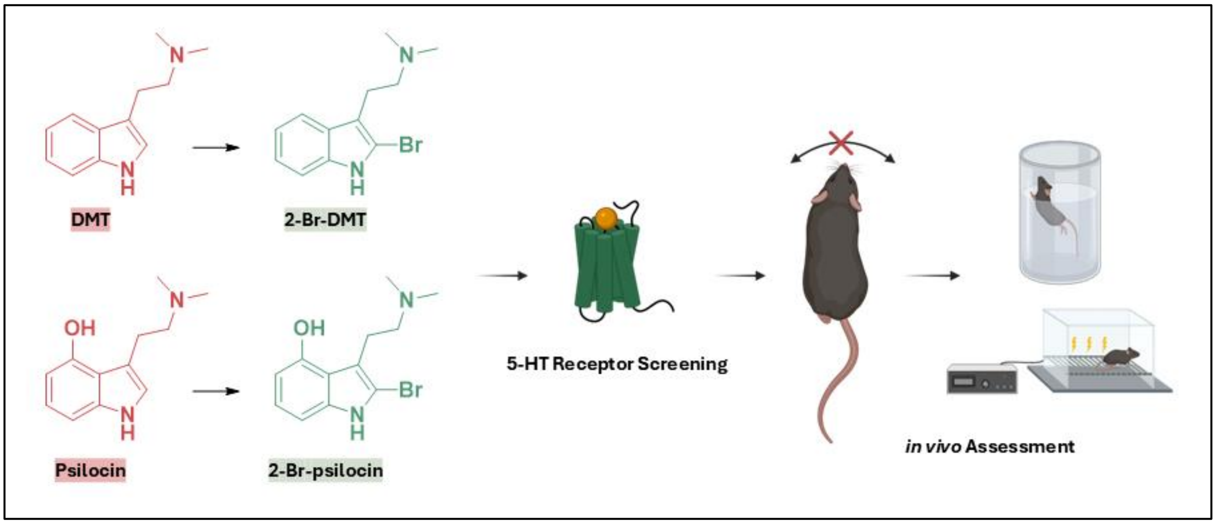

**Table of Contents**

Synthesis Schemes for 2-halogenated tryptamines …………………………..……..... pg. S3
*In Vitro* Pharmacology …………………………………………….……………….…..... pg. S4
Mouse brain binding affinity ………………………….………………………...……..... pg. S5
Second messenger assay for 2-X-tryptamines …..……….…..……….…..….……….….. pg. S6
BBB Permeability ………………………………………………………….……………... pg. S6-S7
PK Parameters of 2-Br-psilacetin ……………………………..……………………… pg. S7
Brain/Plasma exposure in rats ………………………………………………..………. pg. S7
Fear conditioning ……………………………………………………….…………….………. pg. S8
Startle response baseline and 120 dB startle …………….…………….……….………. pg. S8
NMR Spectra ………………………………………………………….……………….. pg. S10-S16
HRMS Spectra ..…..……….…………..……………..……….……………………… pg. S17-S18
UV Purity of Test Compounds ………………………………………………………. pg. S19-S22
References ……………………………………………………………………….……...… pg S23

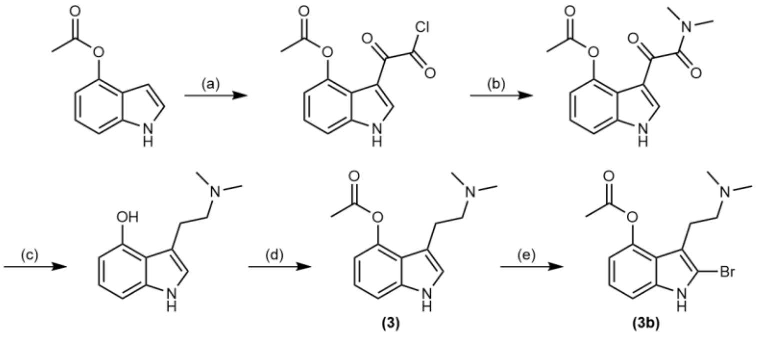


**Scheme S1:** Synthesis of 2-Br-psilacetin; (a) oxalyl chloride, diethyl ether, 0 °C, 12h; (b) $HN(CH_3)_2$, $NEt_3$, 2-Me-THF, 0 °C - rt, 12h, **97%** over two steps; (c) $LiAlH_4$, 2-Me-THF, reflux, 12h, **62%**; (d) Acetyl chloride, DCM, 0 °C - rt, 3h, **98%**; (e) $PhMe_3NBr_3$, DCM, 20 min; **83%**

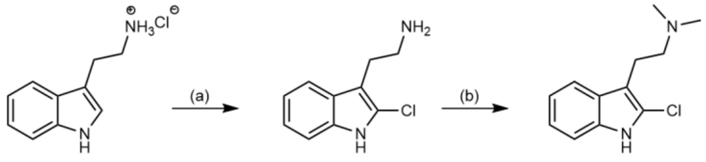


(a) NCS, AcOH/$HCO_2H$, 20 min, **58%**; (b) AcOH, $NaCNBH_3$,$H_2CO$, MeOH, 0 °C - rt, **52%**

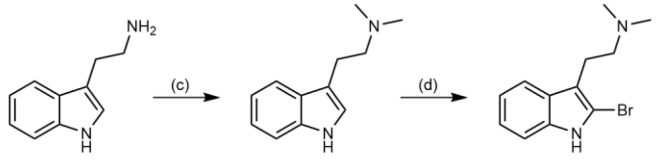


(c) AcOH, $NaCNBH_3$,$H_2CO$, MeOH, 0 °C - rt, 3h, **90%**; (d) $PhMe_3NBr_3$, DCM, 20 min, **52%**

**Scheme S2:** Synthesis of 2-X-DMT derivatives; (a) AcOH, $NaCNBH_3$, $H_2CO$, MeOH, 0 °C - rt, 3h, **90%**; (b) $PhMe_3NBr_3$, DCM, 20 min, **52%**; (c) NCS, AcOH/$HCO_2H$, 20 min, **58%**; (d) AcOH, $NaCNBH_3$, $H_2CO$, MeOH, 0 °C - rt, **52%**

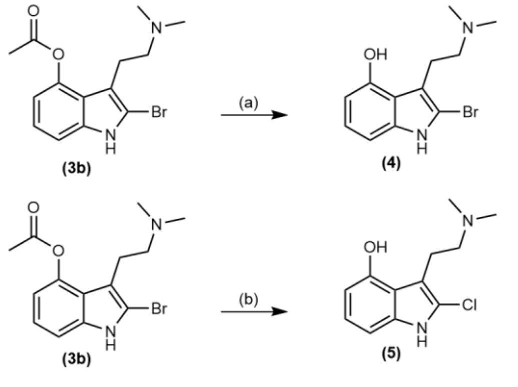


**Scheme S3:** Synthesis of 2-X-psilocin derivatives; (a) 1 M aq. HBr, 70 °C, 2h, **48%**; (b) 1M HCl in dioxane, rt, 18h, **53%**

### *In Vitro* Pharmacology: Cellular binding Assays

2-Cl-psilocin and 2-Br-psilocin cellular assays were tested by Eurofins Cerep, Celle l'Evescault, France. Assays were run using their standard protocols for each receptor: 5-$HT_{1A}$,[1] 5-$HT_{2A}$,[2] 5-$HT_{2B}$,[3] 5-$HT_{2C}$,[4] 5-$HT_6$,[5] and 5-$HT_7$,[6].

2-Cl-DMT, 2-Br-DMT, 2-Br-psilacetin, DMT and psilacetin cellular assays were generously screened by the National Institute of Mental Health (NIMH) Psychoactive Drug Screening Program (PDSP) using their standard conditions for each receptor.[7]

***Table S1. Radioligand binding assays, $K_i$ (nM) at serotonin receptors[a]***

| Compound | 5-$HT_{1A}$ | 5-$HT_{2A}$ | 5-$HT_{2B}$ | 5-$HT_{2C}$ | 5-$HT_6$ | 5-$HT_{7a}$ |
|---|---|---|---|---|---|---|
| ***2-Cl-psilocin*** | 550 | 74 | 25 | 76 | 35 | 47 |
| ***2-Br-psilocin*** | 980 | 97 | 82 | 370 | 29 | 99 |
| ***Reference*** | 0.5 | 0.6 | 2.4 | 0.5 | 1.8 | 2.3 |

***Table S1:*** *Binding affinities of 2-X-psilocin derivatives at 5-HT receptors; [a]Radioligand and reference control compound for each 5-HT receptor: 5-$HT_{1A}$ = [$^3$H]-8-OH-DPAT vs 8-OH-DPAT, 5-$HT_{2A}$ = [$^3$H]-ketanserin vs. ketanserin, 5-$HT_{2B}$ = [$^3$H]-mesulergine vs SB206553, 5-$HT_{2C}$ = [$^3$H]-mesulergine vs. RS 102221, 5-$HT_6$ = [$^3$H]-LSD vs. serotonin, 5-$HT_{7a}$ = [$^3$H]-LSD vs. serotonin.*

***Table S2. Radioligand binding assays, Ki (nM) at non-5-HT receptors and targets[a]***

| Compound | $H_1$ | $H_2$ | KOR | NR2B | $Sigma_1$ | $Sigma_2$ | $alpha_{2A}$ | $alpha_{2B}$ | $alpha_{2C}$ | SERT |
|---|---|---|---|---|---|---|---|---|---|---|
| ***2-Br-DMT*** | 779.47 | 752.84 | 3535.9 | 5011.51 | 239.33 | 2458.67 | 670.35 | 259.84 | 4434.04 | - |
| ***2-Cl-DMT*** | 1414.49 | 678.42 | - | - | 637.68 | - | 866.16 | 759.1 | 8570.38 | - |
| ***DMT*** | 158.82 | 3645.02 | 4371.19 | - | 229.99 | 776.6 | 2034.23 | 621.44 | 4891.03 | - |
| ***2-Br-Psilacetin*** | - | - | 1034.43 | - | - | - | 980.62 | 801.49 | - | - |
| ***Psilacetin*** | 3508.33 | - | - | - | - | - | 4762.12 | 2991.58 | - | - |
| ***Reference*** | 2.21 | 3.25 | 1.84 | 13.47 | 3.78 | 37.9 | 4.64 | 7.54 | 68.36 | 24.37 |

***Table S2:*** *Binding affinities (Ki, nM) of test compounds at aminergic CNS receptors; [a]Radioligand and reference control compound for each receptor: $H_1$ = [$^3$H]-pyrilamine vs chlorpheniramine; $H_2$ = [$^{125}$I]-iodoaminpotentidine vs ORG-5222; KOR = [$^3$H]-U69593 vs salvinorin A; NR2B = [$^3$H]-ifenprodil vs ifenprodil; $Sigma_1$ = [$^3$H]-pentazocine vs haloperidol; $Sigma_2$ = [$^3$H]-DTG vs haloperidol; $alpha_{2A}$, $alpha_{2C}$ = [$^3$H]-rauwolscine vs oxymetazoline; $alpha_{2B}$ = [$^3$H]-rauwolscine vs yohimbe, SERT = [$^3$H]-citalopram vs amitriptyline*

***Table S3. Radioligand binding assays, Ki (nM) at non-5-HT receptors and targets[a]***

| Compound | $D_1$ | $D_2$ | $D_3$ | $D_4$ | $M_4$ | $M_5$ | hERG |
|---|---|---|---|---|---|---|---|
| ***2-Br-DMT*** | 2305.69 | 2987.45 | 462.49 | - | 280.22 | 743.36 | - |
| ***2-Cl-DMT*** | 6694.22 | - | 601.73 | - | 5110.93 | 3036.69 | 5267.44 |
| ***DMT*** | - | - | 1267.07 | - | - | - | - |
| ***2-Br-Psilacetin*** | - | - | - | - | - | - | - |
| ***Psilacetin*** | - | - | - | 3164.46 | - | - | - |
| ***Reference*** | 1.44 | 14.64 | 0.7 | 4.43 | 0.21 | 0.4 | 21.13 |

***Table S3:*** *Binding affinities (Ki, nM) of test compounds at dopamine receptors, muscarinic receptors, and hERG channels; [a]Radioligand and reference control compound for each receptor: $D_1$ = [$^3$H]-SCH23390 vs (+)-butaclamol; $D_2$ = [$^3$H]-N-methylspiperone vs haloperidol; $D_3$ and $D_4$ = [$^3$H]-N-methylspiperone vs chlorpromazine; $M_4$ and $M_5$ = [$^3$H]-NMS or QNB vs. atropine; hERG = [$^3$H]-Dofetilide vs Dofetilide*

| *Compound* | *m5-HT$_{1A}$* | *m5-HT$_{2A}$* |
|---|---|---|
| | ***$K_i$ nM, (95% CI)*** | |
| ***2-Br-DMT*** | 430.3 | 2,427 |
| | (317.9 – 584.8) | (1,892 – 3,139) |
| ***2-Cl-DMT*** | 561.8 | 2,471 |
| | (402.6 – 787.0) | (1,881 – 3,283) |
| ***DMT*** | 87.95 | 374.6 |
| | (64.46 – 120.8) | (306.1 – 459.0) |
| ***2-Br-psilocin*** | 38.37 | 824.3 |
| | (28.11 – 53.38) | (569.3 – 1,201) |
| ***Psilocin[a]*** | 123.6 | 235.3 |
| | (94.19 – 162.9) | (184.2 – 301.4) |
| ***2-Br-psilacetin*** | 384.7 | 1411 |
| | (257.8 – 579.7) | (1,144 – 1,747) |
| ***Psilacetin*** | 197.7 | 349.1 |
| | (150.7 – 260.7) | (256.5 – 477.6) |

***Table S4:*** *Mouse brain binding affinity data; [a]Psilocin data in this table was reported from previously published results; Glatfelter, G. C., et al. (2022). ACS Pharm. and Trans. Sci, 5(11), 1181–1196.*

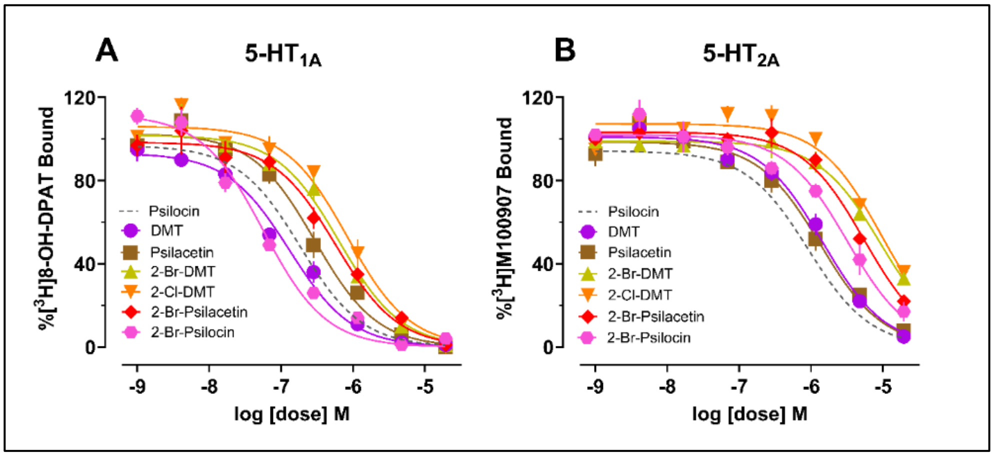


***Figure S1:*** *Mouse brain binding affinity data of 2-halogenated tryptamines*

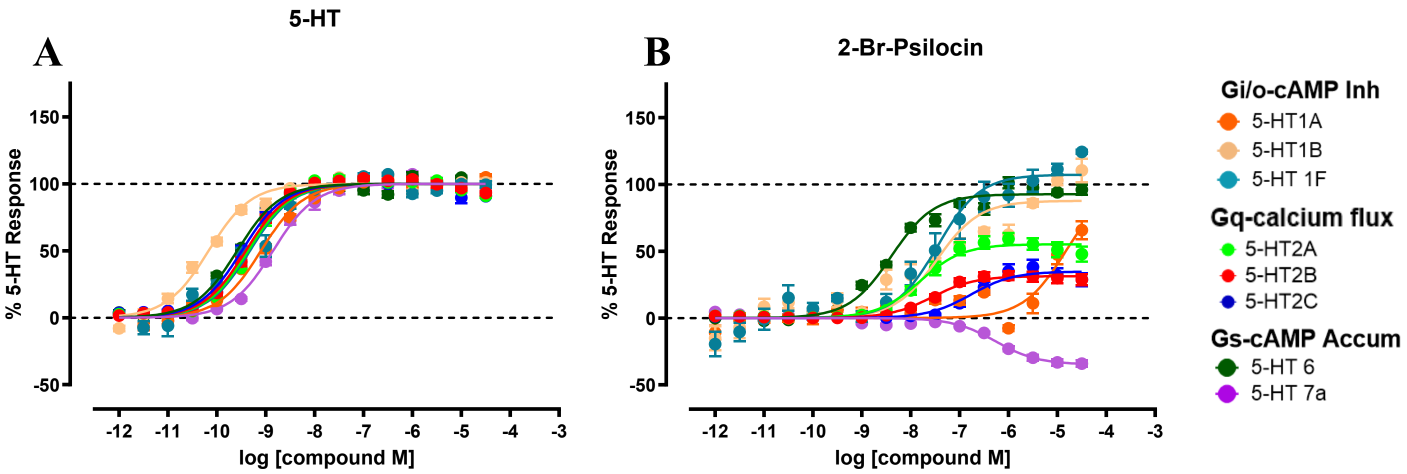


***Figure S2:*** *Second messenger activity comparing 5-HT (A) to 2-Br-Psilocin (B) at 5-HT$_{1A/1B/1D}$ receptors measuring Gi/o-cAMP inhibition, 5-HT$_{2A/2B/2C}$ receptors measuring Gq-mediated calcium flux, and 5-HT$_6$ and 5-HT$_{7a}$ receptors measuring Gs-mediated cAMP accumulation. Data are normalized to percent 5-HT response and represent the mean and SEM from three or more independent experiments.*

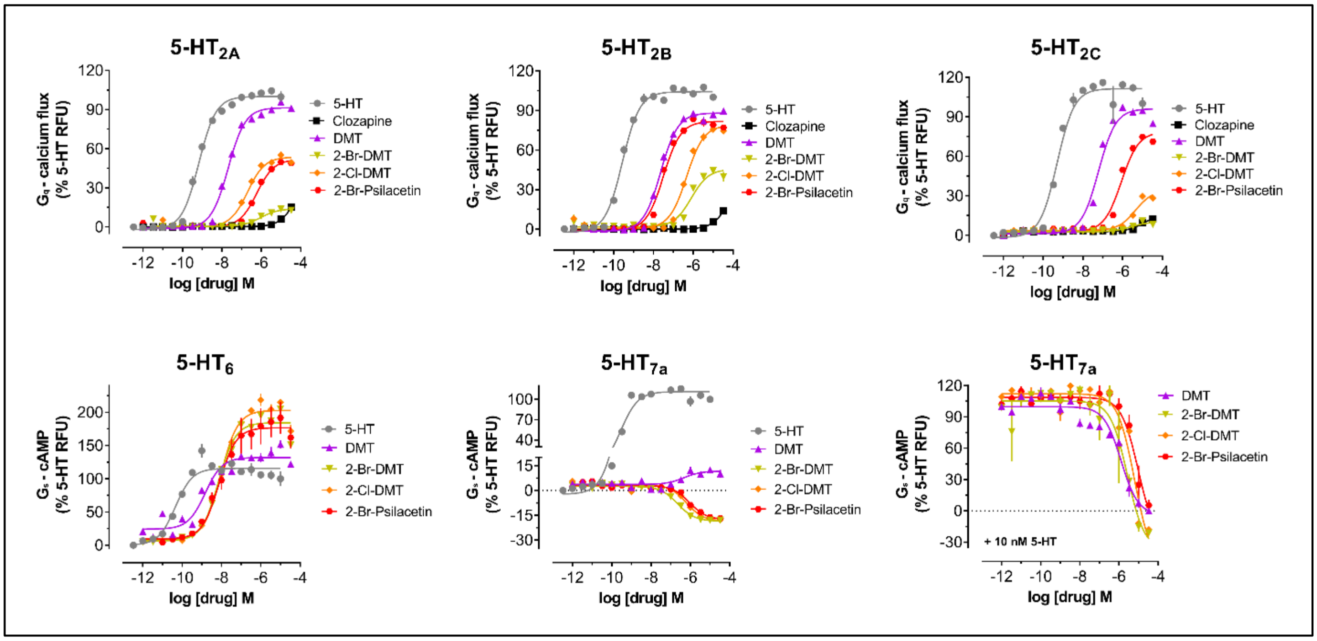


***Figure S3:*** *Gq mediated calcium flux and cAMP second messenger assays for DMT, psilacetin, 2-Cl-DMT, and 2-Br-DMT at 5-HT$_{2A-C}$, 5-HT$_6$, and 5-HT$_7$ receptors.*

| ***Compound*** | ***5-HT$_{2A}$*** | ***5-HT$_{2B}$*** | ***5-HT$_{2C}$*** | ***5-HT$_6$*** | ***5-HT$_{7a}$*** |
|---|---|---|---|---|---|
| | Gq - Calcium mobilization assay | | | Gs - split luciferase cAMP | |
| ***2-Br-DMT*** | 8,107 | 299 | 2,492 | **7** | 225 |
| | (4,428 - 14,840) | (554 - 1,151) | (945 - 6,572) | (5 - 9) | (178 - 286) |
| | $E_{max}$ = 0 | $E_{max}$ = 45 | $E_{max}$ = 10 | $E_{max}$ **= 184** | $E_{max}$ = **-19** |
| ***2-Cl-DMT*** | 202 | 461 | 4,479 | **9** | 630 |
| | (155 - 263) | (380 - 559) | (3,133 - 6,403) | (7 - 11) | (464 - 854) |
| | $E_{max}$ = 54 | $E_{max}$ = 79 | $E_{max}$ = 34 | $E_{max}$ **= 203** | $E_{max}$ = -19 |
| ***DMT*** | 21 | 23 | 61 | **1** | 467 |
| | (18-24) | (20-27) | (49 - 77) | (0.9 - 2) | (124 - 1,750) |
| | $E_{max}$ = 91 | $E_{max}$ = 88 | $E_{max}$ = 96 | $E_{max}$ = 132 | $E_{max}$ = 12 |
| ***2-Br-Psilacetin*** | **598** | **32** | 864 | **8** | 917 |
| | (504 - 709) | (27-38) | (696 - 1,072) | (4 - 13) | (606 - 1,389) |
| | $E_{max}$ **= 52** | $E_{max}$ = 82 | $E_{max}$ = 78 | $E_{max}$ = 177 | $E_{max}$ = **-18** |
| ***Psilacetin**** | 328 | 88 | 408 | N.T. | N.T. |
| | (227-474) | (69 - 114) | (318 - 525) | | |
| | $E_{max}$ = 54 | $E_{max}$ = 29 | $E_{max}$ = 32 | N.T. | N.T |

***Table S5:*** *Second Messenger assay, $EC_{50}$ (nM), (95%, CI), $E_{max}$*

| ***Table S5: hERG inhibition*** | |
|---|---|
| ***Compound*** | **$IC_{50}$ (nM)** |
| ***2-Br-psilocin*** | >30 x $10^3$ |
| ***2-Br-psilacetin*** | 4,900 |
| ***Verapamil*** | 230, 350 |

***Table S6:*** *hERG inhibition for 2-Br-psilocin*

| ***BBB Permeability (PAMPA)*** | |
|---|---|
| ***Compound*** | **Avg. Pe (x 10-6 cm/s)** |
| ***2-Br-DMT*** | 70.22 |
| ***2-Cl-DMT*** | 68.67 |
| ***DMT*** | 59.08 |
| ***2-Br-Psilacetin*** | 85.5 |
| ***2-Br-Psilocin*** | 94.1 |
| ***Psilacetin*** | 50.75 |
| ***Verapamil (+)*** | 167.91 |
| ***Lidocaine*** | 82 |
| ***Theophylline (-)*** | 4.33 |

***Table S7:*** *Passive permeability through PAMPA for prediction of BBB permeability*

| 10 mg/kg, PO | $t_{1/2}$ | $T_{max}$ | $C_{max}$ | AUClast | AUCInf | AUC/D | AUC Extra | MRT | Vz_F | CL_F | F |
|---|---|---|---|---|---|---|---|---|---|---|---|
| | (h) | (h) | (ng/mL) | (h*ng/mL) | (h*ng/mL) | (h*kg*ng/mL/mg) | (%) | (h) | (L/kg) | (mL/min/kg) | % |
| **Rat#1** | 2.33 | 1.00 | 68 | 220.49 | 244 | 24.40 | 9.63 | 3.40 | 137.50 | 683.13 | 86.97 |
| **Rat#2** | 1.30 | 1.00 | 44 | 122.24 | 124 | 12.41 | 1.51 | 1.92 | 150.74 | 1342.85 | 48.22 |
| **Rat#3** | 1.36 | 0.50 | 58 | 161.41 | 165 | 16.53 | 2.37 | 2.01 | 118.52 | 1008.11 | 63.67 |
| **Mean** | 1.66 | 0.83 | 57 | 168.05 | 178 | 17.78 | 4.50 | 2.44 | 135.59 | 1011.36 | 66.29 |
| **SD** | 0.58 | 0.29 | 12 | 49.46 | 61 | 6.09 | 4.46 | 0.83 | 16.20 | 329.87 | 19.51 |

***Table S8.*** *The PK parameters of 2-Br-psilocin in Rat plasma samples after PO administration of 10 mg/kg 2-Br-psilacetin (10 mg/kg, PO) group, oral bioavailability highlighted*

| Sample | Route | Dose (mg/kg) | Time (h) | Avg. Brain Concentration (ng/g) | Avg. Plasma Concentration ((ng/mL) | Brain/Plasma Ratio |
|---|---|---|---|---|---|---|
| **2-Br-Psilacetin** | IV | 1 | 0.5 | 1109 | 39 | 30.00 |
| | | | 1 | 375 | 21 | 19.08 |
| | | | 4 | 14 | 2 | 10.75 |
| | PO | 10 | 0.5 | 2300 | 96 | 24.67 |
| | | | 1 | 2502 | 87 | 28.76 |
| | | | 4 | 590 | 21 | 27.59 |

***Table S9:*** *The Exposure level of 2-Br-Psilocin in Rat samples upon administration of 2-Br-Psilacetin (1 mg/kg, IV and 10 mg/kg, PO)*

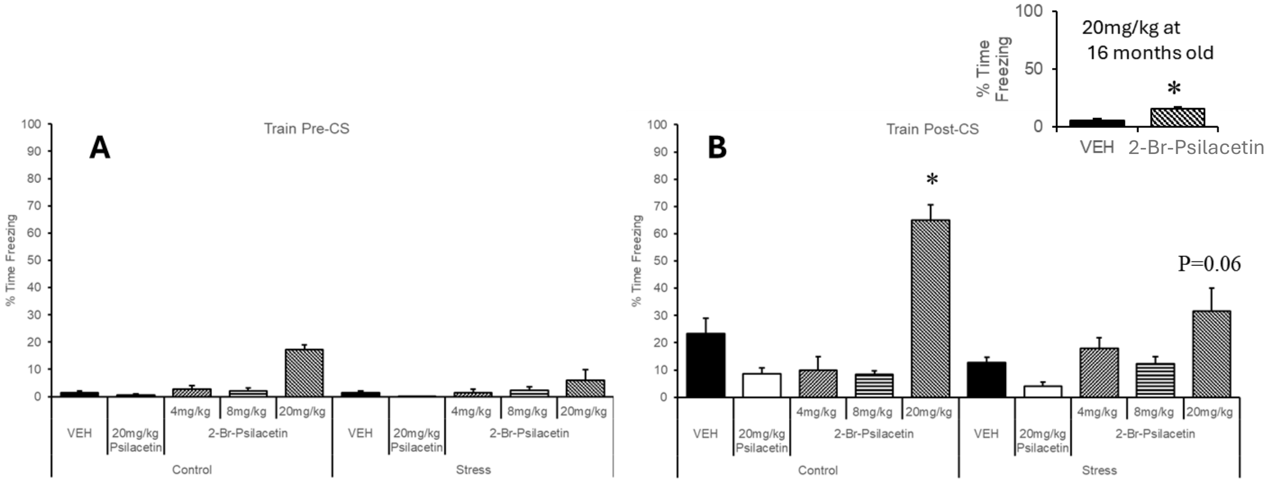


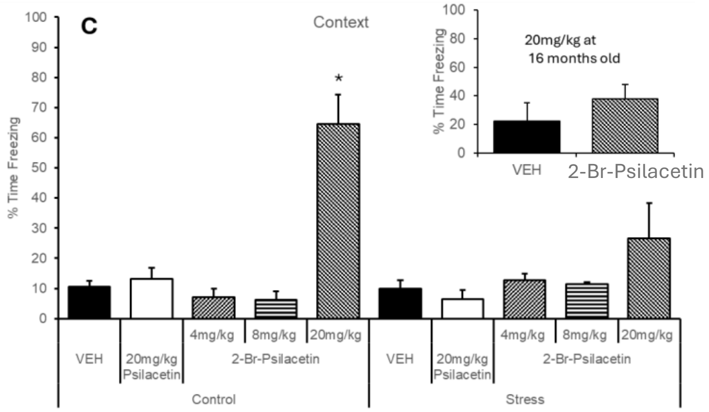


***Figure S4**: (A-B) Fear conditioning, trained conditioned stimulus in C57BL/6 mice, stressed mice, and 16-month-old mice (inset chart); (C) Fear conditioning, context, in C57BL/6 mice, stressed mice, and 16-month-old mice (inset chart)*

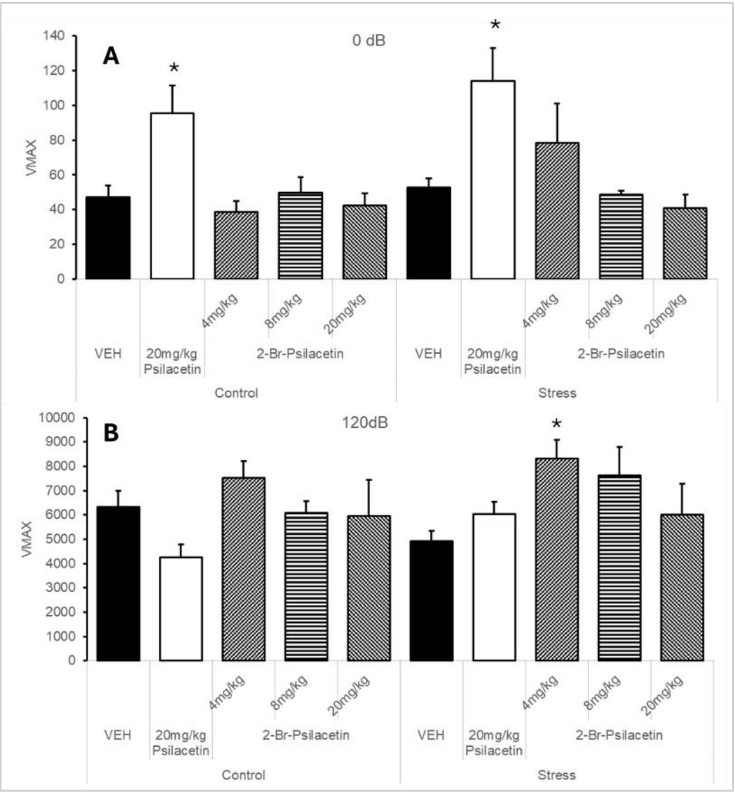


***Figure S5 (left):** Startle response at (A) baseline and (B) 120 dB startle in C57BL/6 mice*

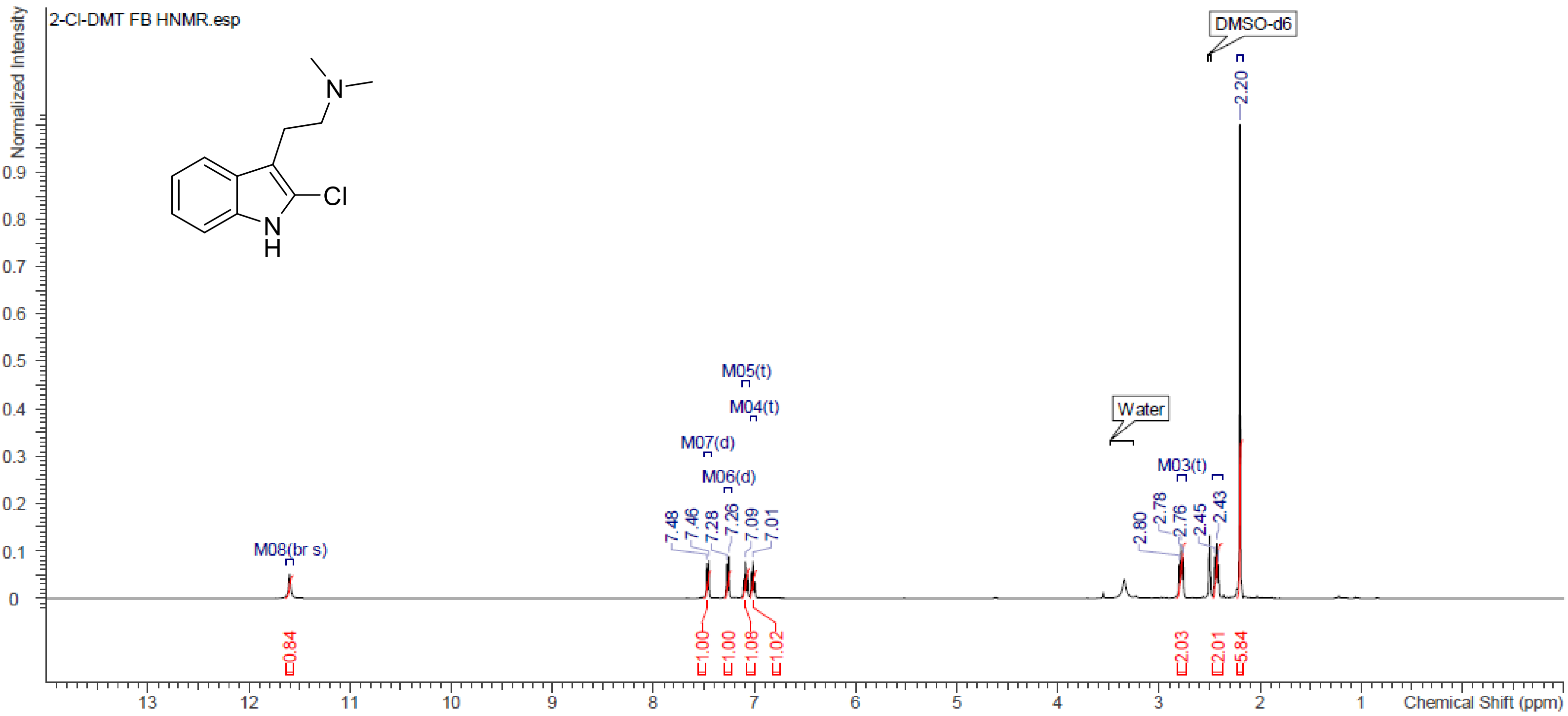


**Figure S6.** $^{1}$H NMR (DMSO-d$_6$, 400 MHz) spectrum 2-Cl-DMT

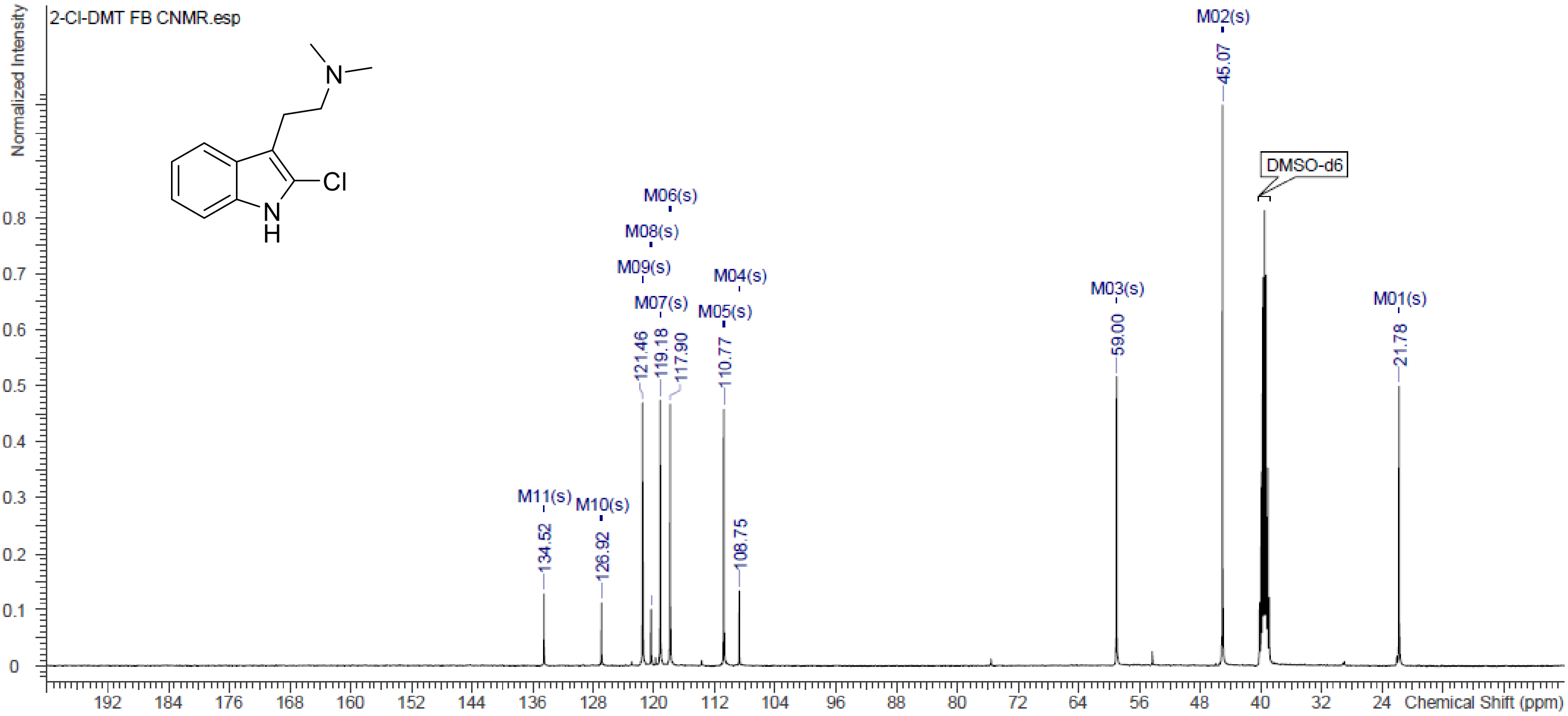


**Figure S7.** $^{13}$C NMR (DMSO-d$_6$, 101 MHz) spectrum 2-Cl-DMT

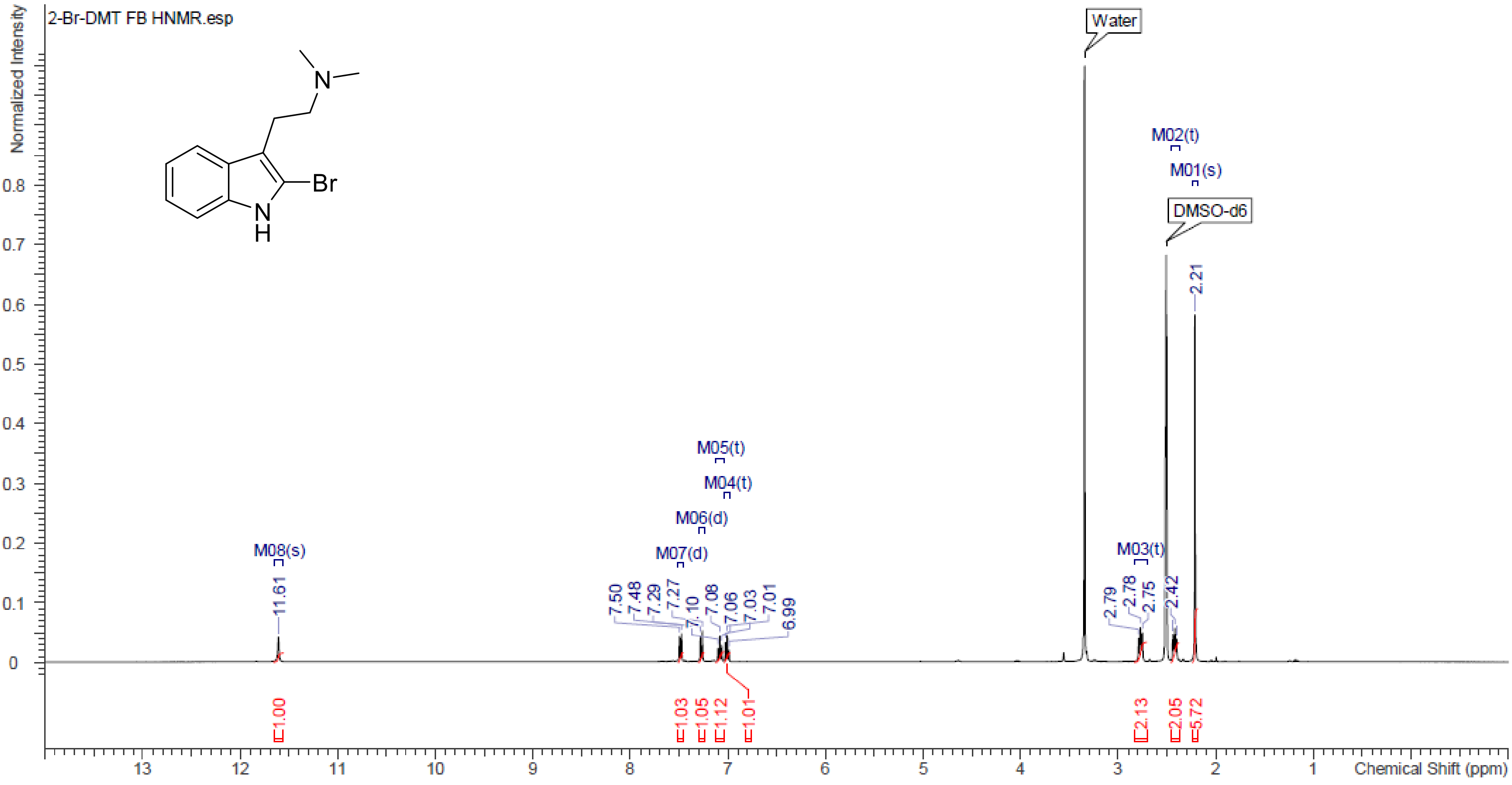


**Figure S8.** $^{1}$H NMR (DMSO-$d_6$, 400 MHz) spectrum 2-Br-DMT

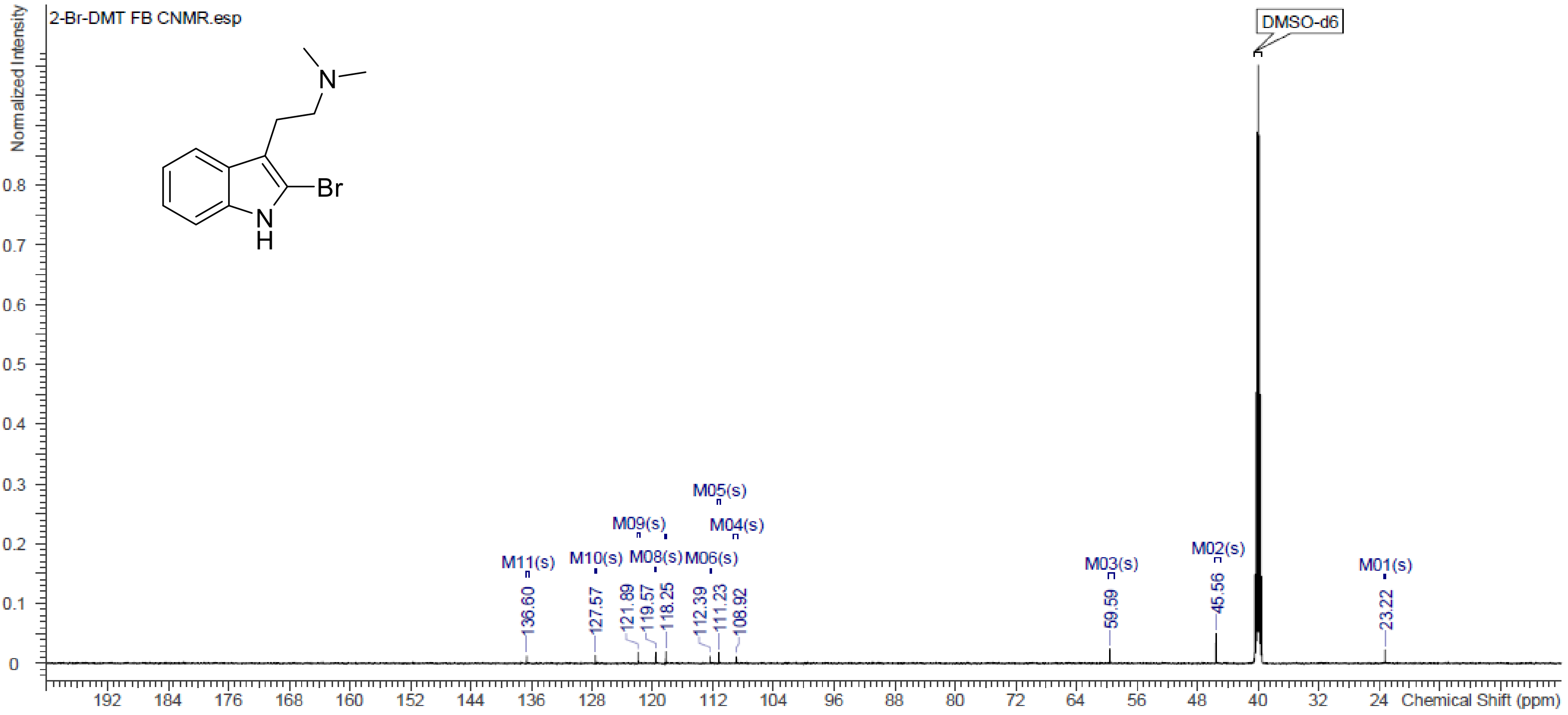


**Figure S9.** $^{13}$C NMR (DMSO-$d_6$, 151 MHz) spectrum 2-Br-DMT

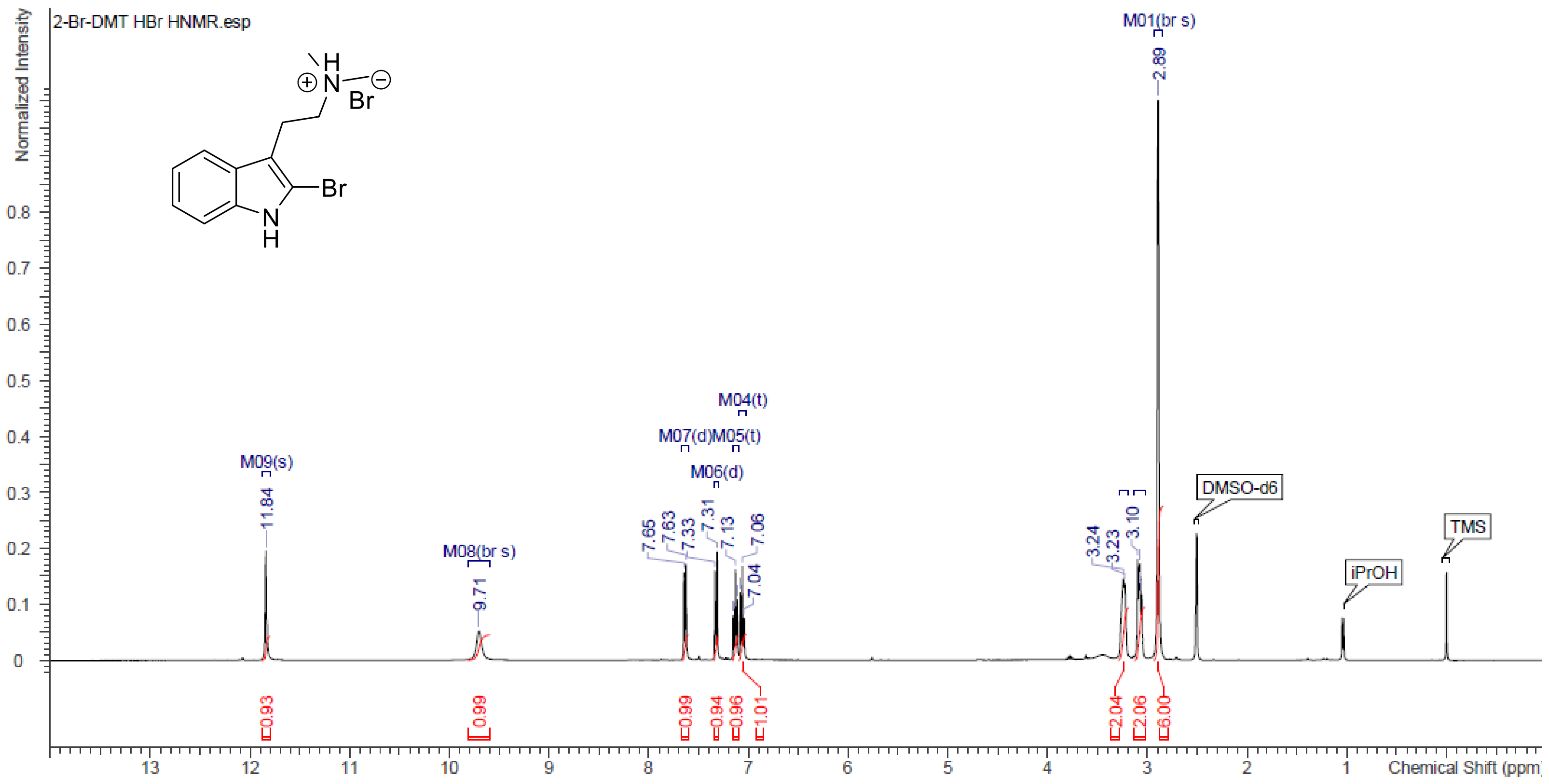


**Figure S10.** $^{1}$H NMR (DMSO-d$_6$, 400 MHz) spectrum 2-Br-DMT HBr

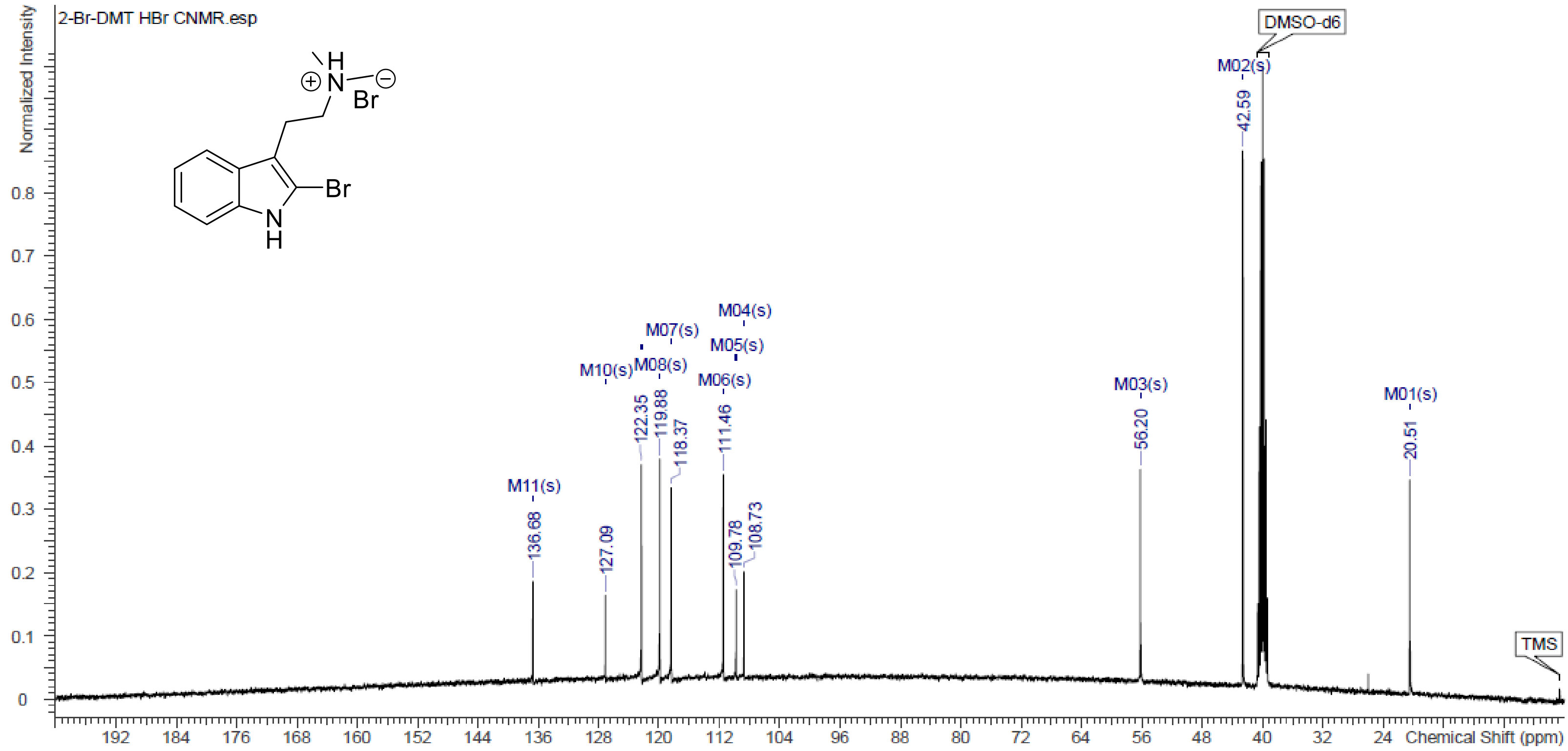


**Figure S11.** $^{13}$C NMR (DMSO-d$_6$, 101 MHz) spectrum 2-Br-DMT HBr

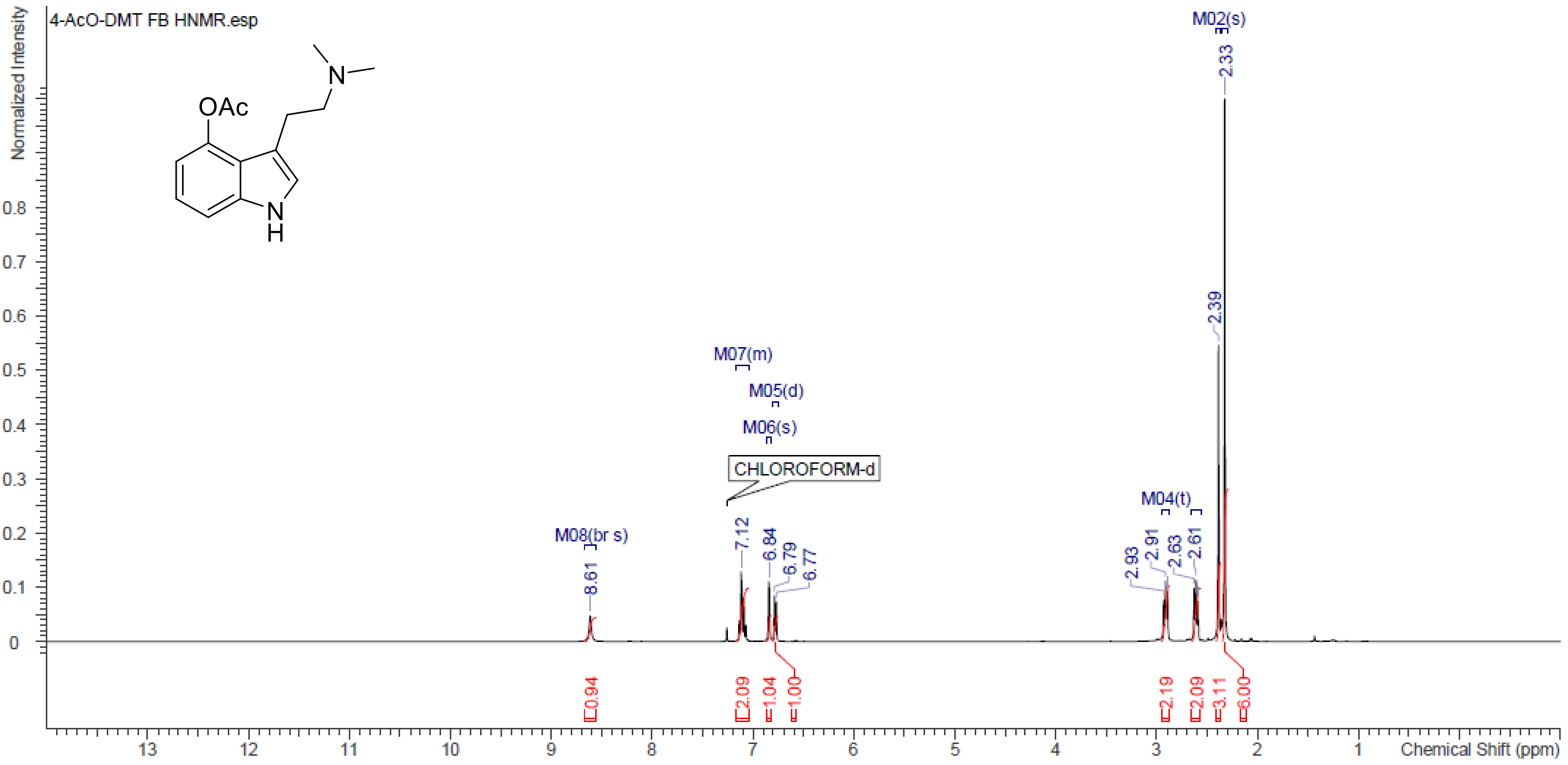


**Figure S12.** $^{1}$H NMR ($CDCl_3$, 400 MHz) spectrum 4-AcO-DMT

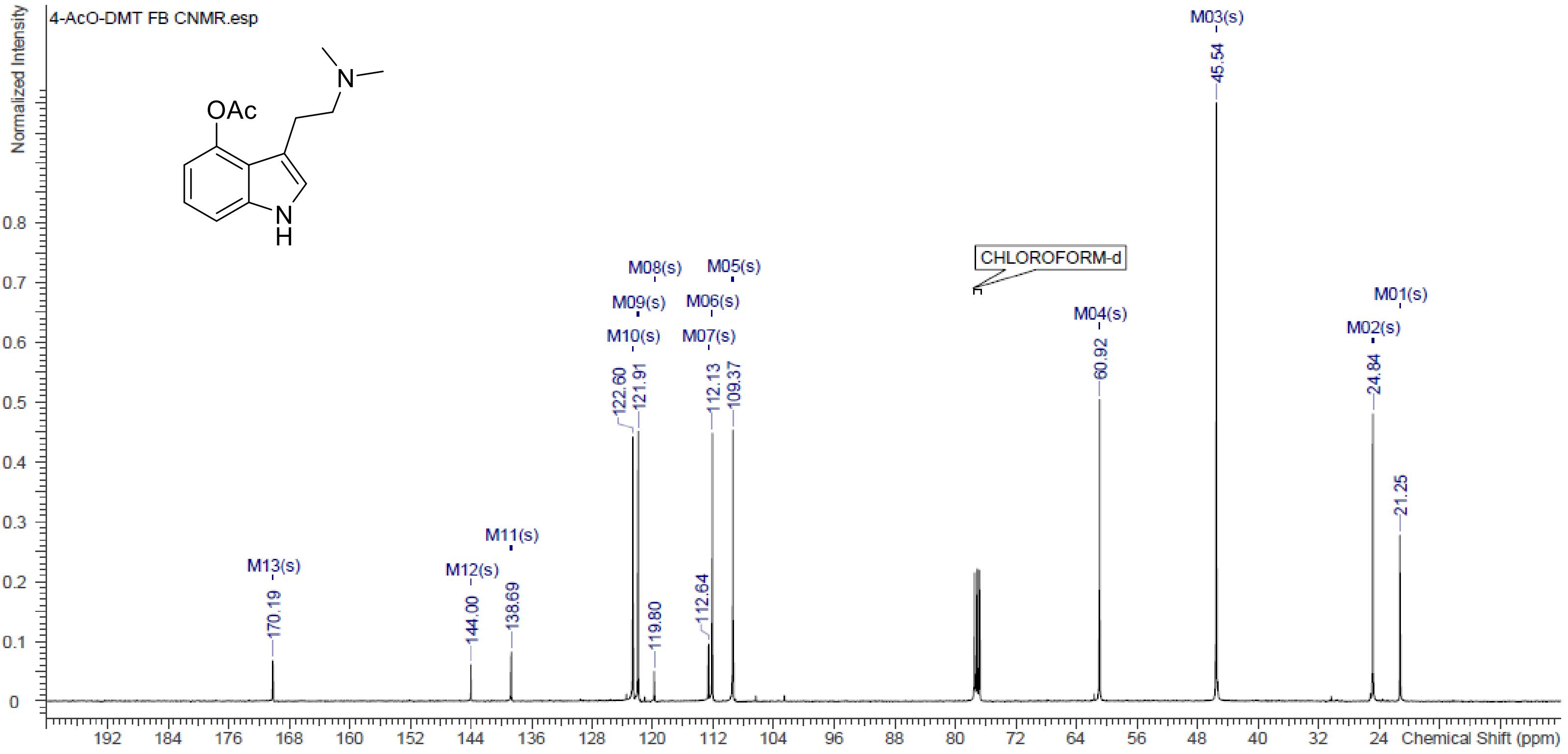


**Figure S13.** $^{13}$C NMR ($CDCl_3$, 101 MHz) spectrum 4-AcO-DMT

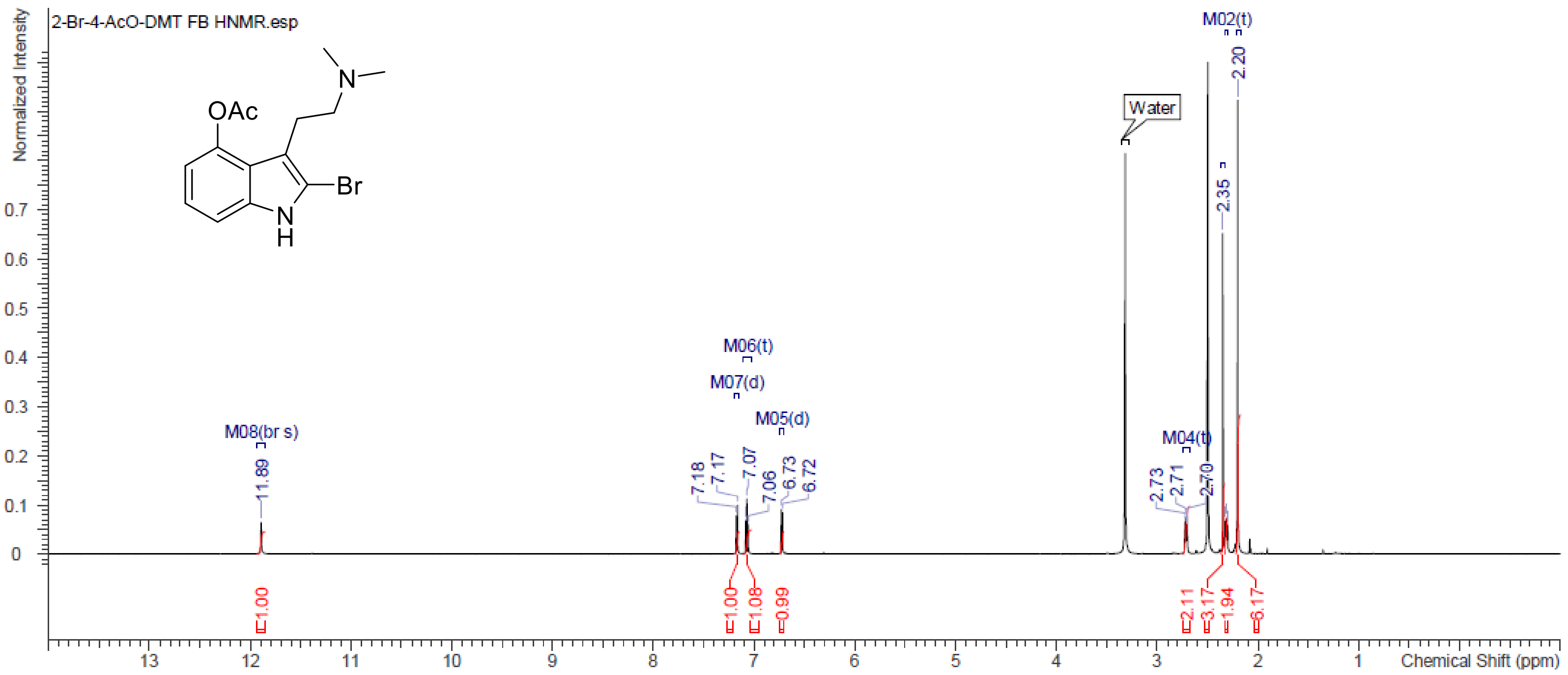

**Figure S14.** $^{1}$H NMR (DMSO-d$_6$, 600 MHz) spectrum 2-Br-4-AcO-DMT

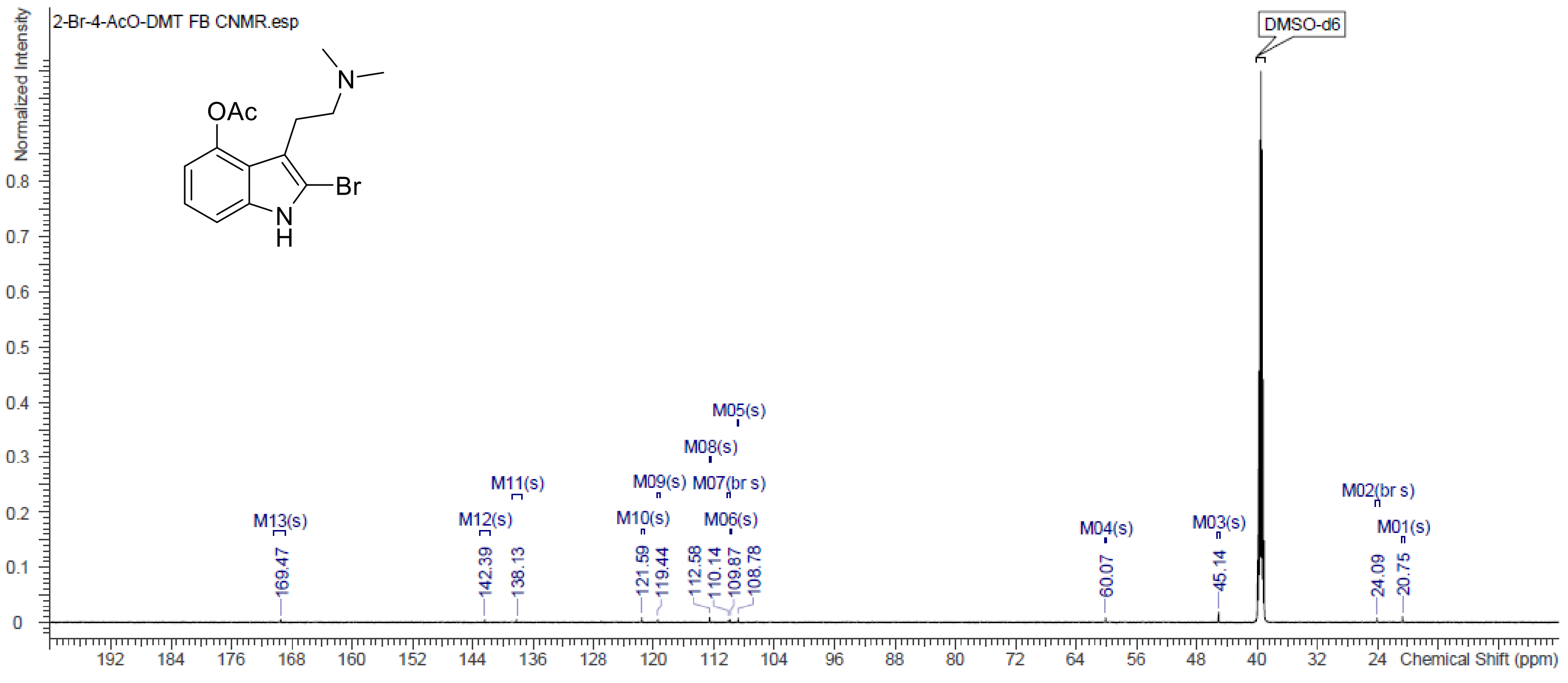

**Figure S15.** $^{13}$C NMR (DMSO-d$_6$, 151 MHz) spectrum 2-Br-4-AcO-DMT.

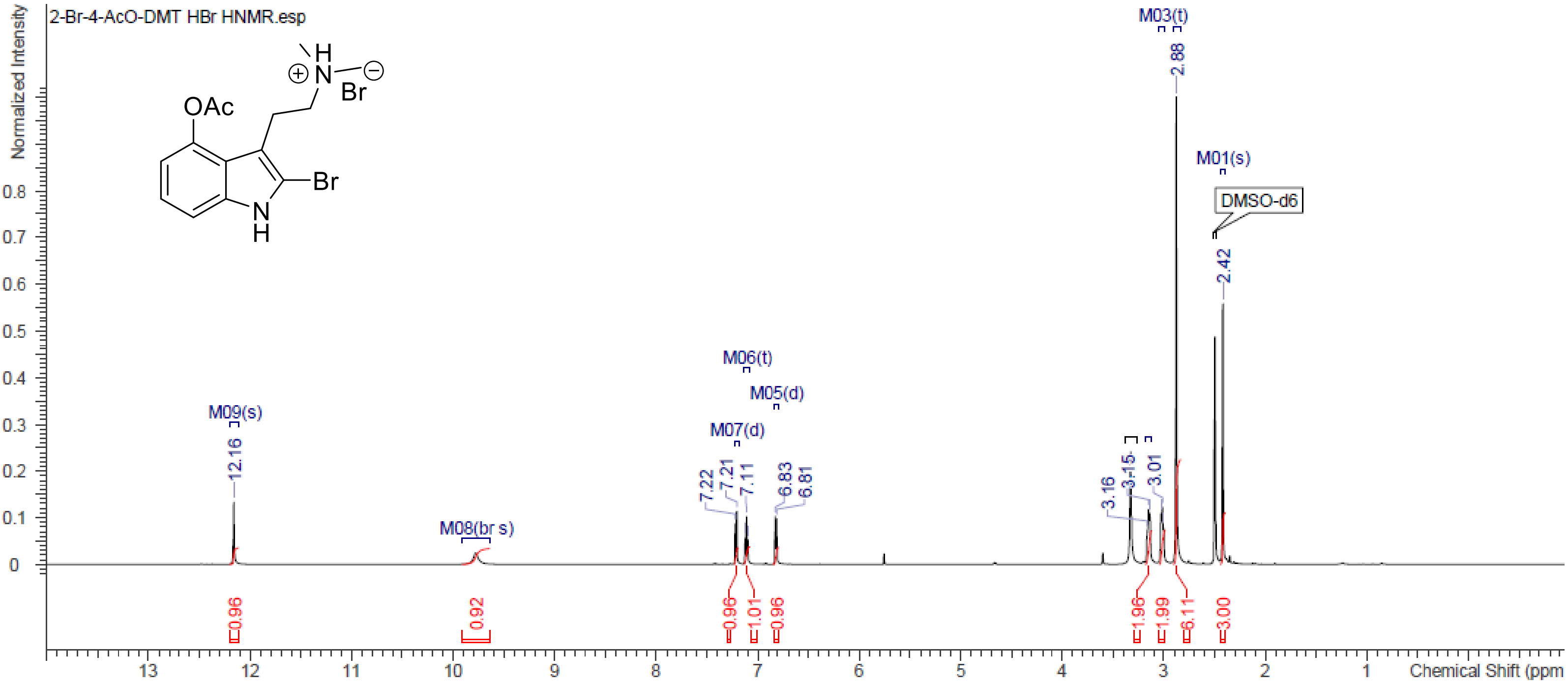


**Figure S16.** $^{1}H$ NMR (DMSO-$d_6$, 600 MHz) spectrum 2-Br-4-AcO-DMT HBr

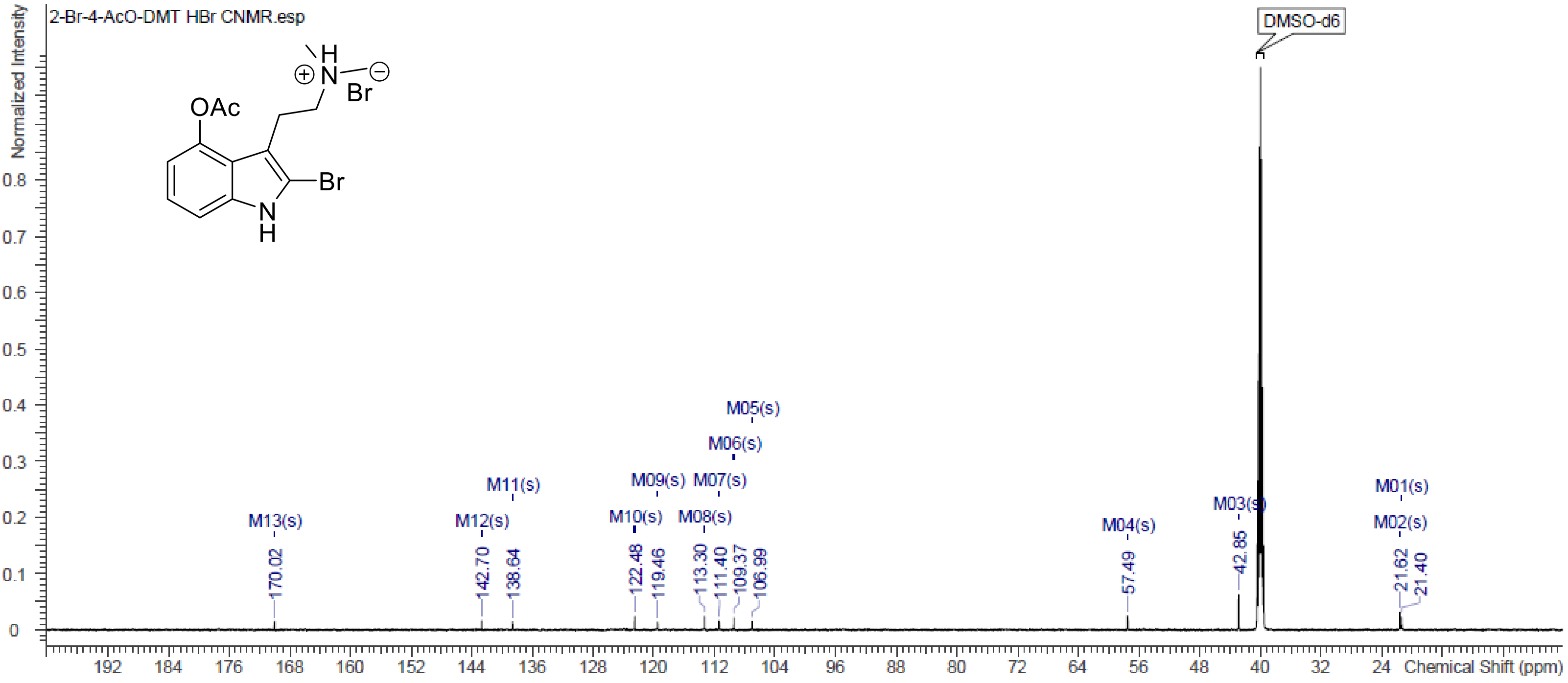


**Figure S17.** $^{13}C$ NMR (DMSO-$d_6$, 151 MHz) spectrum 2-Br-4-AcO-DMT HBr

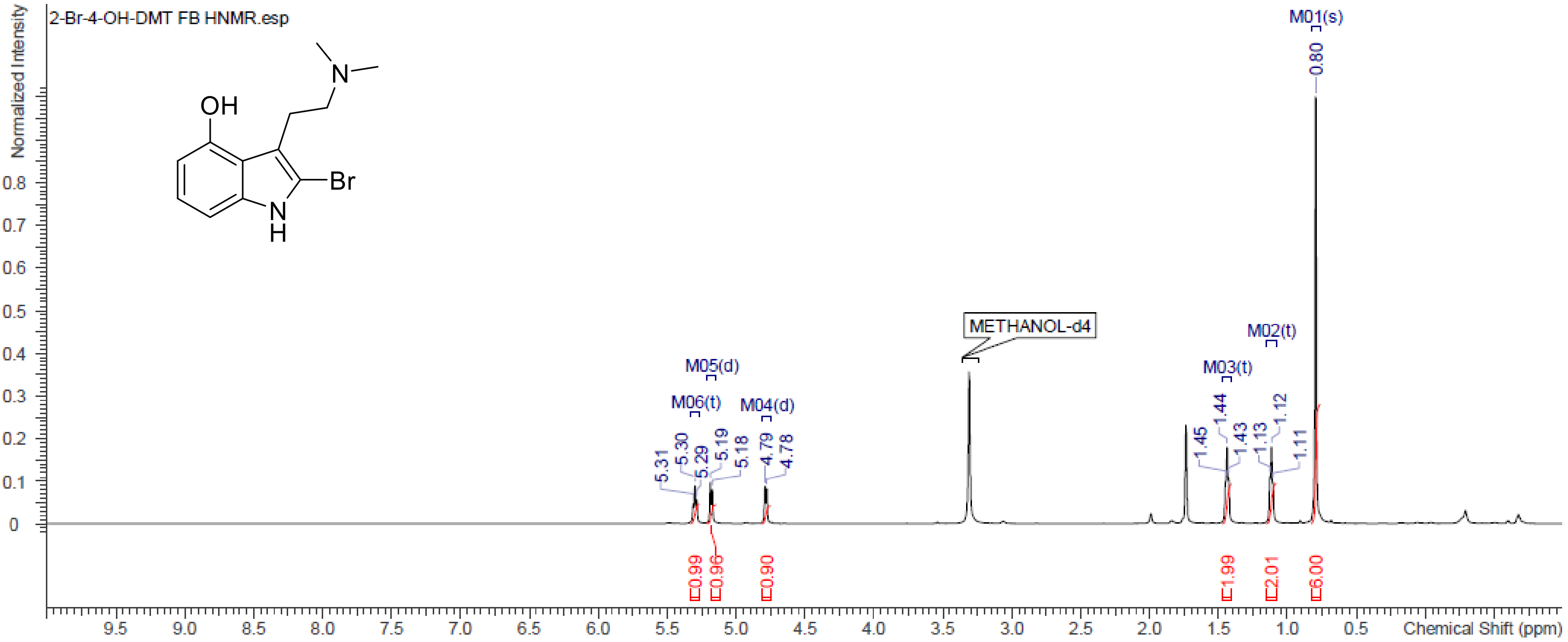


**Figure S18.** $^{1}$H NMR (MeOD, 600 MHz) spectrum 2-Br-4-OH-DMT

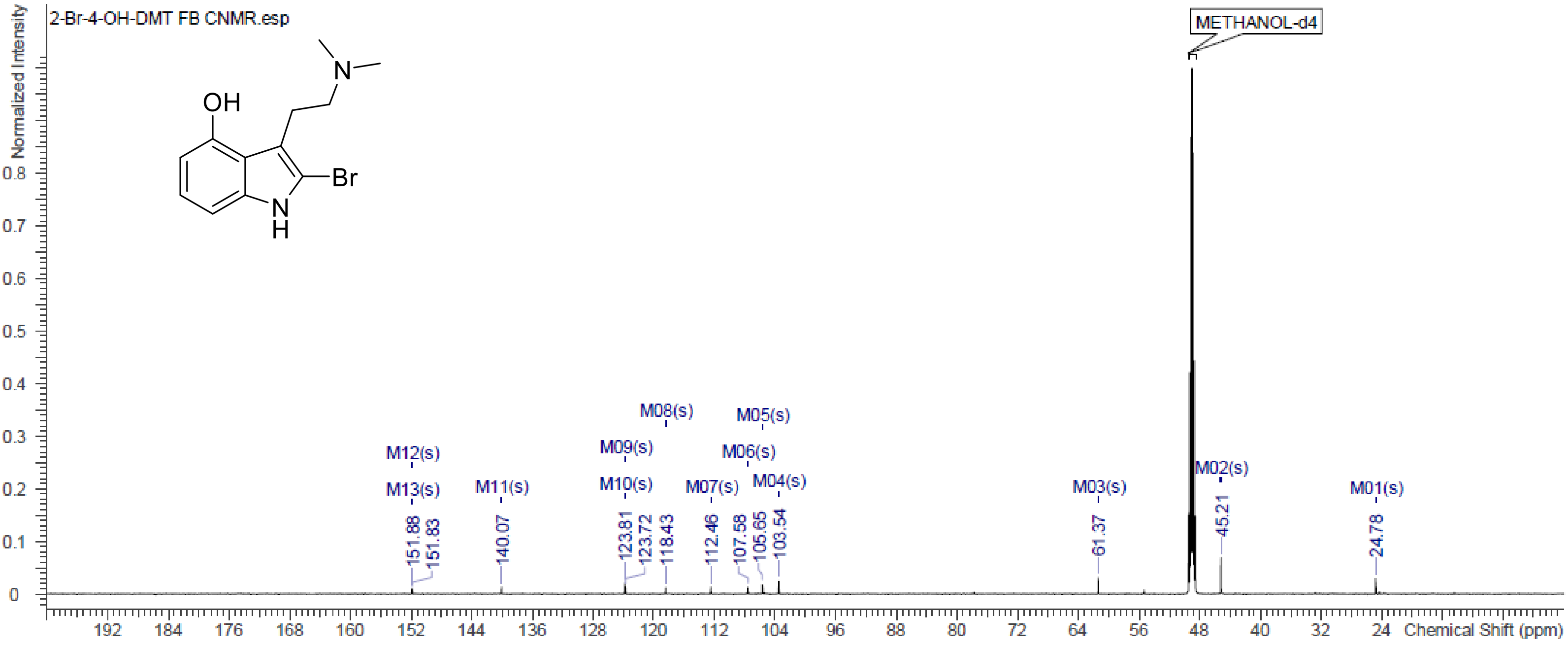


**Figure S19.** $^{13}$C NMR (MeOD, 151 MHz) spectrum 2-Br-4-OH-DMT

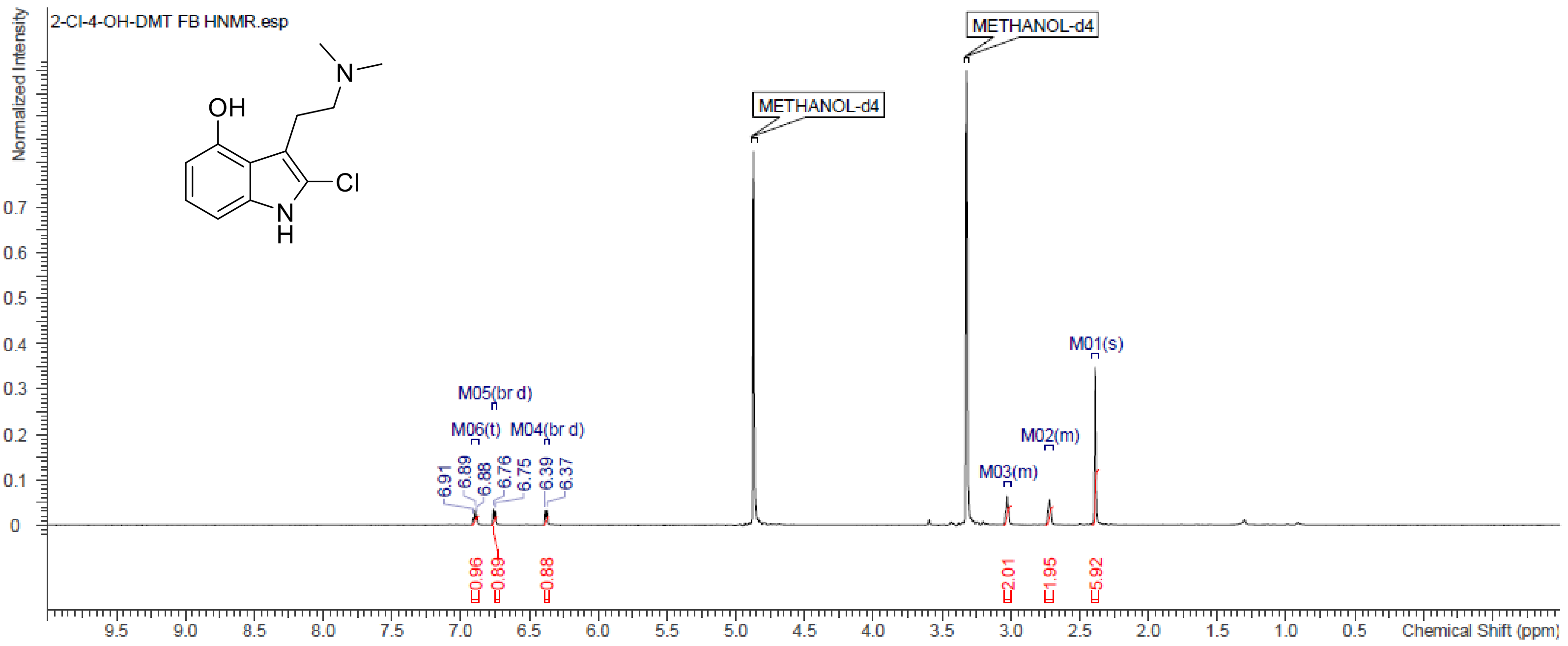


**Figure S20.** $^{1}$H NMR (MeOD, 600 MHz) spectrum 2-Cl-4-OH-DMT

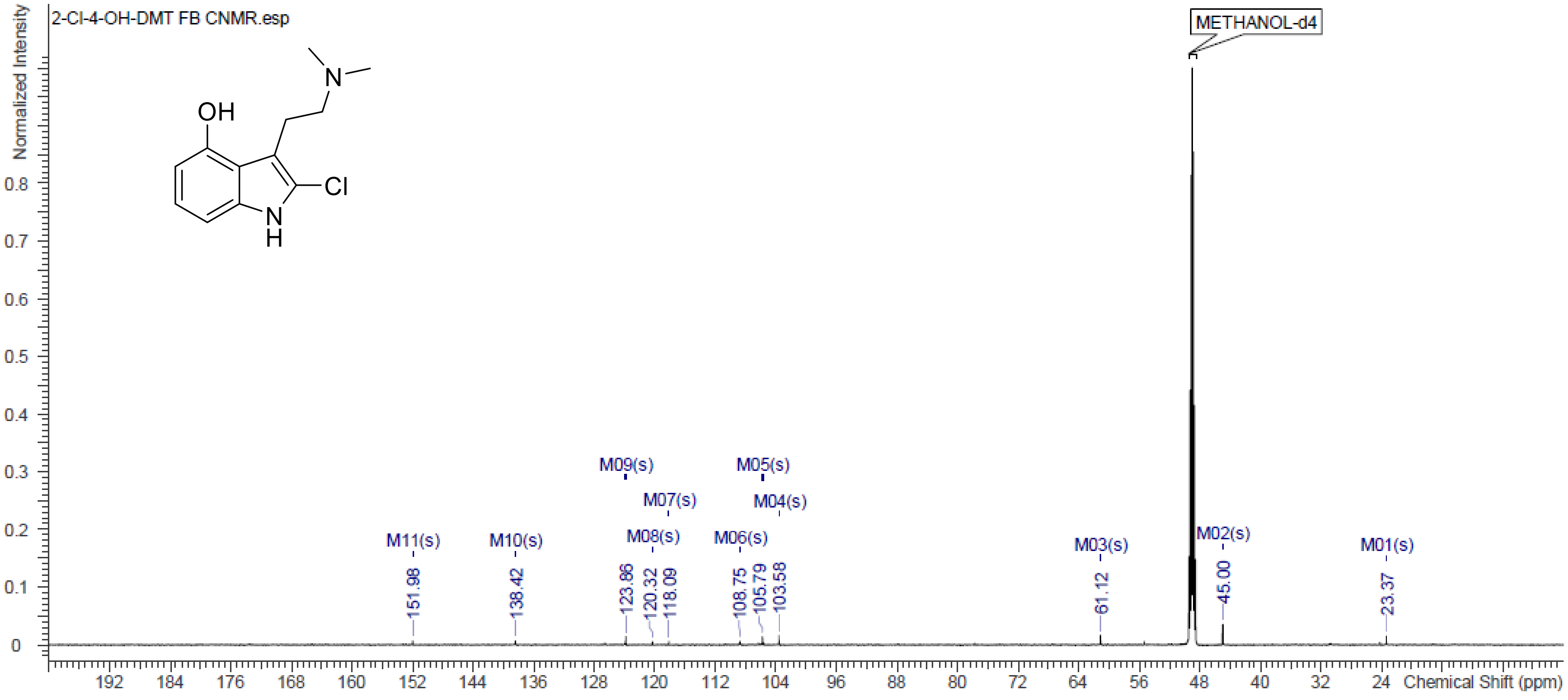


**Figure S21.** $^{13}$C NMR (MeOD, 151 MHz) spectrum 2-Cl-4-OH-DMT

### HRMS Spectra of 2-halogenated tryptamines

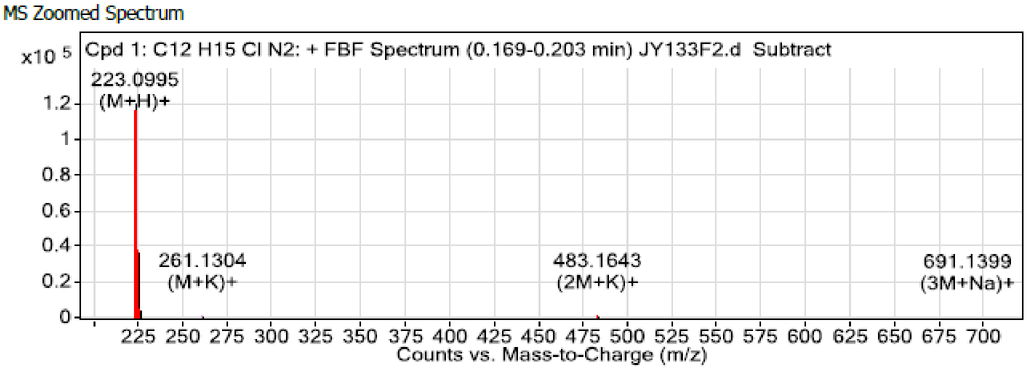


**Figure S22.** 2-Cl-DMT HRMS $[M+H]^+$ spectrum

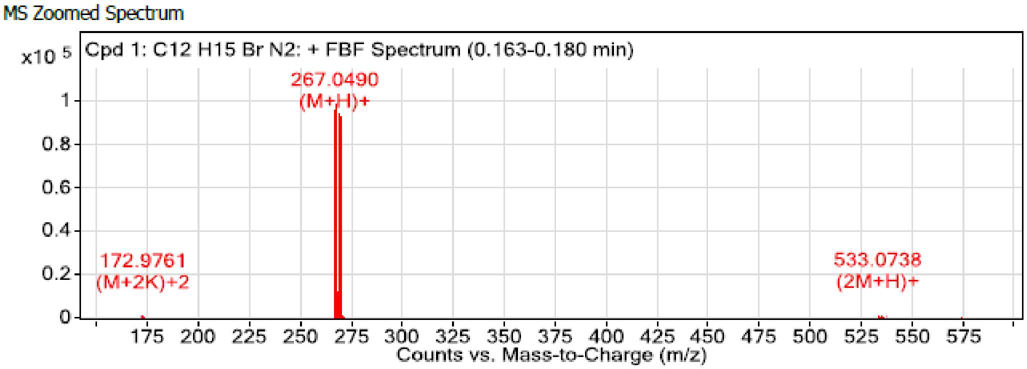


**Figure S23.** 2-Br-DMT HRMS $[M+H]^+$ spectrum

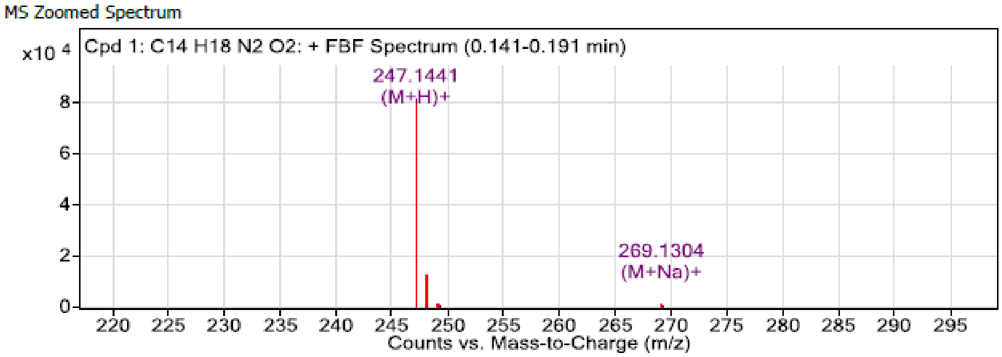


**Figure S24.** 4-AcO-DMT HRMS $[M+H]^+$ spectrum

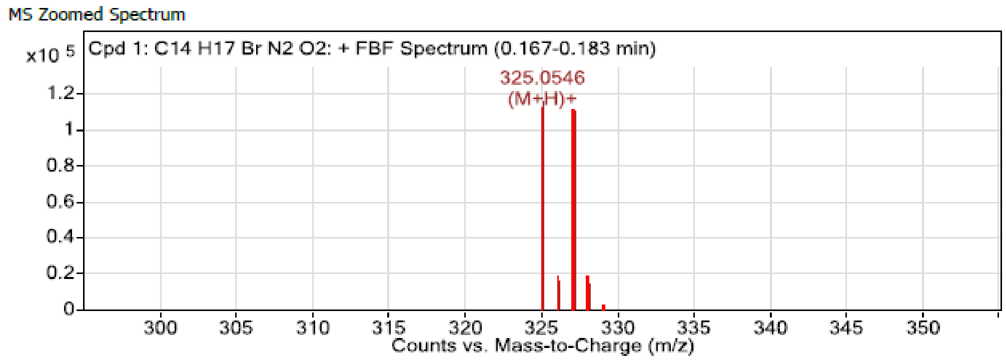


**Figure S25.** 2-Br-4-AcO-DMT HRMS $[M+H]^+$ spectrum

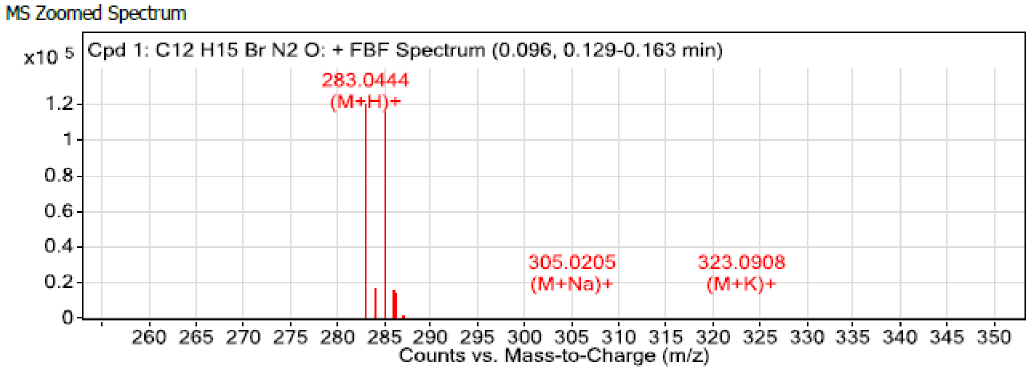


**Figure S26.** 2-Br-4-OH-DMT HRMS $[M+H]^+$ spectrum

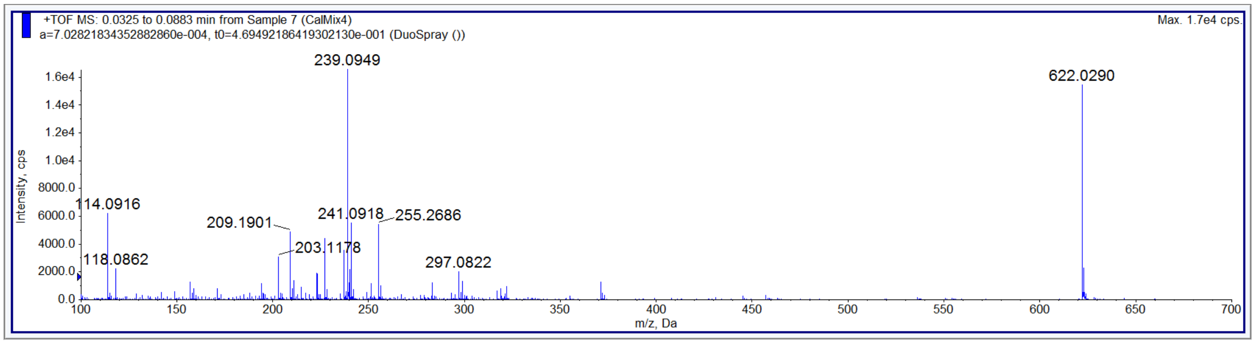


**Figure S27.** 2-Cl-4-OH-DMT HRMS spectrum

**UV Purity of 2-halogenated tryptamines**

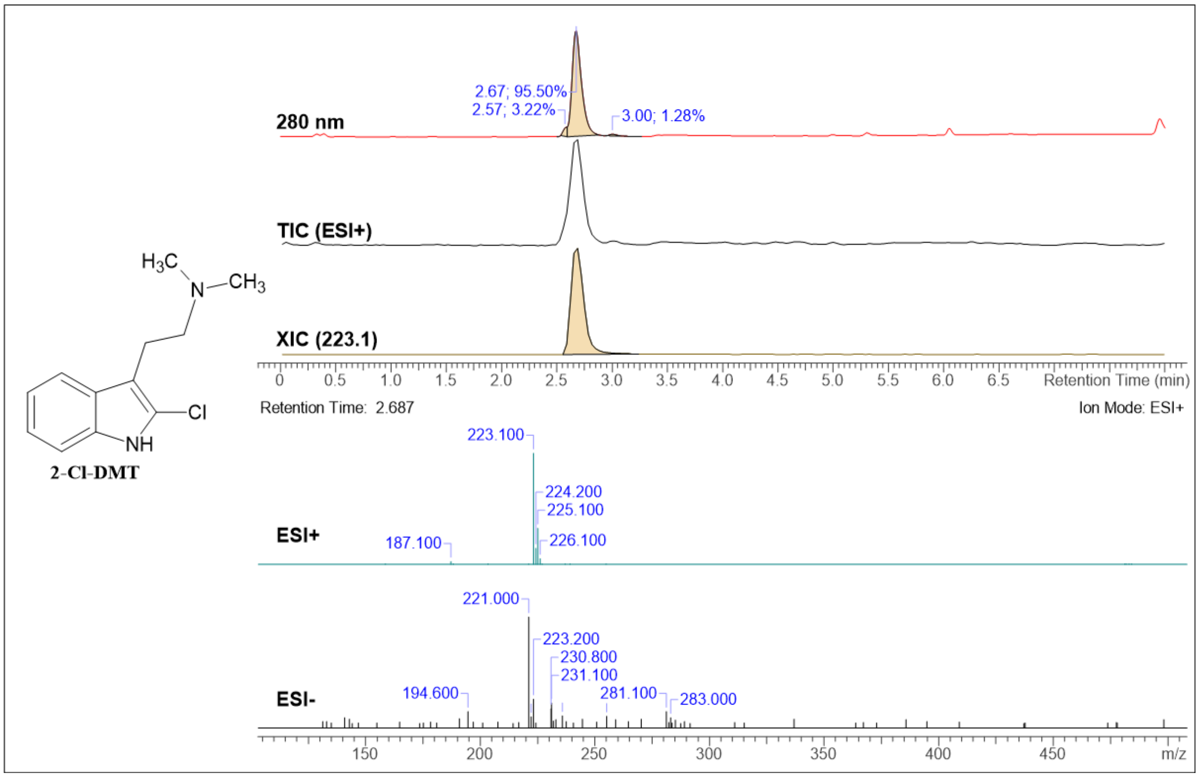


**Figure S28.** UV purity of 2-Cl-DMT

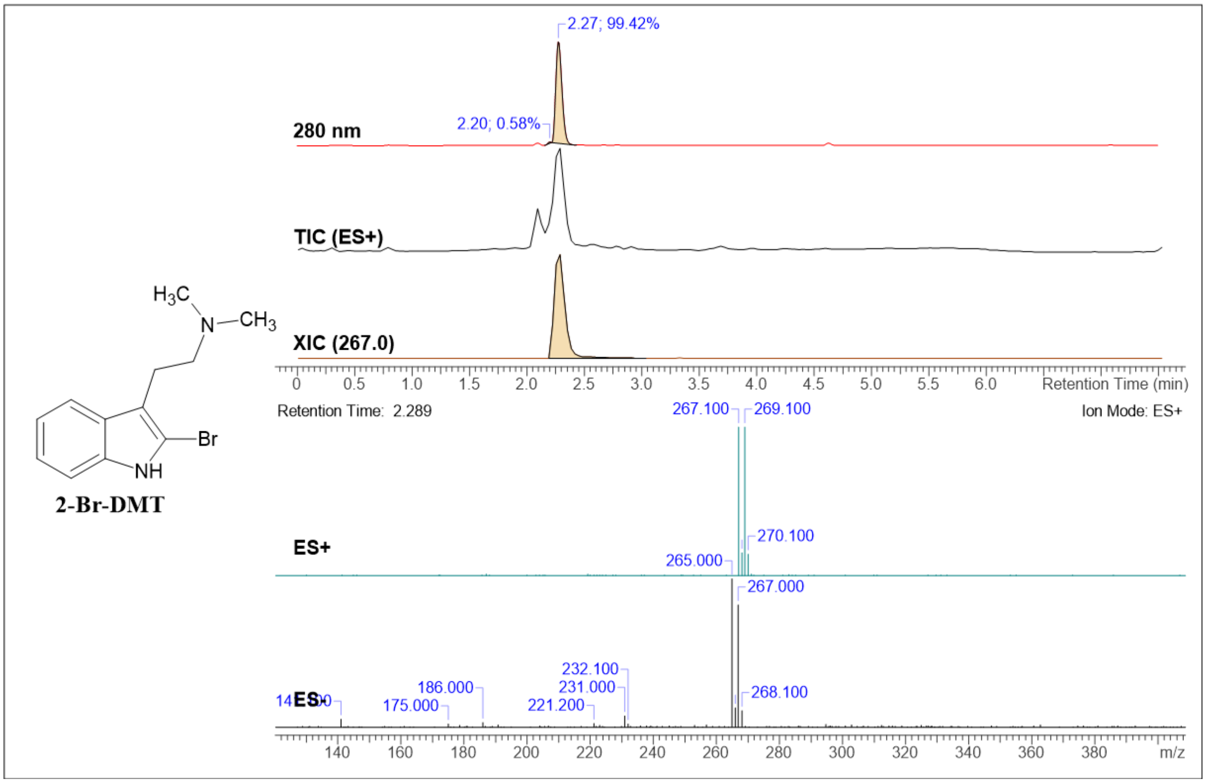


**Figure S29.** UV purity of 2-Br-DMT

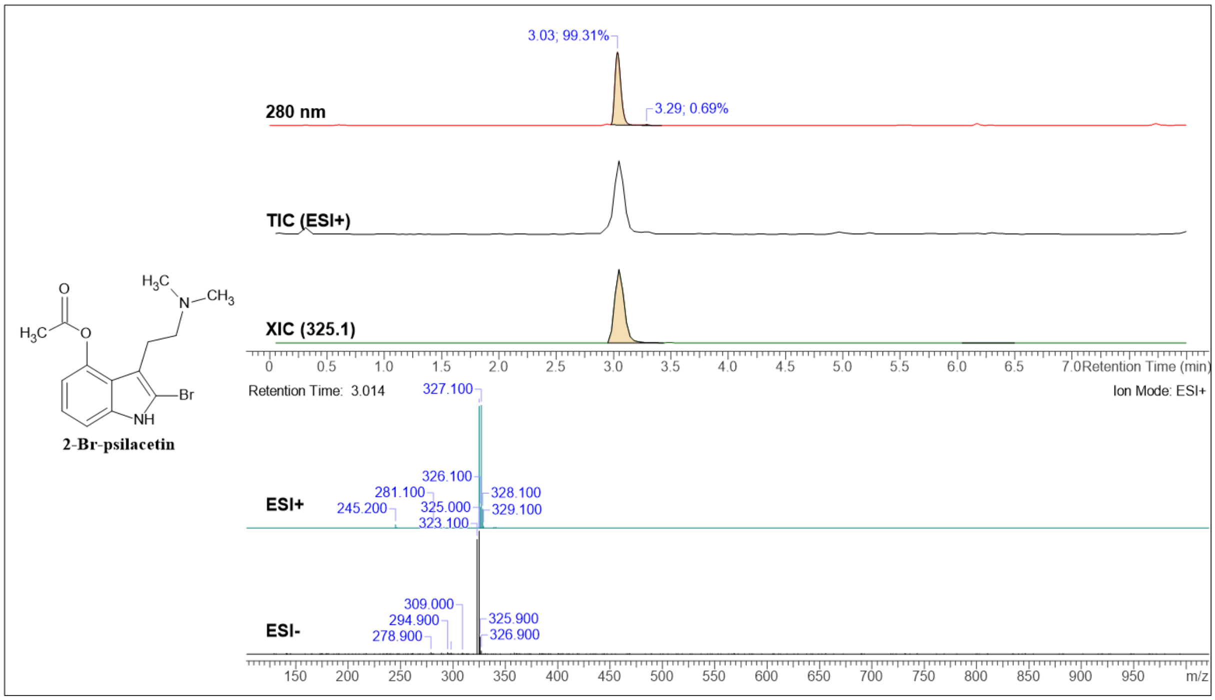


**Figure S30.** UV purity of 2-Br-psilacetin

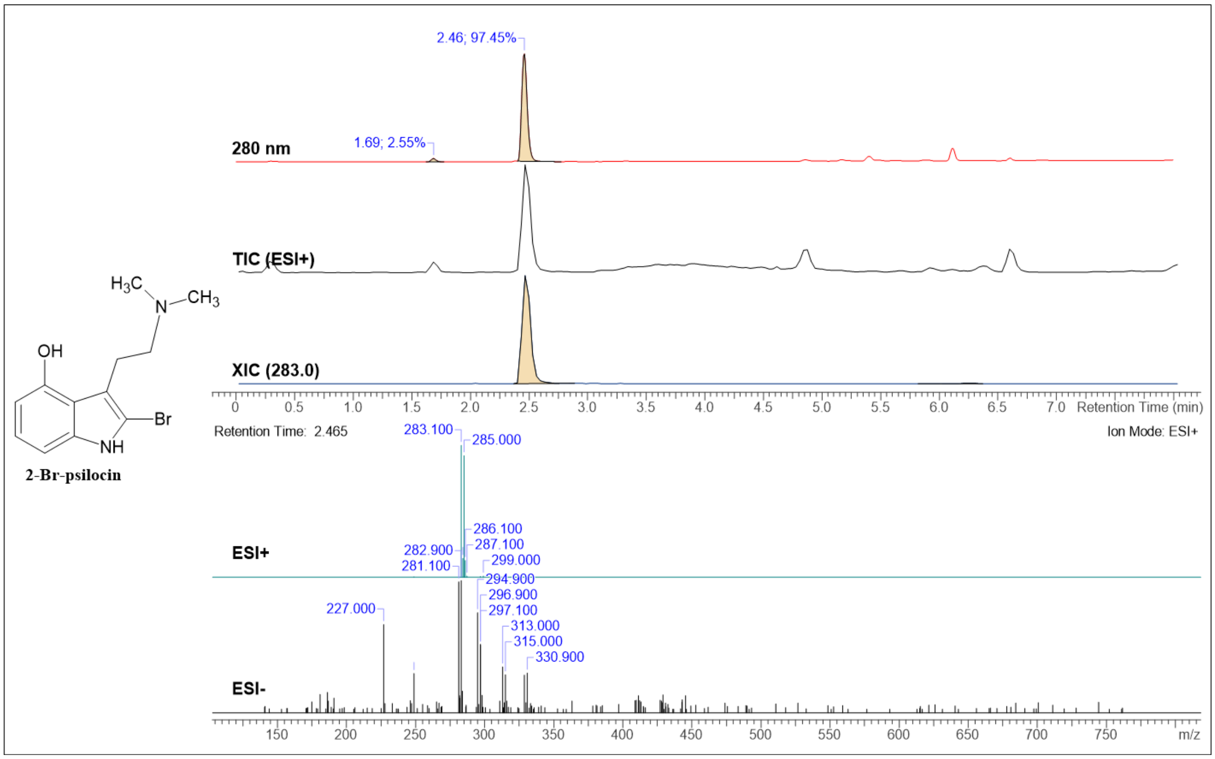


**Figure S31.** UV purity of 2-Br-psilocin

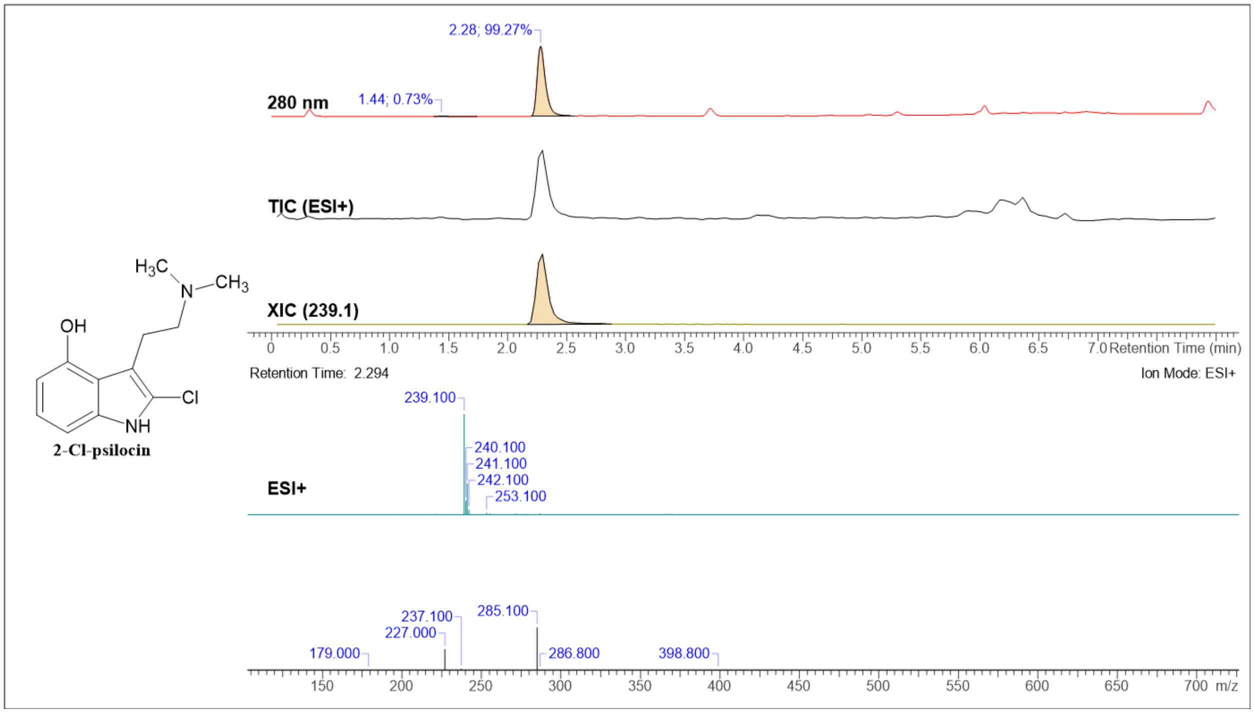


**Figure S32.** UV purity of 2-Cl-psilocin
